# How to design optimal brain stimulation to modulate phase-amplitude coupling?

**DOI:** 10.1101/2024.02.12.579897

**Authors:** Benoit Duchet, Rafal Bogacz

## Abstract

**Objective:** Phase-amplitude coupling (PAC), the coupling of the amplitude of a faster brain rhythm to the phase of a slower brain rhythm, plays a significant role in brain activity and has been implicated in various neurological disorders. For example, in Parkinson’s disease, PAC between the beta (13–30 Hz) and gamma (50–200 Hz) rhythms in the motor cortex is exaggerated, while in Alzheimer’s disease, PAC between the theta (4-8 Hz) and gamma rhythms is diminished. Modulating PAC (i.e. reducing or enhancing PAC) using brain stimulation could therefore open new therapeutic avenues. However, while it has been previously reported that phase-locked stimulation can increase PAC, it is unclear what the optimal stimulation strategy to modulate PAC might be. Here, we provide a theoretical framework to narrow down the experimental optimisation of stimulation aimed at modulating PAC, which would otherwise rely on trial and error.

**Approach:** We make analytical predictions using a Stuart-Landau model, and confirm these predictions in a more realistic model of coupled neural populations.

**Main results:** Our framework specifies the critical Fourier coefficients of the stimulation waveform which should be tuned to optimally modulate PAC. Depending on the characteristics of the amplitude response curve of the fast population, these components may include the slow frequency, the fast frequency, combinations of these, as well as their harmonics. We also show that the optimal balance of energy between these Fourier components depends on the relative strength of the endogenous slow and fast rhythms, and that the alignment of fast components with the fast rhythm should change throughout the slow cycle. Furthermore, we identify the conditions requiring to phase-lock stimulation to the fast and/or slow rhythms.

**Significance:** Together, our theoretical framework lays the foundation for guiding the development of innovative and more effective brain stimulation aimed at modulating PAC for therapeutic benefit.

## 1 Introduction

Phase-amplitude coupling (PAC), a type of cross-frequency coupling where the amplitude of faster brain oscillations is coupled to the phase of slower brain oscillations, is widespread across species and brain regions. Most notably, PAC was shown to be implicated in memory and learning, in particular through coupling of the amplitude of the gamma rhythm (50–200 Hz) to the phase of the theta rhythm (4-8 Hz) in the hippocampus [1, 2, 3, 4, 5, 6]. Beyond memory processes, PAC is for example modulated during movement and speech [7], visual attention [8], auditory processing [9], complex cognitive function [10], as well as during development [11].

PAC was also found to be abnormal in various neurological disorders – see [12] for a review. In Parkinson’s disease (PD), coupling between the beta phase (13–30 Hz) and gamma amplitude in the motor cortex is exaggerated compared to patients with dystonia and patients with epilepsy, both at rest and during movement [13]. Elevated PAC was reported in patients with PD off dopaminergic medication compared to patients on medication, as well as compared to humans without a movement disorder [14]. This increased PAC is reduced by deep brain stimulation (DBS) [15]. Similarly, alpha (8-12 Hz) gamma PAC is exaggerated in the sensorimotor cortex of patients with essential tremor [16]. As expected from its involvement in memory, theta-gamma PAC is impacted in Alzheimer’s disease (AD). Lower theta-gamma PAC than controls was found in AD rodent models [17, 18], with alterations appearing before significant accumulation of amyloid-beta in some animals [19]. In humans, theta-gamma PAC was lower in patients with mild cognitive impairment compared to healthy age-matched participants, lower still in patients with AD [20], and correlated with cognitive and memory performance [21, 20]. PAC was also reported to be elevated during epileptic seizures [22]. Furthermore, PAC was suggested as a biomarker for brain-computer interface-mediated motor recovery in chronic stroke [23], and for rehabilitation of speech discrimination in cochlear implant users [24].

Given changes in PAC from healthy levels in neurological disorders, in some cases correlated with symptoms or recovery, restoring healthy PAC levels is a promising target for neuromodulation therapies. However, how to stimulate to enhance or decrease PAC levels has received very little attention to date. A notable exception is the work by Salimpour and colleagues, which showed that phase-locking motor cortical electrical stimulation to the peak of the beta rhythm increased beta-gamma PAC in humans compared to baseline, and compared to stimulation phase-locked to the trough of the beta rhythm [25]. Similarly, phase-locking hippocampal transcranial ultrasound stimulation to the peak of the theta rhythm increased theta-gamma PAC in rats [26]. Nevertheless, it is unclear what the optimal stimulation strategy to enhance or decrease PAC might be. Here, we develop a theoretical framework to address this question using the analytically tractable Stuart-Landau (SL) model as well as a more biologically realistic neural mass model, the Wilson-Cowan (WC) model. While we focus on PAC-enhancing stimulation (which could be of interest for example in patients with AD), the same framework can be applied to stimulation aimed at reducing PAC. Our framework is directly applicable to neuromodulation modalities where the Fourier coefficients of the stimulation waveform can be tuned, such as transcranial alternating current stimulation (tACS). For modalities that can only generate square pulses (e.g. DBS), the optimal waveforms predicted from the theoretical framework can be approximated by pulsatile waveforms.

## 2 Results

We develop a theory of optimal PAC-enhancing stimulation using the SL model, which offers the possibility of analytical insights. In particular, we build on two key mechanisms contributing to increasing PAC, namely stimulation at the slow frequency, and stimulation with a modulated component at the fast frequency. We show that whether these mechanisms can be leveraged depends on characteristics of the fast population’s response to stimulation. We proceed to verify elements of the theory in a neural mass model, the WC model. We finish by considering practical questions, in particular the balance of the Fourier coefficients of the stimulation waveform as a function of the strength of the endogenous slow and fast rhythms, the necessity (or lack thereof) of phase-locking stimulation to the fast and/or slow rhythm, and how to approximate the optimal waveforms with pulsatile waveforms. Flowcharts that could guide experimentalists in designing optimal-PAC modulating stimulation are presented in figure 10.

### 2.1 Developing optimal PAC-enhancing stimulation in the Stuart-Landau model

The SL model is arguably the simplest phase-amplitude model used in neuroscience [27, 28, 29, 30], and is therefore ideally suited to obtain analytical insights on PAC-enhancing stimulation. The model represents the canonical form of a Hopf bifurcation, and can therefore operate in the fixed-point or limit-cycle regime. We are considering a SL population operating at the fast frequency of interest with order parameter 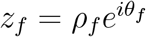 (oscillation amplitude *ρ*_*f*_ and oscillation phase *θ*_*f*_) evolving according to

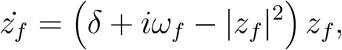

where *ω*_*f*_ = 2*πf*_*f*_ is the angular frequency of the fast population, and *δ* a bifurcation parameter. When *δ* > 0, the fast population is in the limit-cycle regime and generates intrinsic oscillations of amplitude converging to 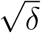. When *δ* ≤ 0, the fast population is in a quiescent state (fixed-point regime).

We assume that neither stimulation nor the fast rhythm significantly affects the slow rhythm contributing to PAC, and we model the slow rhythm as an input to the fast population. In the case of hippocampal theta-gamma PAC, the slow input can represent the theta input from pacemaker neurons in the medial septum for example, believed to be the main contributor to hippocampal theta [31, 32, 33]. We present the limitations of this approach in the Discussion. We further assume that the slow input of strength *k*_*s*_ and angular frequency *ω*_*s*_ = 2*πf*_*s*_ is coupled to the fast population through its mean-field (see figure 1), thereby affecting the amplitude (but not the phase) of the fast oscillations. As shown by Quin and colleagues, such a slow input can generate PAC [27] in the SL model. Indeed, since

**Figure 1:**
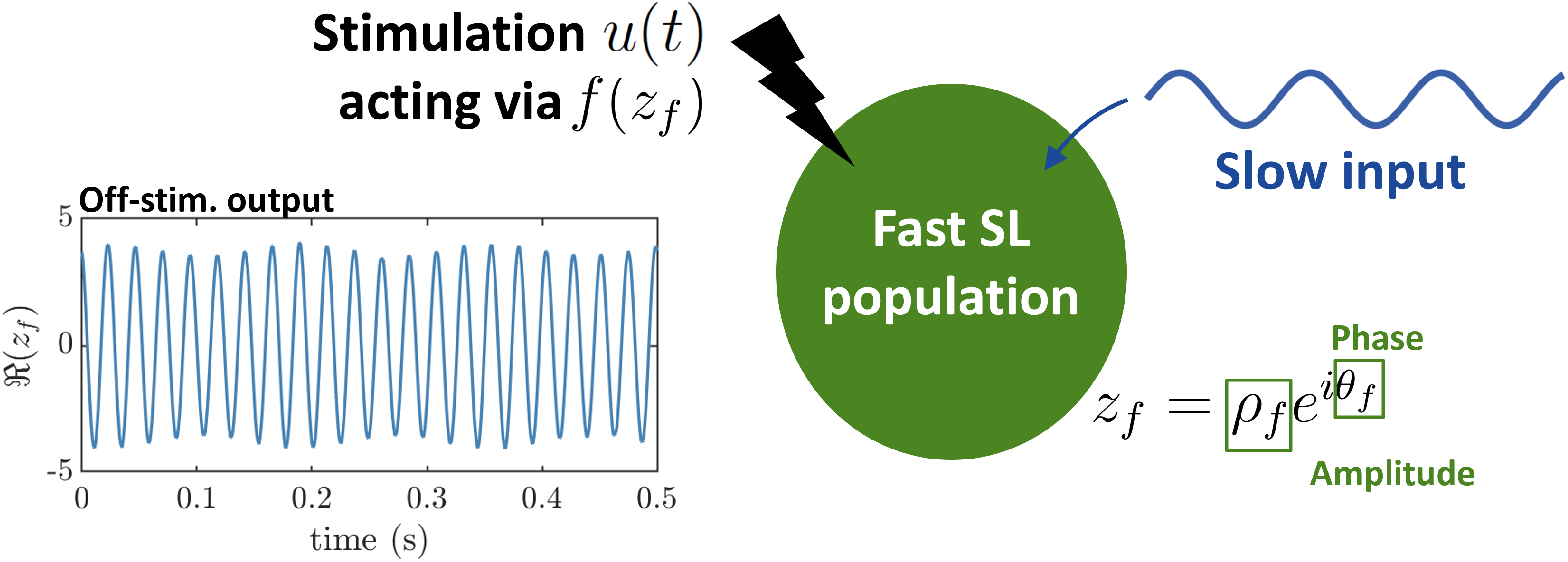
Sketch of the Stuart-Landau model with stimulation. Intrinsic PAC is generated by a slow input (shown in dark blue) interacting with a fast Stuart-Landau population (represented in green). An example of the output of the fast population (real part of the fast-population order parameter) displaying PAC in the absence of stimulation in shown in the left panel. The stimulation *u*(*t*) (in black) acts on the fast population via a stimulation coupling function *f*(*z*_*f*_), where *z*_*f*_ is the order parameter of the fast population with oscillation amplitude *ρ*_*f*_ and oscillation phase *θ*_*f*_.

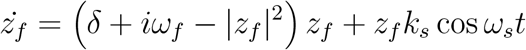

can be re-written as

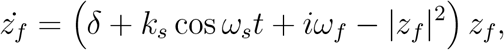

the parameter controlling the amplitude of the fast oscillations becomes *δ*_PAC_ = *δ* + *k*_*s*_ cos *ω*_*s*_*t*. This means that the amplitude of the fast oscillations is controlled by the phase of the slow input.

In the next sections, we will optimise the stimulation waveform to maximally increase PAC for a given stimulation energy budget. We will consider a stimulation input *u*(*t*) provided to the fast SL population receiving a slow input. The stimulation is provided to the population through the stimulation coupling function *f*(its connection with experimental measures is detailed below). The evolution of the order parameter of the fast SL population is given by

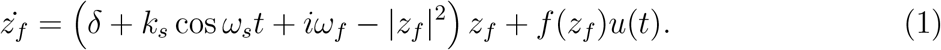

In what follows, we expand *u*(*t*) as a truncated Fourier Series

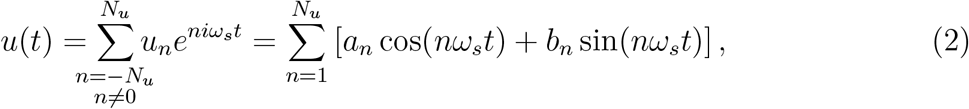

with complex coefficients *u*_*n*_, or equivalently real coefficients *a*_*n*_ and *b*_*n*_, and with truncation order *N*_*u*_. To enforce charge balance of the stimulus (a requirement for brain stimulation to avoid tissue damage), the zeroth-order coefficient is zero.

We will show in the next sections that the way stimulation is coupled to the fast population, and in particular the amplitude response curve (ARC) of the fast population, determines which Fourier coefficients contribute to modulating PAC. The stimulation coupling function is directly related to the ARC and the phase response curve (PRC) of the fast population. Here, the ARC describes the instantaneous change in amplitude of the collective activity of a neural population (e.g. measured in the local field potential [34, 35]) due to stimulation. The ARC is a function of the state of the neural population, e.g. the phase and/or amplitude of the collective oscillation when stimulation is received. Similarly, we take the PRC to refer to the instantaneous change in phase of the collective oscillation due to stimulation, as a function of the state of the neural population. Note that in some studies, the PRC refers instead to the response of individual neurons (e.g. [36, 37, 38]). This is in contrast with this study, where we consider changes on the population level. Given these definitions, we have 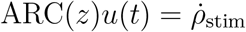 and 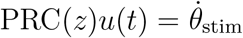, where 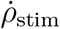 and 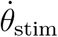 are the instantaneous changes in amplitude and phase of the neural population due to stimulation, respectively, and *z* is the order parameter of the SL model. Using the product rule on the definition of the order parameter, we have 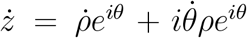. Without loss of generality, the instantaneous change in *z* due to stimulation at time *t* can therefore be written as *ż*_stim_ = [ARC(*z*)*e*^*iθ*^ + *i*PRC(*z*)*z*] *u*(*t*). Since *ż*_stim_ = *f*(*z*_*f*_)*u*(*t*), we can identify the stimulation coupling function as

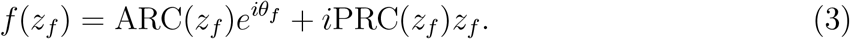

We will show that systems with different ARCs require different stimulation waveforms to optimally modulate PAC. Examples of ARCs with their corresponding optimal PAC-enhancing waveforms are given for the SL model in figure 2, and later for the WC model in figure 9.

**Figure 2:**
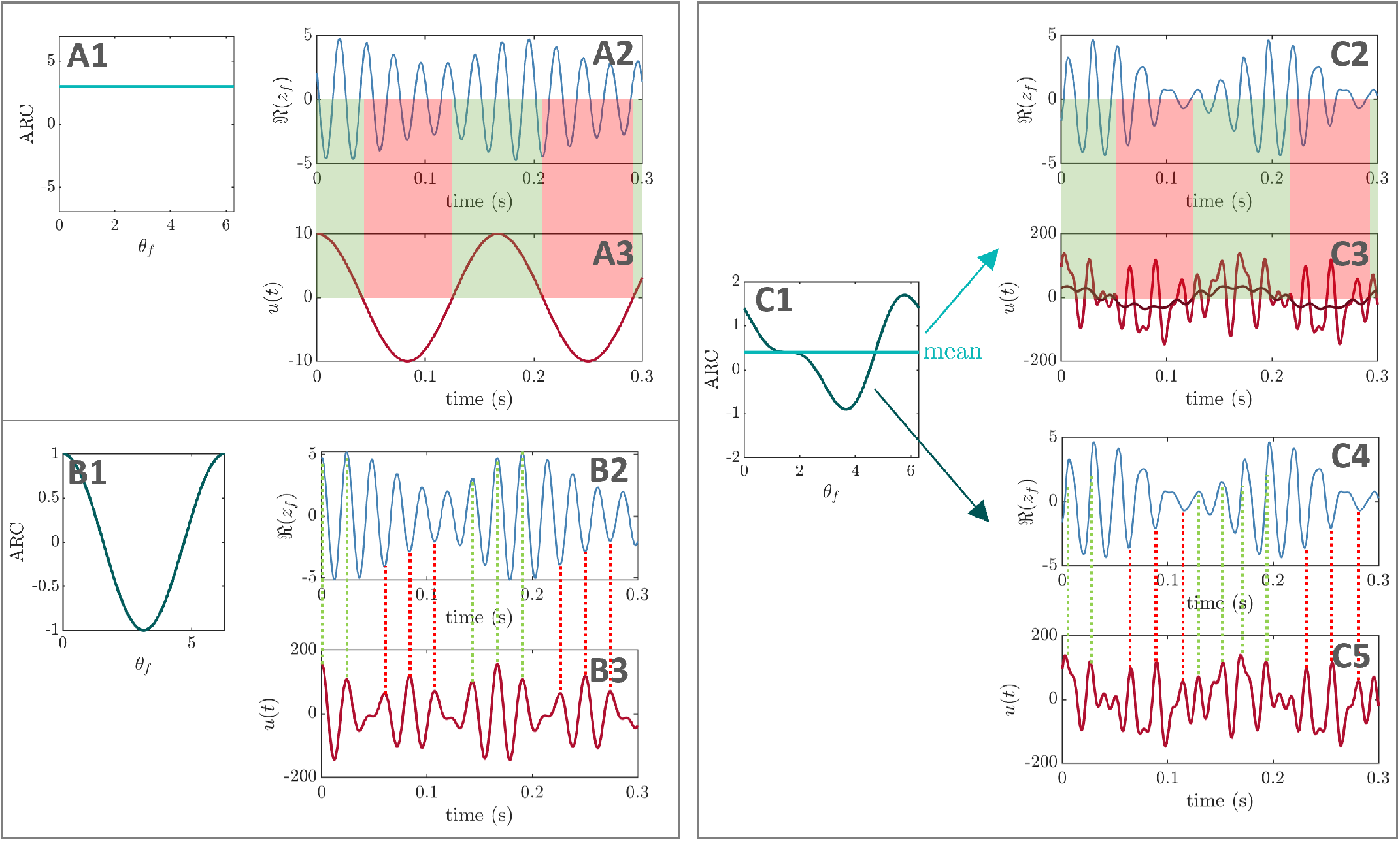
PAC-enhancing mechanisms in the Stuart-Landau model depend on the amplitude response of the fast population. Panel A corresponds to foundational case one, where the amplitude response of the fast population does not depend on its phase (A1, shown for *ρ*_*f*_ = 3). Sinusoidal stimulation (A3) therefore enhances the amplitude of the fast population (A2) when the stimulation waveform is positive (green highlights), and suppresses the amplitude of the fast population when it is negative (red highlights). Panel B corresponds to foundational case two, where the amplitude response of the fast population depends on its phase but has zero mean (B1). Where the fast-oscillation amplitude (B2) should be increased, the optimal stimulation waveform (B3, taken from figure 4C) has fast-frequency components aligned with the peak of the fast rhythm (green dashes). Conversely, where the fast-oscillation amplitude should be decreased, the optimal stimulation waveform has fast-frequency components anti-phase-aligned with the peak of the fast rhythm. Panel C corresponds to a general case where the amplitude response of the fast population does depend on its phase and has a non-zero mean (highlighted in light green in C1). The optimal stimulation waveform (taken from figure 5C) combines mechanisms of PAC-enhancement from panels A and B, as shown in panels C2-C5. The dark red line in C3 represents a moving average of the stimulation waveform (sliding window corresponding approximately to two fast-population cycles.)

Before dealing with arbitrary stimulation coupling functions (i.e. arbitrary ARCs and PRCs), we consider two foundational cases to uncover the two mechanisms of action contributing to PAC enhancement in the general case. We will show below that in the first foundational case where the amplitude response of the fast population does not depend on its phase, the optimal stimulation is at the frequency of the slow rhythm (figure 2A). In the second foundational case where the amplitude response of the fast population depends on its phase but has zero mean, the optimal stimulation is at the fast frequency, with fast frequency components modulated by the slow frequency (figure 2B). The general case combines both strategies (figure 2C). In each case, we derive theoretical results and test them using numerical optimisation.

#### 2.1.1 Foundational case one: stimulation is coupled through the mean-field

In this first foundational case where stimulation is coupled to the fast population through its mean field, i.e. *f*(*z*_*f*_) = *z*_*f*_, the optimal PAC-enhancing waveform can be approximated analytically. Using equation (3), we note that this type of stimulation coupling through the mean-field of the fast population is equivalent to ARC = *ρ*_*f*_ and PRC = 0. The ARC is therefore positive, with no dependence on the fast population phase. We have

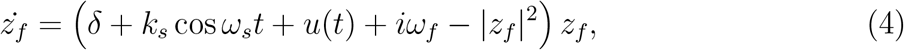

and the parameter controlling the amplitude of the fast oscillations is therefore given by *δ*_PAC_ = *δ* + *k*_*s*_ cos *ω*_*s*_*t* + *u*(*t*).

##### Approximate analytical solution

To analytically quantify PAC in the system described by equation (4), we modify a PAC measure called the mean vector length (MVL) [39, 40]. The MVL is recommended for high signal-to-noise ratio [40], which is the case in this modelling approach. We define our modified MVL measure as

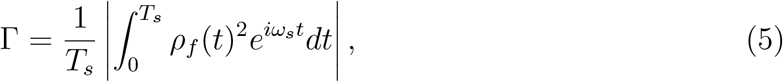

where *T*_*s*_ = 2*π*/*ω*_*s*_ is the period of the slow rhythm, *ρ*_*f*_ is the amplitude of the fast oscillations, and *ω*_*s*_*t* is the phase of the slow input. Our PAC measure Γ is the direct translation of the MVL (as defined in [40]) to continuous time over one period, with the exception that the amplitude of the fast oscillations is replaced by *ρ*_*f*_ (*t*)^2^ (i.e. power) for analytical convenience (as will become apparent below). As in the original definition, when the amplitude of the fast oscillation is high for a consistent range of phases of the slow oscillation, the magnitude of the resulting vector will be large and PAC will be detected. Assuming the square of the envelope of the fast oscillations can be expressed as a Fourier series, our PAC measure Γ can also be interpreted as the modulus of the complex Fourier coefficient of 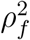 at the slow frequency. The modified PAC measure Γ therefore captures the strength of the modulation of *ρ*_*f*_ at the slow frequency *ω*_*s*_.

Assuming relaxation to the limit cycle arising from equation (4) is fast enough, we have 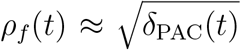 for *δ*_PAC_(*t*) > 0, which allows us to compute Γ (this assumption can be relaxed using a semi-analytical approach described in Supplementary Material section A). We therefore have

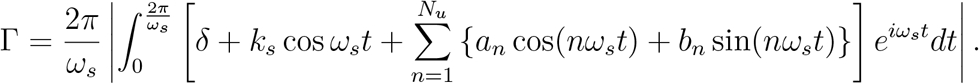

The only non-zero terms correspond to products of sines or cosines at the same frequency, which yields

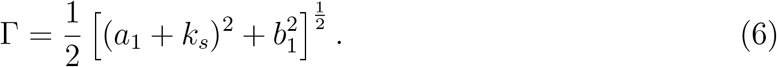

Remarkably, the PAC measure Γ only depends on the first harmonic of the stimulation. For a given stimulation energy Ξ, we can find the values of *a*_1_ and *b*_1_ that maximise Γ using the method of Lagrange multipliers. The energy constraint is 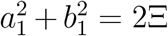, and the corresponding Lagrangian function reads

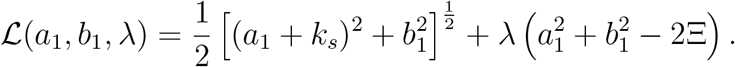

Setting its derivatives with respect to *a*_1_, *b*_1_, and *λ* to 0 leads to 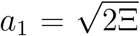 and *b*_1_ = 0, i.e.

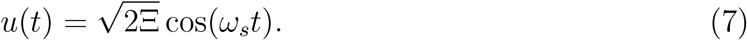

The optimal stimulation strategy therefore consists in providing sinusoidal stimulation at the slow frequency, with its peak aligned to the peak of the slow rhythm. This optimal waveform makes intuitive sense since the ARC of the fast population is positive and does not depend on the phase of the fast population. Sinusoidal stimulation therefore enhances the amplitude of the fast population when the stimulation waveform is positive, and suppresses the amplitude of the fast population when the stimulation waveform is negative as illustrated in figure 2A.

##### Verification using numerical optimisation

We verify using numerical optimisation that the waveform given by equation (7) closely approximates the optimal PAC-enhancing waveform. To this end, we optimise the Fourier coefficients of *u*(*t*) up to *N*_*u*_ = 5 to maximise the MVL (obtained as equation (12), see section 4.1 in the Appendix) while constraining the energy of *u*(*t*) to Ξ. Methodological details of the optimisation process can be found in section 4.2.

The best-ranked stimulation waveform obtained from numerical optimisation is a close approximation of the sinusoidal waveform given by equation (7), as shown in figure 3 (see panel I, and compare panels D and H). These results are consistent across the top-50 optimisations (figure 3E). Note that similar results are obtained when maximising the MVL (figure 3) or a discrete approximation of Γ, i.e. measures based on *ρ*_*f*_ or 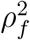, respectively. We also perturb each Fourier coefficient in turn by adding the perturbation 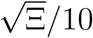, where Ξ is the waveform energy before perturbation. This analysis confirms the dominant impact of the first Fourier component of the stimulation waveform on PAC. This is true both when perturbing PAC-enhancing waveforms (figure 3J) and random waveforms (figure 3K). Methodological details for this analysis can be found in section 4.3.

**Figure 3:**
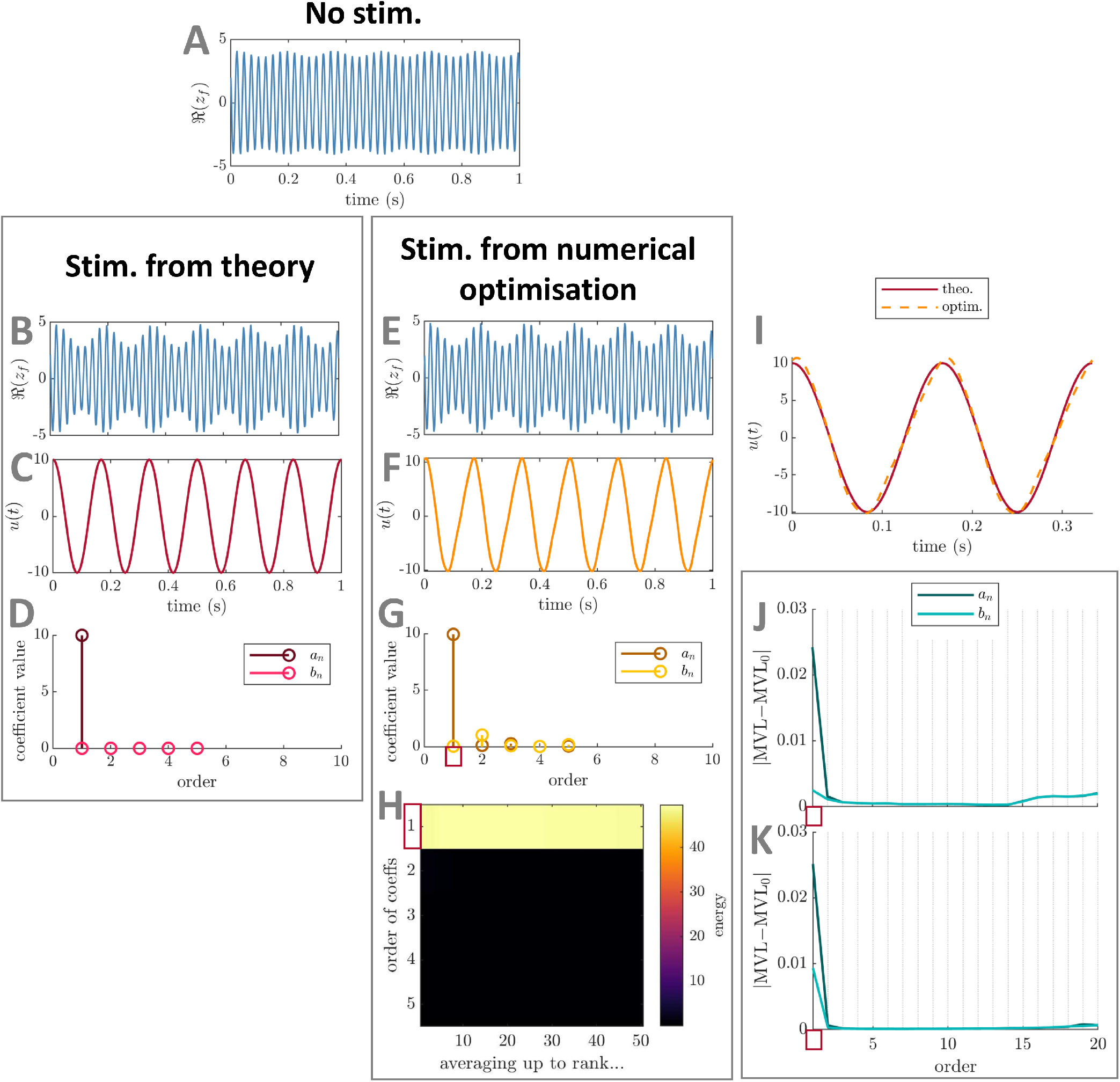
Comparison between optimal PAC-enhancing waveforms predicted by theory and by numerical optimisation – foundational case one (mean-field coupled stimulation) in the Stuart-Landau model. The model output in the absence of stimulation is shown in panel A. The model output when receiving optimal PAC-enhancing stimulation is shown in panels B (stimulation waveform predicted by theory) and F (stimulation waveform obtained though numerical optimisation). The corresponding optimal PAC-enhancing stimulation waveforms are shown in panels C and G, respectively, and are overlaid for comparison in panel I. Their Fourier coefficients are shown in panels D and H, respectively. Panel E represents the energy of PAC-enhancing waveforms obtained from numerical optimisation for all Fourier coefficient orders (vertical axis), when averaging the *x*-best optimisation results (*x* being the horizontal axis value). The absolute change in MVL when increasing the energy of a given stimulation Fourier coefficient is provided in panels J (when starting from PAC-enhancing waveforms obtained from the numerical optimisation process), and K (when starting from random waveforms). Error bars (too small to see here) represent the standard error of the mean. The Fourier coefficients predicted to be key contributors to PAC levels by theory are highlighted by red rectangles in panels H, J, and K. MVL for the stimulation waveform predicted by theory is 0.335, MVL for the stimulation waveform obtained though numerical optimisation is 0.337, MVL in the absence of stimulation is 0.079 (Δ*f*_*f*_ = 10 Hz). In all cases, waveform energy is fixed at Ξ = 50. The parameters of the SL model used are *δ* = 15, *k*_*s*_ = 3, *f*_*f*_ = 40 Hz, and *f*_*s*_ = 6 Hz.

#### 2.1.2 Foundational case two: stimulation acts through a direct coupling

We next consider a second foundational case where stimulation acts through *f*(*z*_*f*_) = 1, which we call “direct” coupling. With the activity of the fast population modelled as ℜ(*z*_*f*_) (where ℜ(.) denotes the real part), this case represents stimulation directly increasing the firing rate of the fast population. Using equation 3, *f*(*z*_*f*_) = 1 corresponds to ARC = cos *θ*_*f*_ and PRC = − sin *θ*_*f*_ /*ρ*_*f*_ (for *ρ*_*f*_ > 0). From equation (1) with direct coupling, the time evolutions of *ρ*_*f*_ and *θ*_*f*_ are given by

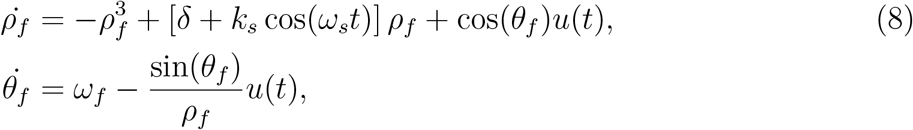

for *ρ*_*f*_ > 0.

##### Theoretical predictions

While an analytical solution is out of reach, we can determine which Fourier coefficients of the stimulation waveform should be considered to enhance PAC. To study the effect of stimulation on *ρ*_*f*_, we approximate *θ*_*f*_ by

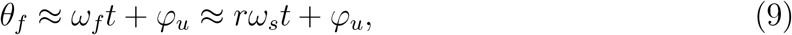

where *r* is the closest integer to *ω*_*f*_ /*ω*_*s*_, and *φ*_*u*_ is a constant phase (the subscript *u* denotes a potential dependence on the stimulation waveform). This approximation is justified if the stimulation amplitude is small (in which case deviations of 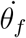 from *ω*_*f*_ would be small), and *ω*_*f*_ ≫ *ω*_*s*_, which is often the case in the brain (e.g theta-gamma coupling). The approximation *θ*_*f*_ *≈ rω*_*s*_*t* + *φ*_*u*_ is also justified for larger stimulation amplitudes leading to *r* : 1 entrainment. We therefore have approximately

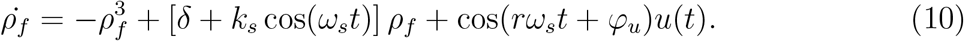

Although an exact solution has recently been found for this type of differential equations (Abel’s equation of the first kind) [41], it cannot be expressed directly as a function of the Fourier coefficients of the stimulation. Instead, we can gain insight by noting that in the steady-state, solutions with PAC will be periodic with period 2*π*/*ω*_*s*_, and thus can be approximated as truncated Fourier series. Since most of the PAC strength is captured by the first harmonic of *ρ*_*f*_, we only consider its zeroth and first order components parametrised by *ρ*_0_, *ρ*_1_, and *θ*_1_ such that *ρ*_*f*_ (*t*) = *ρ*_0_ + 2*ρ*_1_ cos (*ω*_*s*_*t* + *θ*_1_). We show in section 4.4 in the Appendix that equation (10) translates to three equations in *ρ*_0_, *ρ*_1_, and *θ*_1_ (equations (15)-(17)).

While these equations cannot be solved easily, they demonstrate that the zeroth and first harmonic of *ρ*(which will determine PAC strength) only depend on the Fourier coefficients of the stimulation of order *r, r* − 1, *r* + 1 (recall that *r* is the closest integer to *ω*_*f*_ /*ω*_*s*_). Since the base frequency of *u*(*t*) is *ω*_*s*_, these Fourier coefficients correspond to frequencies *f*_*f*_, and *f*_*f*_ ± *f*_*s*_. The small number of Fourier coefficients involved significantly simplifies the task of finding an optimal stimulation to increase PAC, and also highlights that how stimulation couples to the neural circuit of interest has a large influence on the optimal stimulation. The optimal stimulation here is very different from the mean-coupling case where stimulation at *ω*_*s*_ is optimal, i.e. only *u*_1_ is non-zero. Note the theory predicts that no Fourier coefficient other than the coefficients of order *r, r* − 1, *r* + 1 plays a key role in enhancing PAC, but the coefficients of order *r, r* − 1, *r* + 1 need not all have a significant impact on PAC.

##### Verification using numerical optimisation

We verify using numerical optimisation that the key stimulation waveform Fourier coefficients to optimally enhance PAC (for direct coupling) are limited to (possibly a subset of) coefficients of order *r, r* − 1, and *r* + 1 as predicted by theory. To this end, we optimise either the Fourier coefficients of *u*(*t*) predicted by theory, or all Fourier coefficients up to *N*_*u*_ = 10. In both cases, the objective is to maximise the MVL (obtained as equation (12), see section 4.1 in the Appendix) while constraining the energy of *u*(*t*) to Ξ. As before, methodological details of the optimisation process can be found in section 4.2.

The results of numerical optimisation support theoretical predictions. The best-ranked stimulation waveforms obtained from both numerical optimisations show close similarities (figure 4J), and the resulting PAC levels are similar (with a slight advantage when optimising Fourier coefficients predicted by theory). Additionally, the best-ranked waveform obtained when optimising all coefficients concentrates its energy in the Fourier coefficients predicted by theory (highlighted by a red rectangle in figure 4H). This is consistent across the top-50 local optimisations (figure 4I). Moreover, perturbing individual Fourier coefficients in turn confirms the dominant impact of the stimulation waveform Fourier components of order *r, r* − 1, and *r* + 1. This is true both when perturbing PAC-enhancing waveforms (figure 4K) and random waveforms (figure 4L). As before, the perturbation size is 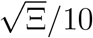 and methodological details for this analysis can be found in section 4.3. These numerical results were obtained for *ω*_*f*_ /*ω*_*s*_ = 7, and we also verify that our theoretical predictions hold true for non-integer values of *ω*_*f*_ /*ω*_*s*_ (see figure S.2 in Supplementary Material).

**Figure 4:**
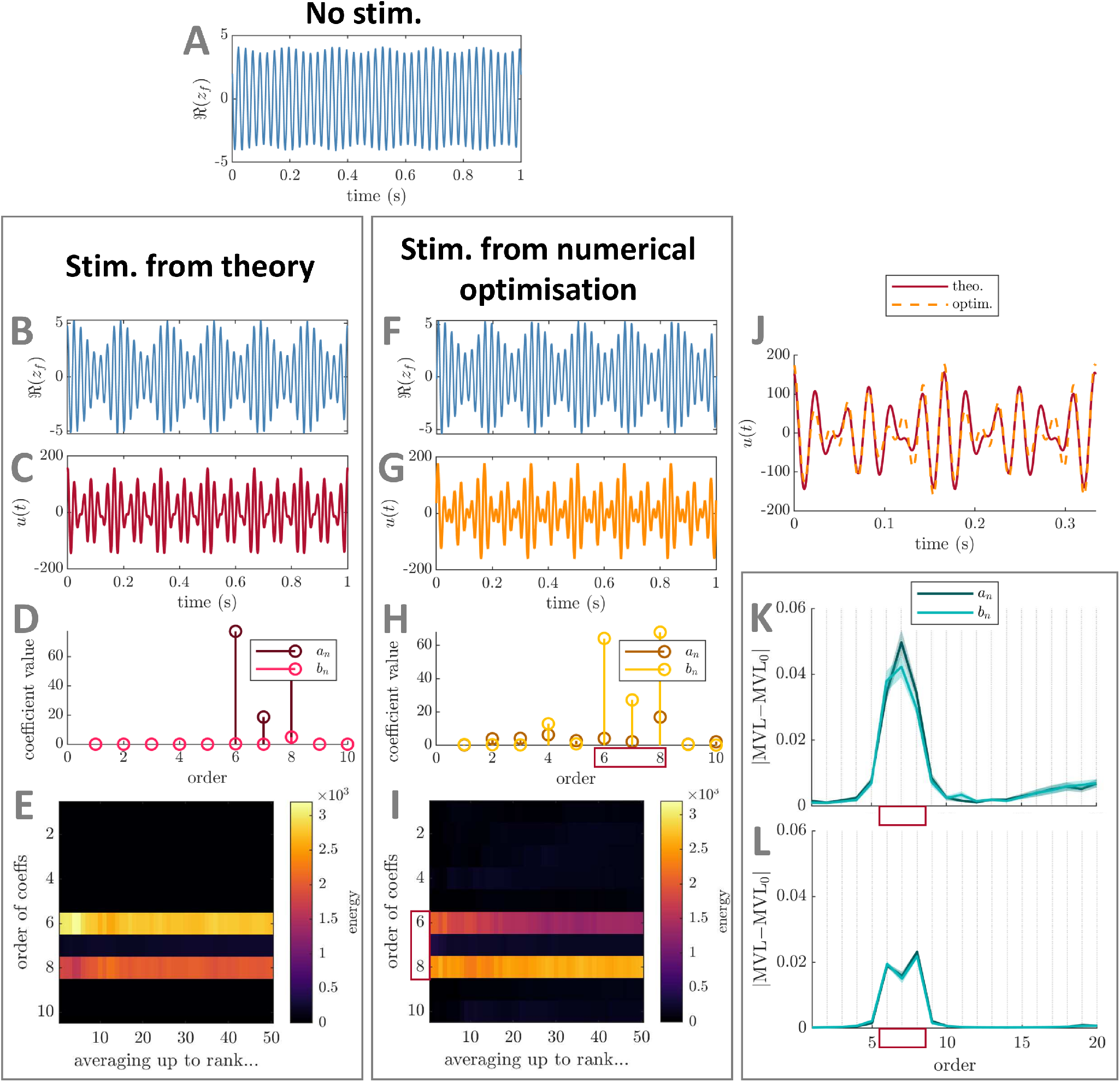
Comparison between best PAC-enhancing waveforms predicted by theory and by numerical optimisation – foundational case two (direct stimulation coupling) in the Stuart-Landau model. The model output in the absence of stimulation is shown in panel A. The model output when receiving PAC-enhancing stimulation is shown in panels B (best stimulation waveform obtained when optimising only Fourier coefficients predicted by theory) and F (best stimulation waveform obtained when optimising all Fourier coefficients). The corresponding best PAC-enhancing stimulation waveforms are shown in panels C and G, respectively, and are overlaid for comparison in panel J (aligned to maximise their cross-correlation). Their Fourier coefficients are shown in panels D and H, respectively. The energy of PAC-enhancing waveforms obtained from numerical optimisation for all Fourier coefficient orders (vertical axis) when averaging the *x*-best optimisation results (*x* being the horizontal axis value) is represented in panels E (only Fourier coefficients predicted by theory were optimised) and I (all Fourier coefficients were optimised). The absolute change in MVL when increasing the energy of a given stimulation Fourier coefficient is provided in panels K (when starting from PAC-enhancing waveforms obtained from the numerical optimisation process with all coefficients optimised), and L (when starting from random waveforms). Error bars represent the standard error of the mean. The Fourier coefficients predicted to be (potential) key contributors to PAC levels by theory are highlighted by red rectangles in panels H, I, K, and L. MVL for the stimulation waveform with only coefficients predicted by theory optimised is 0.563, MVL for the stimulation waveform with all coefficients optimised is 0.533, MVL in the absence of stimulation is 0.082 (Δ*f*_*f*_ = 20 Hz). In all cases, waveform energy is fixed at Ξ = 5000. The parameters of the SL model used are *δ* = 15, *k*_*s*_ = 3, *f*_*f*_ = 42 Hz, and *f*_*s*_ = 6 Hz (*r* = 7).

This second foundational example illustrates a key mechanism of action of PAC-enhancing stimulation when the amplitude response of the fast population depends on its phase and has zero mean. In this case, modulating the amplitude of the fast population necessarily requires fast-frequency oscillations in the stimulation waveform. The optimal waveform obtained from numerical optimisation in figure 2B demonstrate that, if the ARC is maximum and positive at *θ*_*f*_ = 0, the fast-frequency oscillations in the stimulation waveform should phase-align with the oscillations of the fast population in the part of the slow-frequency cycle where the fast-oscillation amplitude should be increased (see green dashed lines in figure 2B). Conversely, if the ARC is minimum and negative at *θ*_*f*_ = *π* for example, the fast-frequency oscillations in the stimulation waveform should anti-phase-align with the oscillations of the fast population in the part of the slow-frequency cycle where the fast-oscillation amplitude should be decreased (see red-dashed lines in figure 2B). While the theoretical analysis presented above does not describe how the fast-frequency oscillations in the optimal waveform should be arranged, the definition of the ARC requires the phase alignment between the stimulation’s fast components and the oscillations of the fast population to change throughout the slow cycle. Indeed, if this weren’t the case, the effect of stimulation on the amplitude of the fast population would be the same throughout the slow cycle as the peaks of stimulation at the fast frequency would consistently occur around the same phase of the fast population’s oscillation. As detailed above, the optimal waveform obtained from numerical optimisation confirms that the ARC of the fast population dictates how phase alignment between the stimulation’s fast components and the fast rhythm should change throughout the slow cycle to optimally enhance PAC. Specifically, stimulation energy should be concentrated close to the phase corresponding to maximum amplification in the ARC where the fast population’s oscillations should be strengthened, and close to phase corresponding to maximum suppression in the ARC where the fast population’s oscillations should be weakened.

#### 2.1.3 Stimulation acts through a general coupling

In this section, we consider that stimulation acts through a general coupling. In general, the ARC of the fast population will combine features of the foundational cases investigated in the previous sections, i.e. non-zero mean and phase dependence. We assume 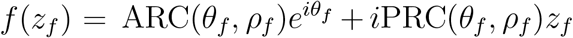 where ARC(*θ*_*f*_, *ρ*_*f*_) is a separable function of *θ*_*f*_ and *ρ*_*f*_. This is for instance the case for the mean-field of a population of neurons represented by phase oscillators [42]. Since the ARC of the fast population is also a periodic function of *θ*_*f*_, it can be approximated as 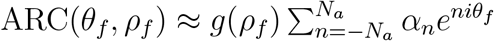.

##### Theoretical predictions

As before, we aim to determine which Fourier coefficients of the stimulation waveform should be considered to enhance PAC. We show in section 4.5 in the Appendix that the zeroth and first order components of *ρ*_*f*_ (parametrised by *ρ*_0_, *ρ*_1_, and *θ*_1_ as previously) must satisfy equations (21)-(23), where *g*(*ρ*_*f*_) was approximated by a truncated Fourier series 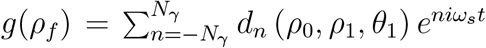 with truncation order *N*_*γ*_ (note that each *d*_*n*_ depends on the Fourier coefficients of *ρ*_*f*_).

While these equations cannot be solved analytically, they demonstrate that if the ARC of the fast population has *N*_*a*_ Fourier coefficients, the zeroth and first harmonic of *ρ*_*f*_ (and therefore PAC strength) only depend on (possibly a subset of) the Fourier coefficients of the stimulation of order

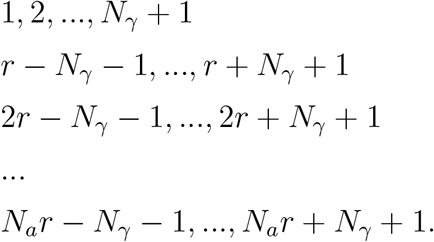

To summarize, a *k*^th^ harmonic in the ARC of the fast population leads to coefficients of frequency *kf*_*f*_ and *kf*_*f*_ ± *f*_*s*_ in the optimal stimulation waveform for *k* > 0, while a non-zero mean in the ARC results in the addition of the slow frequency *f*_*s*_. Significant dependence of the ARC on *ρ*_*f*_ requires additional neighbouring frequencies in steps of *f*_*s*_ until ± (*N*_*γ*_ + 1)*f*_*s*_ from *kf*_*f*_, and until (*N*_*γ*_ + 1)*f*_*s*_ from *f*_*s*_. In particular, if the ARC of the fast population has a dominant first harmonic and does not depend strongly on *ρ*_*f*_, it will be sufficient to optimise the Fourier coefficients of the stimulation waveform corresponding to *f*_*s*_, *f*_*f*_, and *f*_*f*_ ± *f*_*s*_ to determine the optimal stimulation strategy. This corresponds to the combination of the two foundational cases presented earlier.

##### Verification using numerical optimisation

To verify these predictions using numerical optimisation, we consider response curves of the fast population with a non-zero mean and two harmonics given by

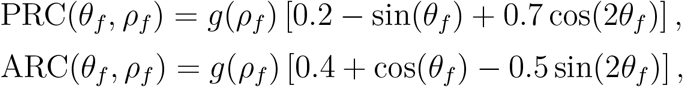

for *g*(*ρ*_*f*_) = 1 and *g*(*ρ*_*f*_) = 1/(*ρ*_*f*_ + 0.01). Thus, we verify in the former case that the key stimulation waveform Fourier coefficients contributing to enhancing PAC are limited to (possibly a subset of) coefficients of order 1, *r, r* − 1, *r* + 1, as well as 2*r* − 1, 2*r*, and 2*r* + 1 as predicted (second harmonic in ARC). In the latter case, the set of predicted potential dependences expand to also include coefficients of order 2, *r* − 2, *r* + 2 as well as 2*r* − 2, and 2*r* + 2 (we take *N*_*γ*_ = 1 since *g*(*ρ*_*f*_) is well described off-stimulation by one harmonic as shown in figure S.3A in Supplementary Material). As before, we optimise either the Fourier coefficients of *u*(*t*) predicted by theory, or all Fourier coefficients up to *N*_*u*_ = 20 to maximise the MVL while constraining the energy of *u*(*t*) to Ξ (see sections 4.1 and 4.2 for methodological details).

In both cases, the results of numerical optimisation support theoretical predictions (see figure 5 for *g*(*ρ*_*f*_) = 1 and figure S.4 for *g*(*ρ*_*f*_) = 1/(*ρ*_*f*_ +0.01) in Supplementary Material). The best-ranked stimulation waveforms obtained from optimising coefficients predicted by theory and all coefficients show similarities (compare panels C and G), and the resulting PAC levels are similar (in both cases with a slight advantage when optimising Fourier coefficients predicted by theory). Additionally, the best-rank waveforms obtained when optimising all coefficients concentrate their energy in a subset of the Fourier coefficients predicted by theory (highlighted by red rectangles in panels H in both figures). In both cases, this is consistent across the top-50 local optimisations (panels I in both figures), and confirmed by perturbation analysis (perturbation of PAC-enhancing waveforms in panels K and of random waveforms in panels L, see section 4.3 for methodological details, perturbation of size 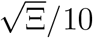 as before). Comparing figure 5H and figure S.4H, significant energy is introduced in Fourier coefficients of order 2 and 2*r* − 1 when the ARC of the fast population depends on *ρ*_*f*_ as opposed to when it doesn’t. However the Fourier components with the largest impact on PAC are the same in both cases (see panels K and L). In figure S.4H, the energy of *b*_3_ is not negligible, indicating that *g*(*ρ*_*f*_) = 1/(*ρ*_*f*_ + 0.01) is best described by two harmonics when stimulation is on (see figure S.3B2 in Supplementary Material).

**Figure 5:**
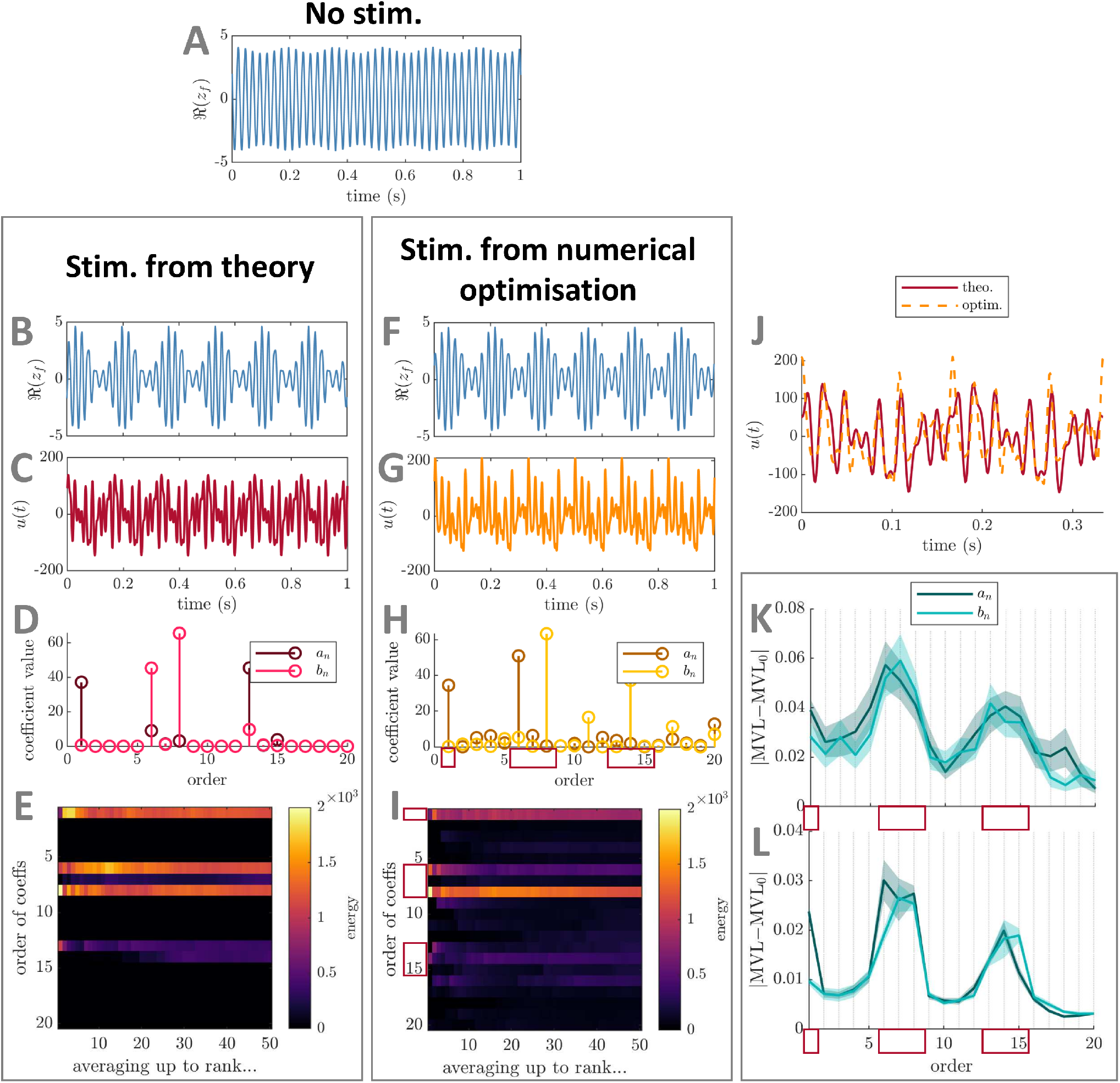
Comparison between best PAC-enhancing waveforms predicted by theory and by numerical optimisation – example for the general stimulation coupling case (no *ρ*_*f*_ dependence) in the Stuart-Landau model. The model output in the absence of stimulation is shown in panel A. The model output when receiving PAC-enhancing stimulation is shown in panels B (best stimulation waveform obtained when optimising only Fourier coefficients predicted by theory) and F (best stimulation waveform obtained when optimising all Fourier coefficients). The corresponding best PAC-enhancing stimulation waveforms are shown in panels C and G, respectively, and are overlaid for comparison in panel J (aligned to maximise their cross-correlation). Their Fourier coefficients (absolute values) are shown in panels D and H, respectively. The energy of PAC-enhancing waveforms obtained from numerical optimisation for all Fourier coefficient orders (vertical axis) when averaging the *x*-best optimisation results (*x* being the horizontal axis value) is represented in panels E (only Fourier coefficients predicted by theory were optimised) and I (all Fourier coefficients were optimised). The absolute change in MVL when increasing the energy of a given stimulation Fourier coefficient is provided in panels K (when starting from PAC-enhancing waveforms obtained from the numerical optimisation process with all coefficients optimised), and L (when starting from random waveforms). Error bars represent the standard error of the mean. The Fourier coefficients predicted to be (potential) key contributors to PAC levels by theory are highlighted by red rectangles in panels H, I, K, and L. MVL for the stimulation waveform with only coefficients predicted by theory optimised is 0.679, MVL for the stimulation waveform with all coefficients optimised is 0.643, MVL in the absence of stimulation is 0.082 (Δ*f*_*f*_ = 20 Hz). In all cases, waveform energy is fixed at Ξ = 5000. The parameters of the Stuart-Landau model used are *δ* = 15, *k*_*s*_ = 3, *f*_*f*_ = 42 Hz, and *f*_*s*_ = 6 Hz (*r* = 7). Stimulation is acting through PRC(*θ*) = 0.2 − sin(*θ*) + 0.7 cos(2*θ*) and ARC(*θ*) = 0.4 + cos(*θ*) − 0.5 sin(2*θ*).

The optimal waveforms obtained from numerical optimisation in figure 5C and figure S.4C combine the two PAC-enhancing mechanisms presented in the two foundational cases (see figure 2A-B). Because the mean amplitude response of the fast population across phases is non-zero (figure 2C1), the optimal stimulation waveform has a slow-frequency component that directly participates in expanding and shrinking the fast-frequency oscillations to produce PAC (figure 2C2-3 and figure S.5C-D in Supplementary Material). Moreover, because the amplitude response of the fast population strongly depends on its phase (figure 2C1), modulating the amplitude of the fast population requires fast-frequency components in the stimulation waveform whose alignment with the fast rhythm is modulated throughout the slow cycle (figure 2C4-5 and figure S.5E-F in Supplementary Material).

Using the SL model, we have identified how characteristics of the fast population’s response to stimulation dictate which frequencies should be included in the optimal stimulation waveform, and how to align potential fast stimulation components throughout the slow cycle according to the fast population’s ARC. We next investigate whether these predictions carry over to a neural mass model generating PAC.

### 2.2 Testing optimal PAC-enhancing stimulation in the Wilson-Cowan model

To test predictions obtained with the SL model, we use a more realistic neural mass model representing interacting neural populations, the Wilson-Cowan model [43]. This model was proposed as a canonical circuit to generate theta-gamma PAC in the presence of a theta input [44]. The biologically-inspired WC model describes the interactions of an excitatory (E) and an inhibitory (I) populations (see figure 6). The model is presented in details in section 4.6 in the Appendix.

**Figure 6:**
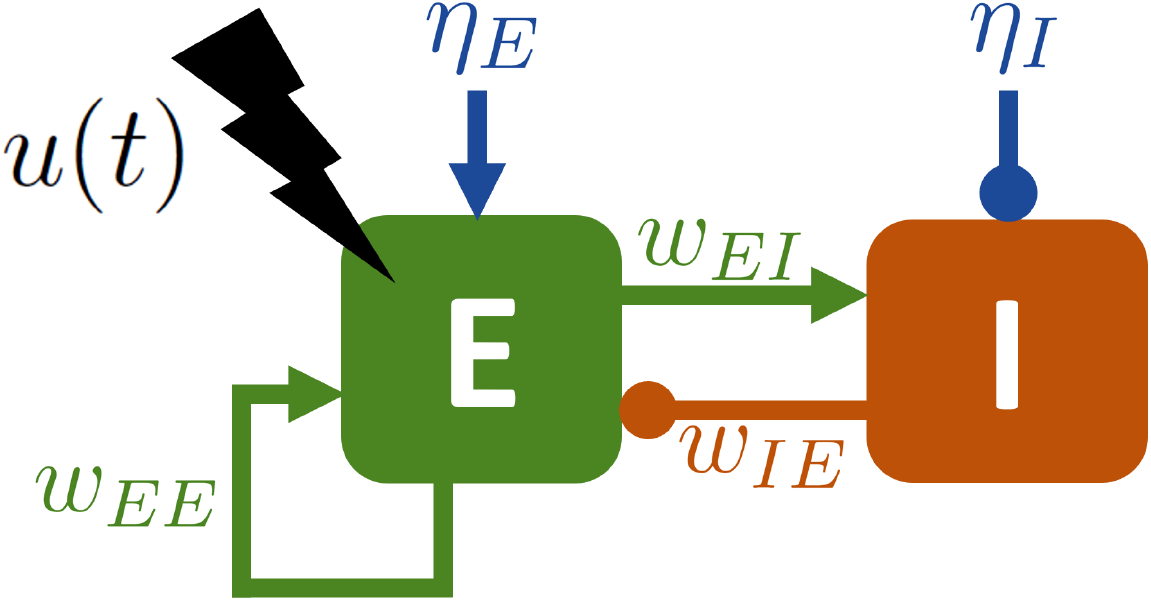
Sketch of the Wilson-Cowan model with stimulation. Intrinsic PAC can be generated by a slow oscillatory input *η*_*E*_ provided to the excitatory population (denoted E and shown in green). The inhibitory population (denoted I and shown in red) receives a constant input *η*_*I*_. The excitatory and inhibitory populations are reciprocally coupled, and the excitatory population has a self-excitatory connection. The stimulation *u*(*t*) (in black) acts on the excitatory population.

We test our predictions using two dynamically distinct cases. The first is a theta-dominant example with some theta-gamma PAC in the absence of stimulation (figure 7A) based on the parameters used in [44] (values given in table 1 in the Appendix). This case is inspired by situations where gamma is locked to the peak of theta (e.g. in the human hippocampus during memory encoding [6]), and increasing PAC could be beneficial (e.g in AD). In this case, the WC is in a fixed-point regime when the slow input is low, and crosses the Hopf bifurcation to the limit-cycle regime (gamma oscillations) when the slow input increases (see [44] for more details). We call this example the “strong theta case”. Our second case corresponds to a hypothetical scenario where PAC has almost completely disappeared due to pathology and should be restored externally by stimulating the fast rhythm. This second example displays pure gamma oscillations (limit-cycle regime) with no slow input and no PAC in the absence of stimulation (see figure 8A, parameters in table 1 in the Appendix). We call this example the “pure gamma case”. To investigate whether predictions from the theory developed using the SL model in section 2.1 carry over to the WC model, we optimise Fourier coefficients of *u*(*t*) up to *N*_*u*_ = 20 under energy constraint for both the strong theta case and the pure gamma case. Methodological details of the optimisation process can be found in section 4.2, and methodological details of the perturbation analysis can be found in section 4.3 (perturbation size is as before).

**Table 1:**
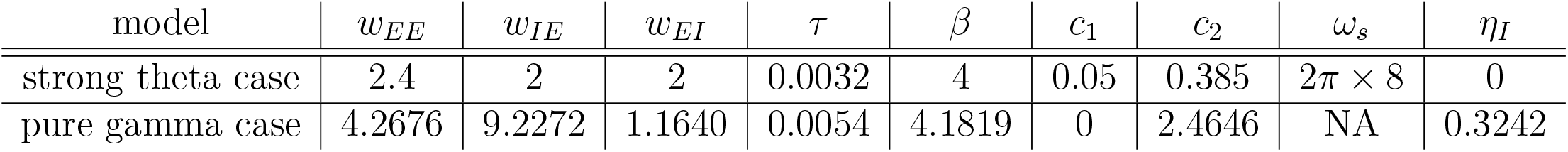
Parameters of the Wilson-Cowan model used in simulations. The strong theta case correspond to figure 7, and the pure gamma case to figure 8. Parameters of the strong theta case are taken from [44].

| model | $w_{EE}$ | $w_{IE}$ | $w_{EI}$ | $\tau$ | $\beta$ | $c_1$ | $c_2$ | $\omega_s$ | $\eta_I$ |
| --- | --- | --- | --- | --- | --- | --- | --- | --- | --- |
| strong theta case | 2.4 | 2 | 2 | 0.0032 | 4 | 0.05 | 0.385 | $2\pi \times 8$ | 0 |
| pure gamma case | 4.2676 | 9.2272 | 1.1640 | 0.0054 | 4.1819 | 0 | 2.4646 | NA | 0.3242 |

**Figure 7:**
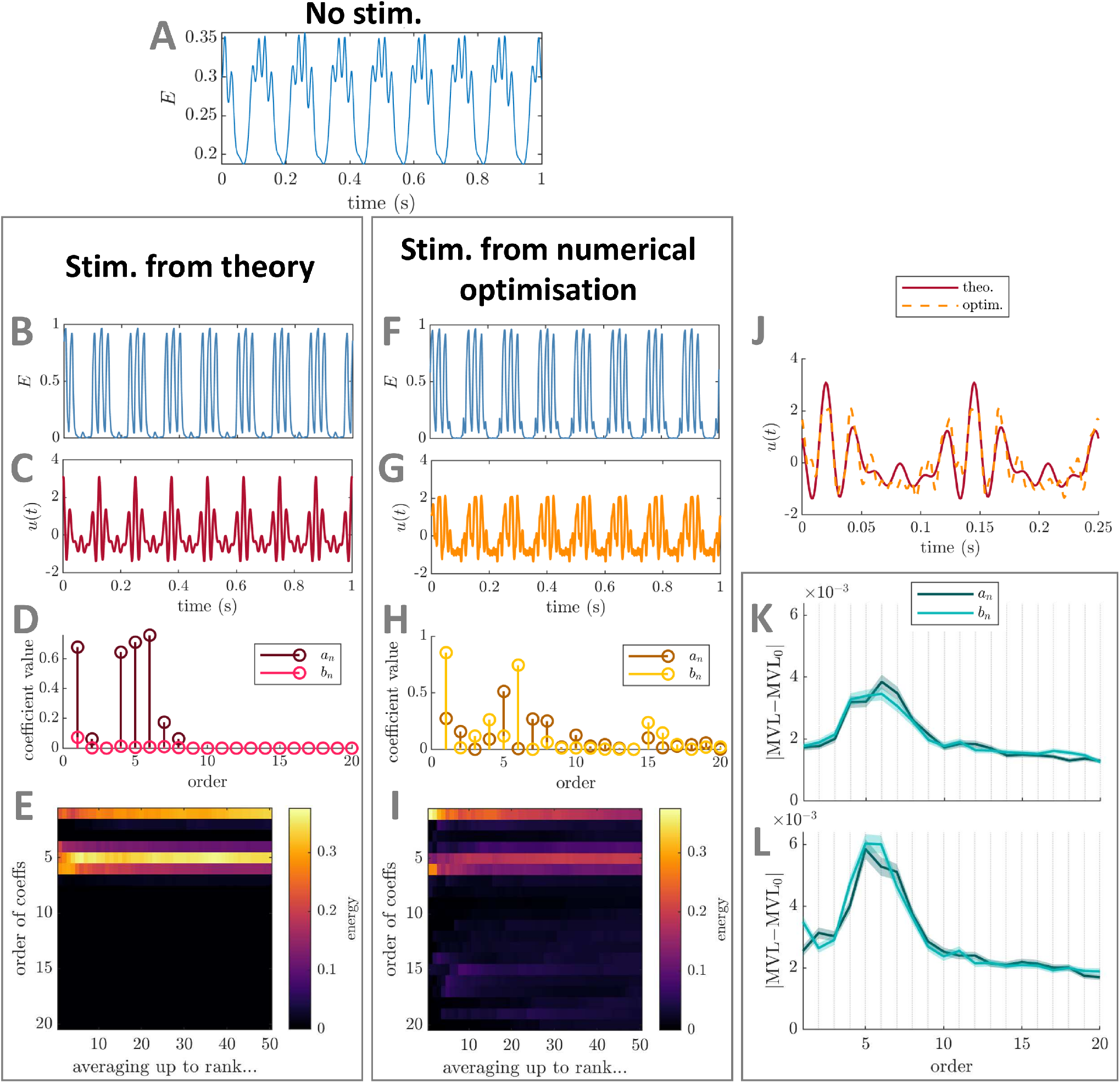
Comparison between best PAC-enhancing waveforms predicted by theory and by numerical optimisation – strong theta case in the Wilson-Cowan model. The model output in the absence of stimulation is shown in panel A. The model output when receiving PAC-enhancing stimulation is shown in panels B (best stimulation waveform obtained when optimising only Fourier coefficients predicted by theory) and F (best stimulation waveform obtained when optimising all Fourier coefficients). The corresponding best PAC-enhancing stimulation waveforms are shown in panels C and G, respectively, and are overlaid for comparison in panel J (aligned to maximise their cross-correlation). Their Fourier coefficients are shown in panels D and H, respectively. The energy of PAC-enhancing waveforms obtained from numerical optimisation for all Fourier coefficient orders (vertical axis) when averaging the *x*-best optimisation results (*x* being the horizontal axis value) is represented in panels E (only Fourier coefficients predicted by theory were optimised) and I (all Fourier coefficients were optimised). The absolute change in MVL when increasing the energy of a given stimulation Fourier coefficient is provided in panels K (when starting from PAC-enhancing waveforms obtained from the numerical optimisation process with all coefficients optimised), and L (when starting from random waveforms). Error bars represent the standard error of the mean. MVL for the stimulation waveform with only coefficients predicted by theory optimised is 0.102, MVL for the stimulation waveform with all coefficients optimised is 0.101, MVL in the absence of stimulation is 0.0045 (Δ*f*_*f*_ = 20 Hz). In all cases, waveform energy is fixed at Ξ = 1. The parameters of the Wilson-Cowan model used are taken given in Table 1 (strong theta row), and *r* = 6 (off-stimulation).

**Figure 8:**
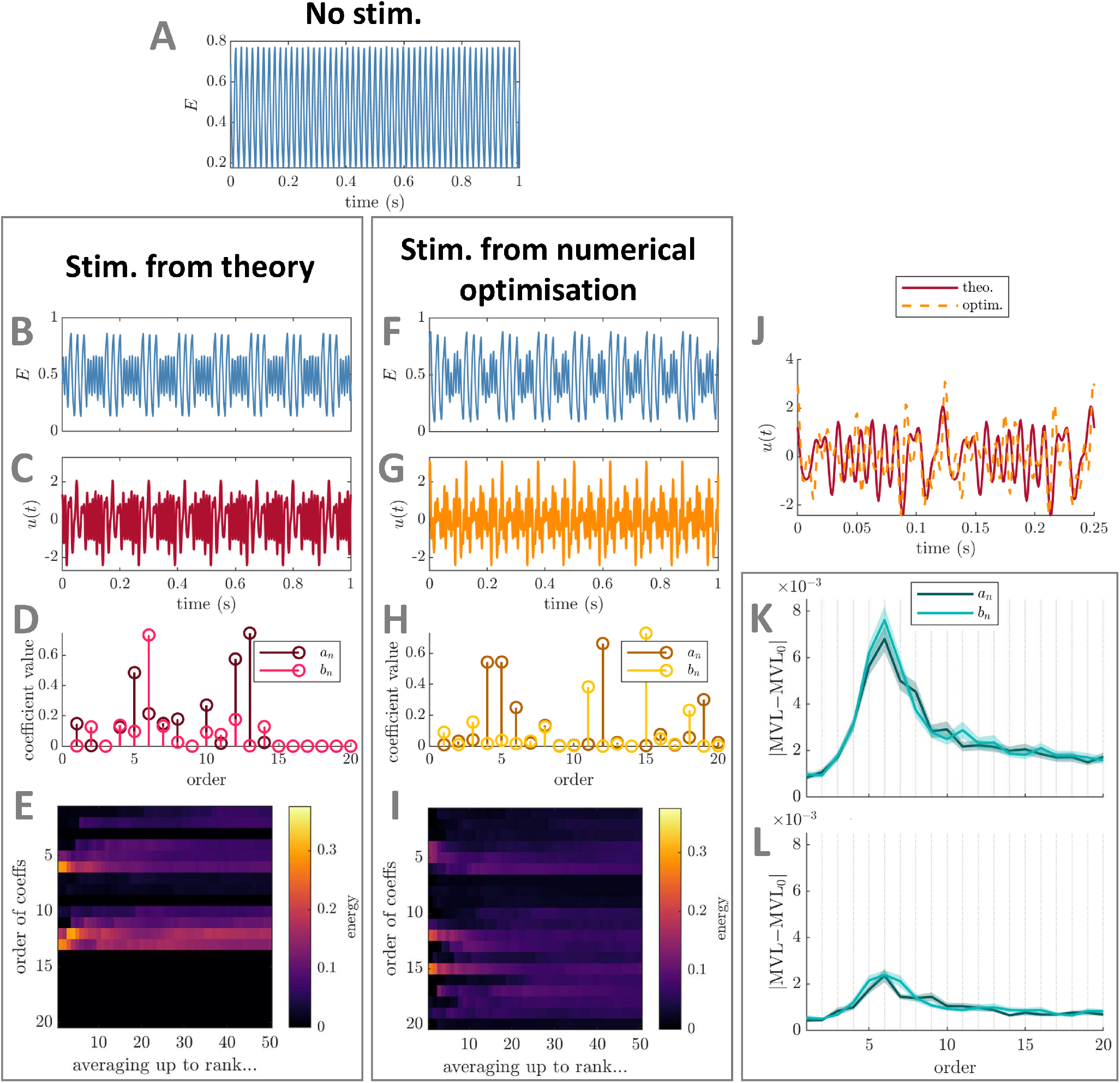
Comparison between best PAC-enhancing waveforms predicted by theory and by numerical optimisation – pure gamma case in the Wilson-Cowan model. The model output in the absence of stimulation is shown in panel A. The model output when receiving PAC-enhancing stimulation is shown in panels B (best stimulation waveform obtained when optimising only Fourier coefficients predicted by theory) and F (best stimulation waveform obtained when optimising all Fourier coefficients). The corresponding best PAC-enhancing stimulation waveforms are shown in panels C and G, respectively, and are overlaid for comparison in panel J (aligned to maximise their cross-correlation). Their Fourier coefficients are shown in panels D and H, respectively. The energy of PAC-enhancing waveforms obtained from numerical optimisation for all Fourier coefficient orders (vertical axis) when averaging the *x*-best optimisation results (*x* being the horizontal axis value) is represented in panels E (only Fourier coefficients predicted by theory were optimised) and I (all Fourier coefficients were optimised). The absolute change in MVL when increasing the energy of a given stimulation Fourier coefficient is provided in panels K (when starting from PAC-enhancing waveforms obtained from the numerical optimisation process with all coefficients optimised), and L (when starting from random waveforms). Error bars represent the standard error of the mean. MVL for the stimulation waveform with only coefficients predicted by theory optimised is 0.069, MVL for the stimulation waveform with all coefficients optimised is 0.070, MVL in the absence of stimulation is 1.3 *×* 10^−5^ (Δ*f*_*f*_ = 20 Hz). In all cases, waveform energy is fixed at Ξ = 1. The parameters of the Wilson-Cowan model used are taken given in Table 1 (pure gamma), and *r* = 6 (off-stimulation).

To test the predictions of our framework, we also compute the ARC in both cases (methodological details can be found in section 4.7 in the Appendix). In general, the ARC may depend on the full state of the system (we investigated the case of the dependence on *ρ*_*f*_ in the SL model in section 2.1.3), so for the two-dimensional WC model the ARC may also depend on the amplitude of the oscillations in addition to their phase. Here, we compute the ARC along representative trajectories selected to reflect the dynamical regimes where significant levels of stimulation are provided. For the strong theta case, this corresponds to the high amplitude regime highlighted in gray in figure 9A4. Stimulation is very low during the adjacent low amplitude regime so the ARC there is not relevant (in this case it is approximately a scaled version of the ARC in the high amplitude regime). For the pure gamma case, we compute the ARC for the two distinct dynamical regimes where significant levels of stimulation are provided: a high amplitude regime (left gray rectangle in figure 9B3), and a lower amplitude, higher frequency regime (right rectangle in figure 9B3). The details of the trajectories used to compute the ARC are shown in figure S.6 in supplementary material.

**Figure 9:**
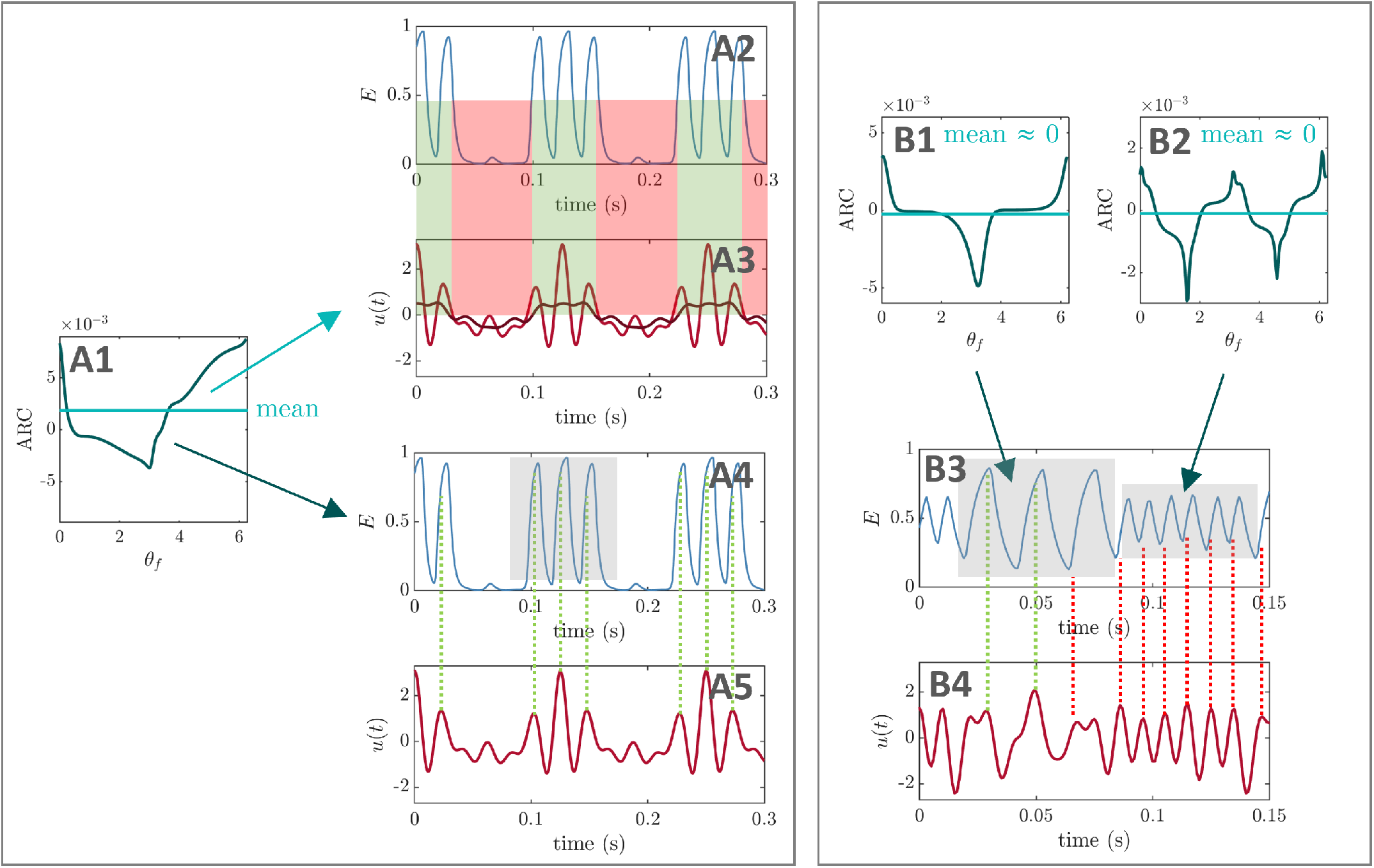
PAC-enhancing mechanisms in the Wilson-Cowan model. A corresponds to the strong theta case, and B to the pure gamma case. The ARCs shown were calculated in the regimes highlighted in grey. In A, the amplitude response of the excitatory population depends on its phase and has a non-zero mean (highlighted in light green in A1). The optimal stimulation waveform (taken from figure 7C) combines the mechanisms of PAC-enhancement corresponding to the foundational cases one and two in the SL model (slow-frequency stimulation in A2-3, and fast-frequency stimulation in A4-5). The dark red line in A3 represents a moving average of the optimal stimulation waveform (sliding window corresponding approximately to two fast-population cycles). In B, the amplitude response of the excitatory population has a mean close to zero (light green line in B1-2), but a strong phase dependence. Thus, only the mechanism corresponding to foundational case two in the SL model is at play here (fast-frequency stimulation in B3-4). The ARC strongly depends on the amplitude of the excitatory population, and the presence of a strong second harmonic in B2 leads to a strong component around twice the fast frequency in the stimulation waveform.

In the strong theta case, when optimising all Fourier coefficients of the stimulation waveform, energy is concentrated in coefficients of order 1, *r* − 1, and *r*(see figure 7H-I, *r* = 6 based on off-stimulation frequencies). Perturbation analysis highlights the role of coefficients of order *r* − 2 to *r* + 2 when starting from optimised waveforms, and of order *r* − 1 to *r* + 1 when starting from random waveforms. When only optimising coefficients of order 1, 2, and *r* − 2 to *r* + 2, we find significant energy only in coefficients of order 1 and *r* − 2 to *r*(figure 7D-E), and a slightly more favorable MVL value than when optimising all coefficients (see figure caption). In light of the ARC of the stimulated WC population, these results are consistent with the predictions obtained previously. In particular, the ARC has a non-zero mean (light green line in figure 9A1), which gives leverage to the Fourier coefficients of the stimulation waveform at the slow frequency, i.e. of order 1 (figure 9A2-3). Furthermore, the phase-dependence of the ARC is described by a strong first harmonic (with some dependence on *ρ*_*f*_), hence the key role played by coefficients belonging to orders *r* − 2 to *r* + 2 (no strong involvement of coefficients at the second harmonic). Both mechanisms of actions of optimal PAC-enhancing waveforms identified in the SL model are therefore preserved in this example (figure 9A2-A5).

In the pure gamma case, the involvement of stimulation waveform Fourier coefficients around the second harmonic of the fast frequency is much more pronounced, and the slow frequency is absent. When optimising all coefficients, energy is concentrated in coefficients of order *r* − 2 to *r*, and 2*r* to 2*r* + 3 (see figure 8H-I, *r* = 6 based on off-stimulation frequencies). Perturbation analysis underlines the impact of coefficients of order *r* − 1 to *r* + 1 (figure 8K-L). When only optimising coefficients of order 1, 2, *r* − 2 to *r* + 2, and 2*r* − 2 to 2*r* +2, we find significant energy only in coefficients of order *r* − 2 to *r*, and 2*r* − 2 to 2*r* + 1 (figure 8D-E), and a slightly less favorable MVL value than when optimising all coefficients (see figure caption). Given the ARC of the stimulated WC population, these findings align with prior predictions. Since the ARC mean is close to zero (light green lines in figures 9B1-B2), optimal stimulation waveforms have no significant energy at the slow frequency. Moreover, the ARC shows a strong first harmonic when *ρ*_*f*_ is high (figure 9B1). When *ρ*_*f*_ is low, the frequency of the fast oscillations doubles (faster frequency associated with the unstable fixed point enclosed by the limit cycle), which corresponds to a dominant second harmonic in the ARC (figure 9B2). According to the previously developed theory and given the dependence on *ρ*_*f*_, this corresponds to the potential involvement of stimulation coefficients of order *r* − 2 to *r* + 2, and 2*r* − 2 to 2*r* + 2, which is verified here. An exception to this is the 2*r* + 3 term seen in figure 8I, which may be due to the speed-up of fast oscillations at low *ρ*_*f*_ (*r* = 5 on stimulation), or to the fact that the dependence on *ρ*_*f*_ cannot be described sufficiently well by a single harmonic.

### 2.3 Practical considerations

We begin by summarising in a flowchart (figure 10A) the insights from the previous sections with a view to help experimentalists design PAC-enhancing stimulation. As a reminder, we are assuming that stimulation solely affects the fast population, and our predictions are based on the ARC of the fast population (representing the change in the amplitude of the fast population as a function of the phase of stimulation), which can be measured experimentally [45, 35]. If the amplitude response to stimulation of the fast rhythm does not depend on its phase, optimal stimulation is at the slow frequency *f*_*s*_. If the amplitude response does depend on the phase of the fast rhythm and its mean is negligible, then the optimal stimulation is a combination of the fast frequency *f*_*f*_, as well as *f*_*f*_ ± *f*_*s*_ (and corresponding harmonics if there are strong harmonics in the ARC of the fast population). Otherwise, the optimal stimulation is a combination of these two strategies. Neighbouring frequency components may be added if the result is not satisfactory, potentially indicating a dependence of the amplitude response on the amplitude of the fast population. We also note that the same framework applies if one aims to reduce rather than enhance PAC. The resulting optimal stimulation waveforms will however be different (for example, the waveform will be anti-phase in the case of slow-frequency stimulation). We next examine the trade-offs between Fourier coefficients of optimal PAC-enhancing stimulation waveforms, investigate whether phase-locking to the slow and/or fast rhythms is necessary, and how to approximate optimal PAC-enhancing stimulation waveforms using pulses.

**Figure 10:**
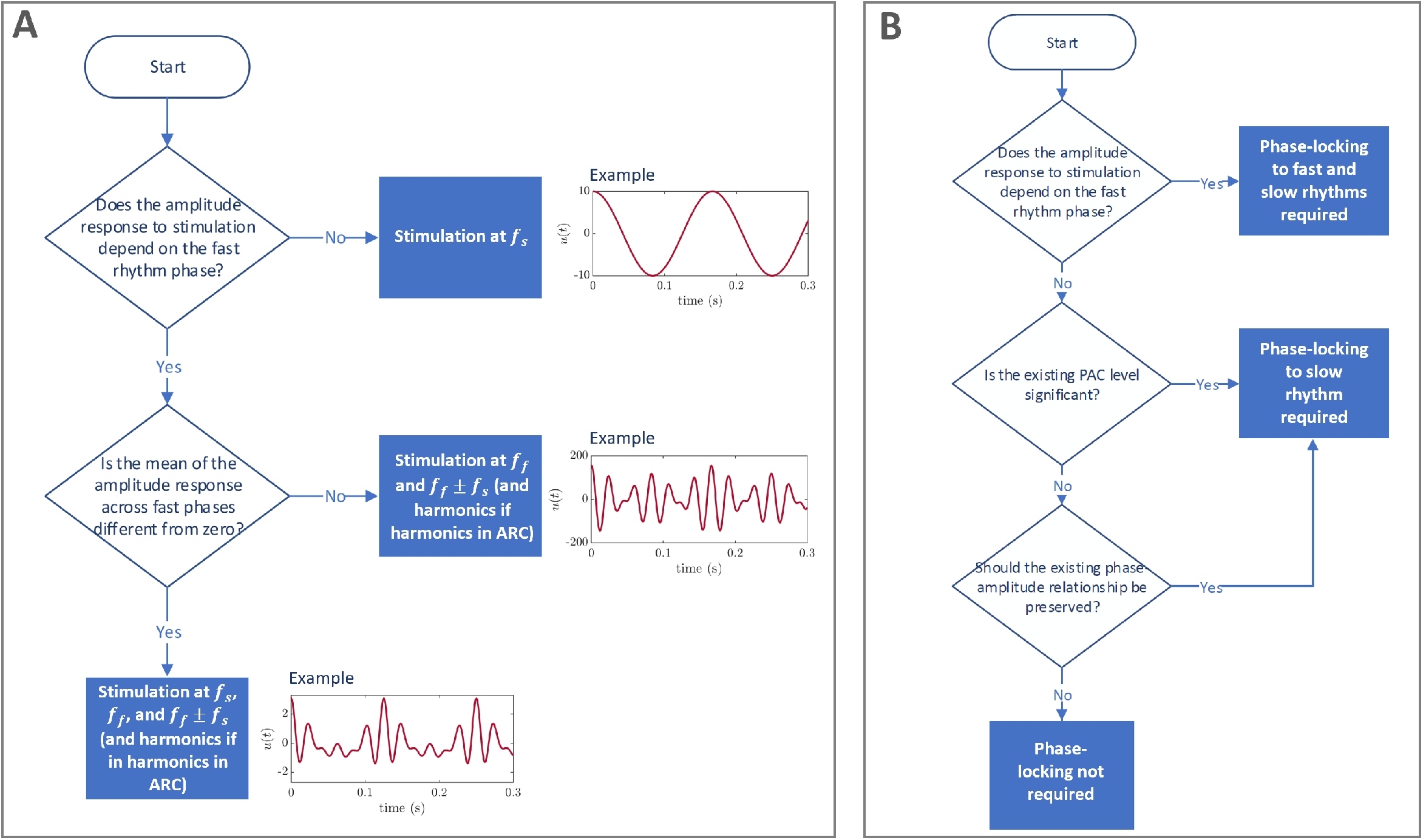
Simplified flowcharts to guide the design of optimal PAC-enhancing stimulation. We are assuming that stimulation acts solely on the fast population. The flowchart in panel A presents which Fourier coefficients of the stimulation waveform to optimise. Neighbouring frequency components may be added if the result is not satisfactory, potentially indicating a dependence of the amplitude response on the amplitude of the fast population. The flowchart in panel B assumes that stimulation does not significantly entrain the slow rhythm and presents a guide to decide whether phase-locking the stimulation to the fast and/or slow rhythms is necessary.

#### 2.3.1 Trade-offs between Fourier coefficients of the stimulation waveform

The theory developed in section 2.1 identifies which Fourier coefficients should be considered to build an optimal PAC-modulating stimulation waveform, but does not prescribe how much energy should be assigned to these coefficients (except in the simplest case of stimulation being coupled through the mean-field of the fast population, where only one Fourier component is involved). From the mechanism illustrated in figure 2B, the waveform can be manually designed such that where the amplitude of the fast oscillation should be increased, the phase alignment between the fast component in the stimulation waveform and the fast rhythm correspond to the maximum amplification in the ARC. Conversely, where the amplitude of the fast oscillation should be decreased, the phase alignment between the fast component in the stimulation waveform and the fast rhythm should correspond to the maximum suppression in the ARC. Here, we investigate numerically whether other principles can be found to guide the design of optimal PAC-modulating stimulation waveforms.

In particular, we aim to contrast optimal PAC-enhancing stimulation for different levels of endogenous fast oscillations and slow input. To this end, we consider the SL model (equations (1) and (3)), with stimulation coupled to the fast population through ARC(*θ*_*f*_) = 0.5 + cos(*θ*_*f*_) and PRC(*θ*_*f*_) = sin(*θ*_*f*_) for simplicity. As previously outlined, the level of endogenous fast oscillations is controlled in this model by *δ*, and the level of slow oscillations is controlled by *k*_*s*_.

To efficiently optimise the stimulation waveform across combinations of *δ* and *k*_*s*_, we simplify the optimisation problem as follows. We parametrise the stimulation waveform Fourier series as

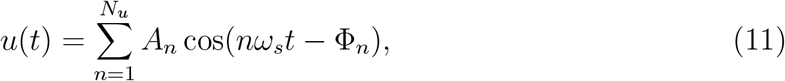

where the amplitudes *A*_*n*_ are positive, and the phases Φ_*n*_ are in [0, 2*π*). In the SL model considered, only Fourier coefficients of order 1, *r* − 1, *r*, and *r* + 1 significantly impact PAC levels (see section 2.1.3 with *N*_*α*_ = 1 and *g*(*ρ*_*f*_) = 1). We therefore perform a parameter sweep for combinations of *A*_1_, *A*_*r* −1_, and *A*_*r*_ (*A*_*r*+1_ is obtained by matching the target stimulation energy, i.e.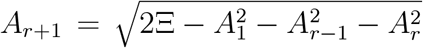). For each combination of amplitudes, we only need to optimise the phases Φ_1_, Φ_*r* −1_, Φ_*r*_, and Φ_*r*+1_. These are simpler optimisations than the method described in section 4.2 because there is no non-linear constraint to enforce by the optimiser, and there are only four parameters to optimise. Because of the coarse-graining of amplitudes, this approach is less precise than the full optimisations performed previously, but is more efficient which allows us to explore changes in optimal stimulation waveform as a function of the strength of endogenous fast and slow oscillations in the model. We use 10 equally spaced amplitude values for each of *A*_1_, *A*_*r* −1_, and *A*_*r*_ (i.e. 1000 amplitude combinations), and perform five local optimisations per combination. Other methodological details are unchanged and per section 4.2 in the Appendix.

Changes in the balance between Fourier amplitudes as a function of the strength of endogenous fast and slow oscillations in the model (figure 11A-C) are relatively minor but can be explained intuitively. As endogenous fast oscillations become stronger (increase in *δ*), the Fourier amplitudes corresponding to the fast frequency (*A*_*r*_) and the slow frequency (*A*_1_) decrease in favor of the Fourier amplitudes corresponding to the modulation of the fast frequency at the slow frequency (*A*_*r* −1_ and *A*_*r*+1_). There is relatively less endogenous modulation, so an increase in the modulation of the fast frequency is necessary to bring down the trough of *ρ*_*f*_ as per the mechanism described in foundational case two (section 2.1.2 and figure 2B). Conversely, provided that endogenous fast oscillations are relatively weak (low *δ*), as endogenous slow oscillations become stronger (increase in *k*_*s*_), the Fourier amplitudes corresponding to the modulation of the fast frequency at the slow frequency (*A*_*r* −1_ and *A*_*r*+1_) decrease in favor of the Fourier amplitude corresponding to the fast frequency (*A*_*r*_). Because endogenous modulation relative to fast-frequency activity is already high, less modulation is needed from stimulation and boosting the fast frequency is advantageous. We choose the model parameters studied in figure 11A-C to cover a broad range of relative strength of fast to slow oscillations (model output in the absence of stimulation across parameters shown in figure S.7 in the Supplementary Material). The optimal PAC-enhancing waveforms resulting from the optimisations, as well as the on-stimulation model outputs are shown in figure S.8 and S.9 in the Supplementary Material, respectively. While small changes in the balance of Fourier amplitudes cannot be detected due to coarse-graining, examination of the cost (here −MVL) across Fourier amplitudes (figure 11D) supports the convergence to a single local minimum (for a given set of model parameters). Furthermore, the shift of the entire area of low cost in the space of Fourier amplitudes confirms the trends described above. Figure 11D is given for *A*_*r* −1_ = 55.56, but similar results are obtained across all *A*_*r* −1_ considered in the sweep.

**Figure 11:**
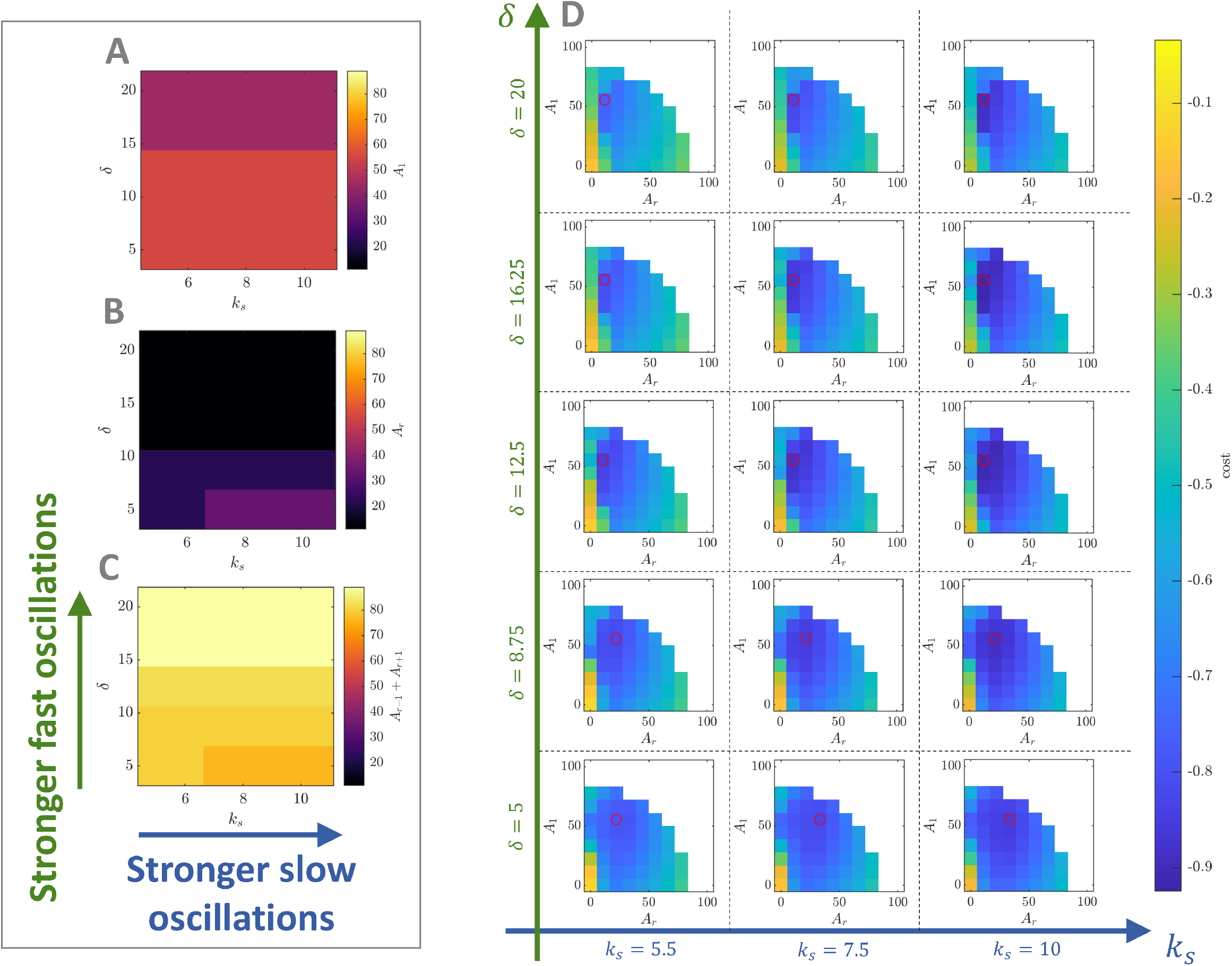
Balance between optimal Fourier amplitudes and cost landscape as a function of the strength of endogenous fast and slow oscillations in the Stuart-Landau model. The strength of endogenous slow oscillations is controlled by model parameter *k*_*s*_ (blue arrows), and the strength of endogenous fast oscillations by model parameter *δ*(green arrows). The balance between Fourier amplitudes of PAC-enhancing optimal stimulation waveforms as a function of the strength of endogenous fast and slow oscillations is shown in panels A-C. Panel A corresponds to the slow-frequency Fourier amplitude (*A*_1_), panel B to the fast-frequency Fourier amplitude (*A*_*r*_), and panel C to the modulation of the fast frequency at the slow frequency (*A*_*r* −1_ + *A*_*r*+1_). Panel D shows in color the best objective function values (costs) resulting from optimising Fourier phases to enhance PAC for *A*_*r* −1_ = 55.56, as a function of *A*_1_ and *A*_*r*_, and as a function of the strength of endogenous fast and slow oscillations (specific values indicated on the blue and and green axes). For a given combination of *k*_*s*_ and *δ*, the minimum cost for the *A*_*r* −1_ slice shown is highlight by a red circle. In all panels, the total stimulation waveform energy is kept at Ξ = 5000, and the frequency of endogenous oscillations is *f*_*f*_ = 42 Hz, and *f*_*s*_ = 6 Hz. Stimulation is coupled to the fast population through ARC(*θ*_*f*_) = 0.5 + cos(*θ*_*f*_) and PRC(*θ*_*f*_) = sin(*θ*_*f*_).

#### 2.3.2 Is phase-locking to the slow rhythm necessary?

In the theory and examples presented in this work, stimulation is provided with a period corresponding to the slow-oscillation frequency. Thus, stimulation is phase aligned to the slow rhythm at steady-state, and the specifics of the phase alignment are dictated by the Fourier component make-up of the stimulation waveform. With practical applications in mind, we investigate in this section whether phase-locking to the slow-rhythm is necessary to modulate PAC.

To this end, we simulate the SL model with stimulation coupled to the fast population through its mean-field (equation (4), see section 2.1.1). We already know from the analytical analysis in section 2.1.1 that the optimal stimulation is a sinusoid with its peak aligned to the peak of the slow rhythm. However it is unclear how essential this optimal phase alignment is. To answer this question, we provide the optimal PAC-enhancing stimulation at various phases of the slow rhythm and measure resulting PAC levels. We also perform the same analysis in the SL model with stimulation directly coupled to the fast population (as in section 2.1.2), thereby investigating the two foundational cases presented in this work.

The importance of phase alignment between stimulation and the slow rhythm depends on PAC levels in the absence of stimulation in both cases. When off-stimulation PAC levels are low (figure 12A3 and B3), a significant PAC-enhancing effect can still be achieved without phase alignment (figure 12A1 and B1). However, when off-stimulation PAC levels are higher (figure 12A4 and B4), providing stimulation close to the optimal phase is critical to enhance PAC (figure 12A2 and B2). For example, providing stimulation half a period too late/early leads to a significant decrease in PAC. We note that the fast-frequency oscillations can be entrained by stimulation in the direct coupling case, but not in the mean-field coupling case (because the PRC is zero). Regardless of the stimulation coupling function, our simulations assume that stimulation does not affect the slow rhythm, and therefore cannot entrain it. Generally, enhancing the existing phase-amplitude relationship between two rhythms requires phase locking, but may be more physiological. This is the case in figure 12A2 and B2 where overriding the existing phase-amplitude relationship would require too much energy, and enhancing the existing phase-amplitude relationship is the only viable strategy. Conversely, a phase-amplitude relationship different from the existing phase-amplitude relationship between the fast and slow rhythms is enforced by stimulation for a stimulation phase of e.g. *π* in figure 12A2 and B2.

**Figure 12:**
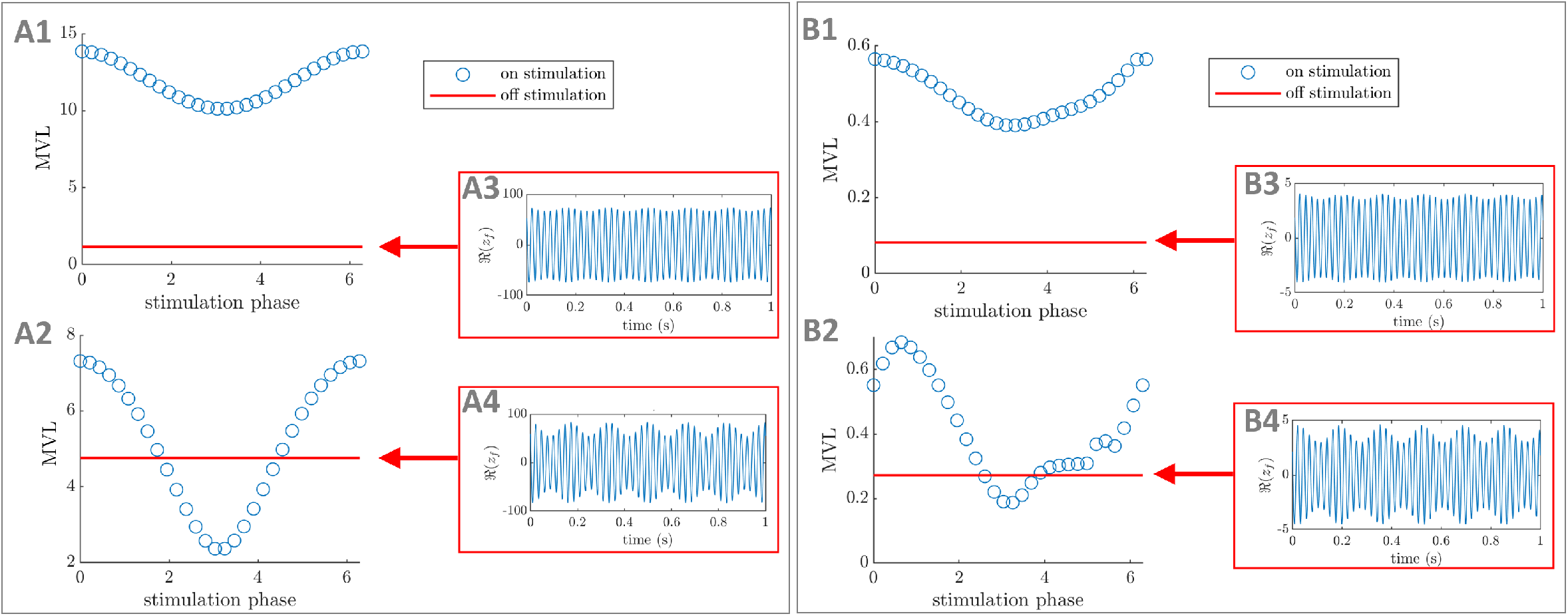
The importance of phase alignment between stimulation and the slow rhythm depends on off-stimulation PAC levels. PAC-enhancing stimulation waveforms are provided at different phases of the slow rhythm. Panels A1-4 correspond to the SL model with stimulation coupled to the fast population through its mean-field and the stimulation waveform given in figure 3C, while panels B1-4 correspond to the SL model with direct stimulation coupling and the stimulation waveform given in figure 4C. Panels A1, A2 and B1, B2 show the MVL as a function of stimulation phase of the slow rhythm in blue (a stimulation phase of zero corresponds to the peaks of the stimulation waveform and the slow rhythm being aligned), and the off-stimulation MVL level in red. Panels A3, A4 and B3, B4 represent the corresponding off-stimulation model output (real part of the order parameter). In A1-4, *f*_*f*_ = 40Hz, *f*_*s*_ = 6Hz, and *δ* = 5000. Panels A1, A3 correspond to *k*_*s*_ = 500, and Ξ = 1 *×* 10^7^. Panels A2, A4 correspond to *k*_*s*_ = 2000, and Ξ = 5 *×* 10^5^. In B1-4, *f*_*f*_ = 42Hz, *f*_*s*_ = 6Hz, *δ* = 15, and Ξ = 5000. Panels B1, B3 correspond to *k*_*s*_ = 3, and panels B2, B4 to *k*_*s*_ = 10. Note that the stimulation waveform provided in B is optimal for B1 but not for B2.

#### 2.3.3 Is phase-locking to the fast rhythm necessary?

In situations where stimulation acts through the mechanism described in foundational case two (see figure 2B), the differential alignment (as prescribed by the ARC) of the stimulation fast frequency components with the fast rhythm at the peak and trough of the slow rhythm is critical. In the ideal case where *f*_*f*_ is constant and an integer multiple of *f*_*s*_, this alignment is enforced as a consequences of phase-aligning stimulation with the slow rhythm. However, we show that in the more realistic scenario where *f*_*f*_ is not an integer multiple of *f*_*s*_ or significantly varies over time, adapting stimulation to the frequency fluctuations of the fast rhythm will give better results. To this end, we simulate the SL model as in the previous section (stimulation coupling corresponding to foundational case two) for different values of *f*_*s*_, as well as with *f*_*f*_ varying according to a Wiener process.

If the stimulation waveform (optimised for *f*_*f*_ = 42*Hz*) does not change, the maximum achievable PAC modulation decreases as *f*_*f*_ is varied from 42Hz (figure 13). The optimal phase alignment between the stimulation waveform and the slow rhythm also changes (figure 13A1 and B1) Similarly, we show that when *f*_*f*_ varies according to a Wiener process, increasing the level of noise drastically reduces the ability of open-loop stimulation to modulate PAC (figure S.10 in Supplementary Material). Together, these results suggest that if the stimulation waveform contains fast frequency components, it is advisable to lock these with the fast rhythm (in the manner prescribed by the ARC of the fast rhythm). We summarise the conclusions of this subsection and the previous one in the flowchart presented in figure 10B.

**Figure 13:**
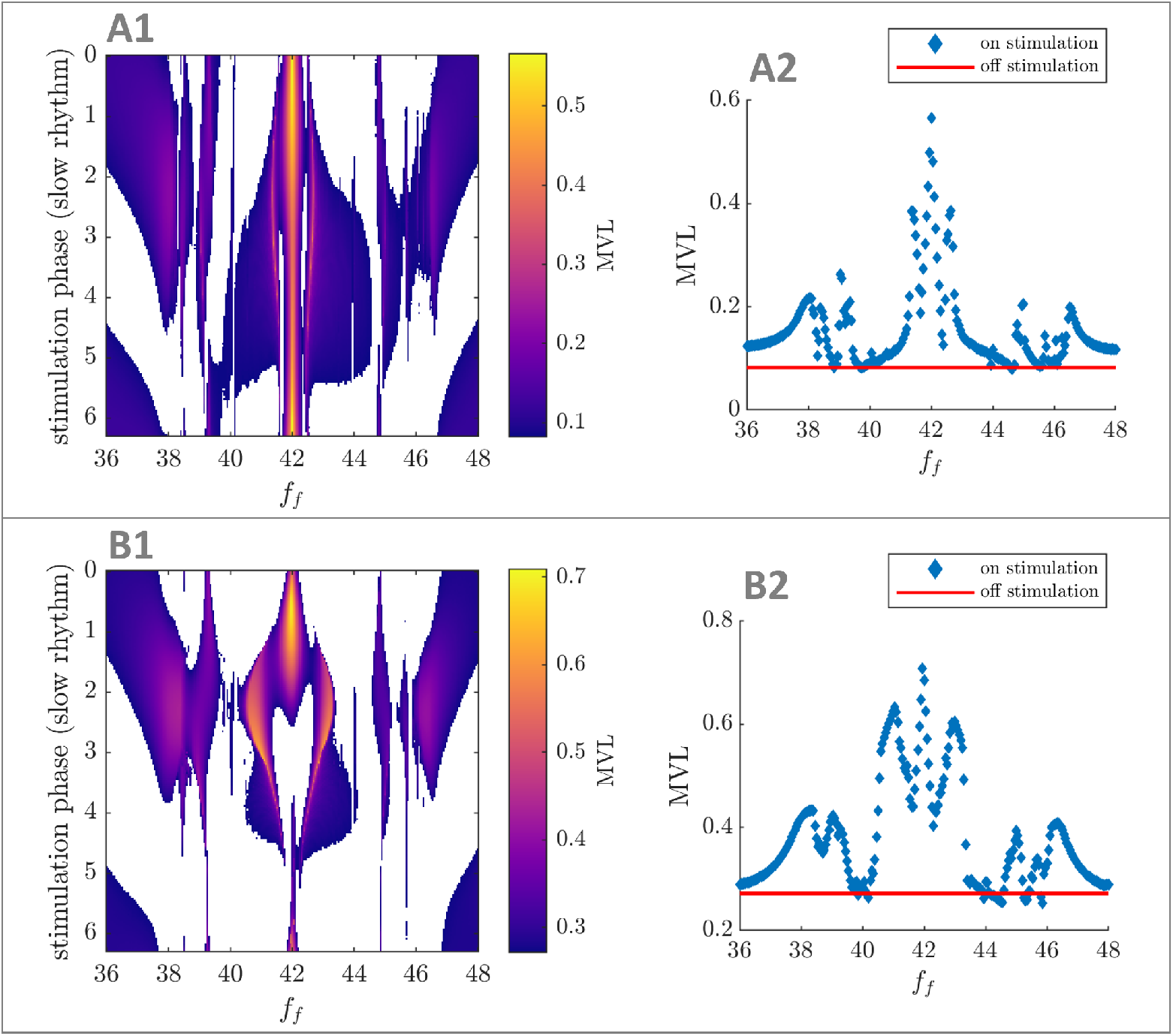
Effect of changes in the fast rhythm frequency on PAC modulation. The PAC-enhancing stimulation waveform optimised for *f*_*s*_ = 6Hz and *f*_*f*_ = 42Hz was provided at different phases of the slow rhythm, and for different values of the fast rhythm frequency. Panels A1 and B1 show the MVL as a function of *f*_*f*_, and of the stimulation phase of the slow rhythm (a stimulation phase of zero corresponds to the peaks of the stimulation waveform and the slow rhythm being aligned). No color is shown when the MVL is below the off-stimulation value. The optimal stimulation phase relative to the slow rhythm depends on *f*_*f*_, and the maximum achievable MVL decreases away from *f*_*f*_ = 42 (in the absence of adjustment of the stimulation to the fast rhythm). This is confirmed in panels A2 and B2, where the maximum achievable MVL for a given value of *f*_*f*_ is shown on the vertical axis (off-stimulation MVL level in red). In all panels, simulations are performed using an SL model with direct stimulation coupling and the stimulation waveform given in figure 4C. Parameters used are *f*_*s*_ = 6Hz, *δ* = 15, and Ξ = 5000. Panels A1-2 correspond to *k*_*s*_ = 3, and panels B1-2 to *k*_*s*_ = 10.

### 2.4 Pulsatile waveforms

Optimal waveforms parametrised by Fourier coefficients may be applicable to stimulation modalities such as tACS, but are not directly applicable to stimulation modalities that can only generate square pulses (e.g. DBS). However, these smooth optimal waveforms can be approximated using pulsatile waveforms. We suggest different ways of doing so, and compare the resulting pulsatile waveforms to the corresponding optimal smooth waveforms in terms of their effects on PAC in the SL and WC models presented before. We approximate smooth waveforms using regularly spaced square pulses, with a certain pulse frequency and pulse duration. The amplitude (intensity) of each pulse is simply given by the amplitude of the smooth waveform at the center of the pulse (as shown for e.g. figure 14E1), with a scaling factor determined to either match the energy of the smooth waveform (i.e. ∫*u*(*t*)^2^d*t*), or its cumulative absolute intensity (i.e. ∫|*u*(*t*)|d*t*). Model simulation with pulsatile waveforms required to use Euler’s method, as a variable step solver (used in the rest of this work) would lead to pulse durations varying with the integration step.

**Figure 14:**
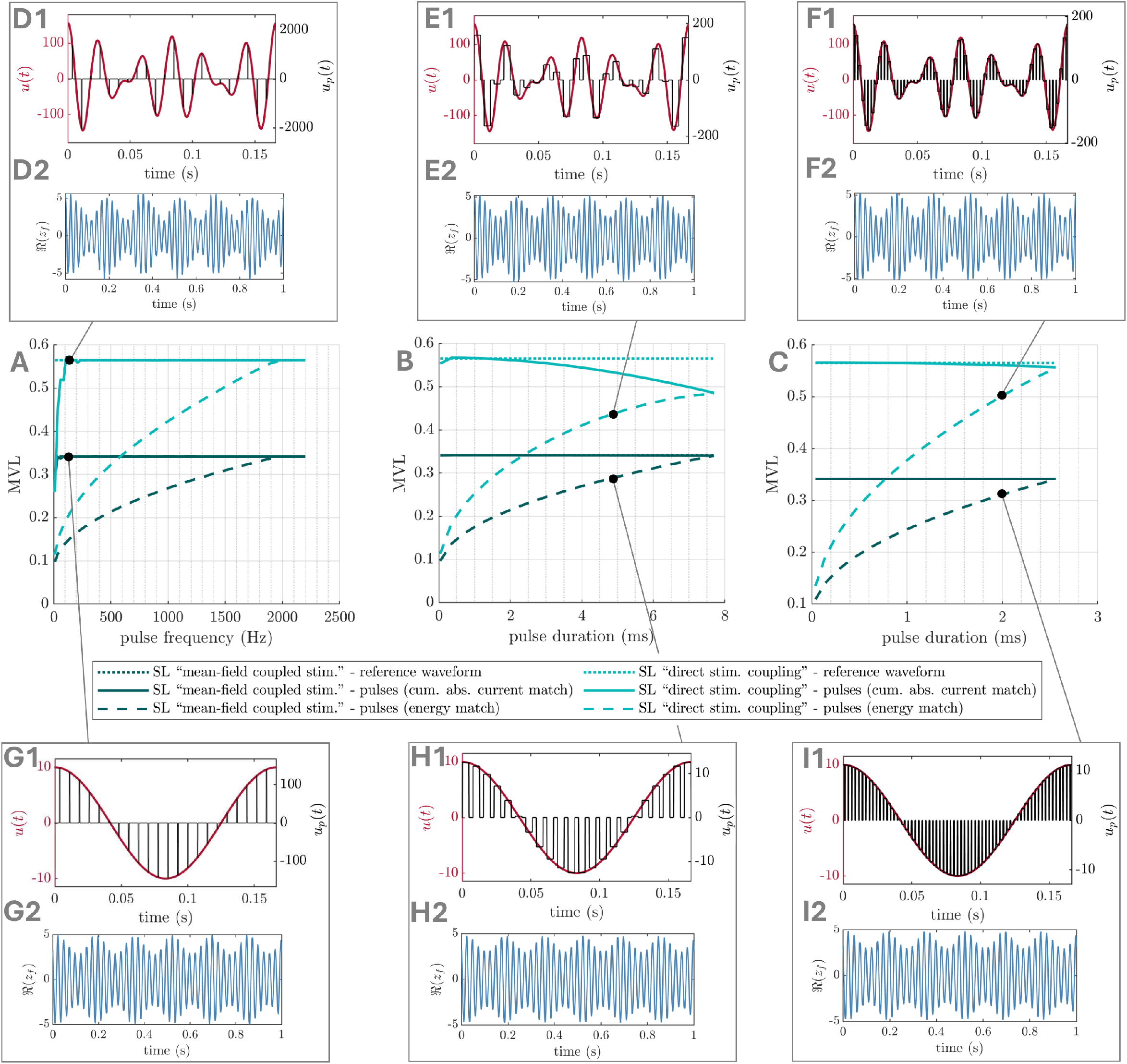
Optimal smooth waveforms can be approximated with pulses – foundational cases one and two in the SL model. MVL is shown as a function of pulse frequency, for a pulse duration of 0.5ms (A), and as a function of pulse duration, for a pulse frequency of 130Hz (B) and 390Hz (C). In these panels, foundational case one (mean-field coupled stimulation, parameters corresponding to figure 3B-D) is shown in dark green, and foundational case two (direct stimulation coupling, parameters corresponding to figure 4B-D) is shown in light green. Solid lines correspond to pulsatile waveforms obtained by matching the cumulative absolute intensity of the optimal smooth waveforms, while dashed lines correspond to pulsatile waveforms obtained by matching the energy of the optimal smooth waveforms. Dotted lines (behind the solid lines where they are not visible) correspond to the smooth optimal waveforms. Panels D-I show the smooth optimal waveform in red and the pulsatile approximation in black denoted by *u*_*p*_(*t*) (top), as well as the resulting activity of the fast population (bottom). Pulse frequencies/durations are as follow: 135Hz/0.5ms in D and G, 130Hz/4.8ms in E and H, 390Hz/2.0ms in F and I.

For the SL models investigated in foundational cases one (section 2.1.1) and two (section 2.1.2), relatively low pulse frequencies (135Hz for 0.5ms pulse duration) are sufficient for the corresponding pulsatile waveform to increase PAC as much as the smooth optimal waveform when matching cumulative absolute intensity (figure 14A). When matching waveform energy, pulse frequency has to be markedly increased to notably affect PAC (figure 14B); alternatively, pulse duration can be lengthened while maintaining a low pulse frequency (figure 14B-C). For the SL models investigated in the general stimulation coupling case (section 2.1.3), higher pulse frequencies are generally required to match the effects on PAC of the smooth optimal waveforms (around 360Hz for 0.5ms pulse duration, see figure S.11 in supplementary material). This is also the case with the WC models investigated in section 2.2, see figure S.12 in supplementary material. However, increasing pulse duration can considerably lower the pulse frequency required (figure 15B-C). Moreover, irregular pulse spacing chosen such that pulses are centered on the local peaks of the smooth waveform (as shown in figure 15D1 and G1) can further reduce the (average) pulse frequency required to 75Hz (10ms pulse) for the strong theta case, and to 197Hz (3ms pulse) for the pure gamma case (figure 15A).

**Figure 15:**
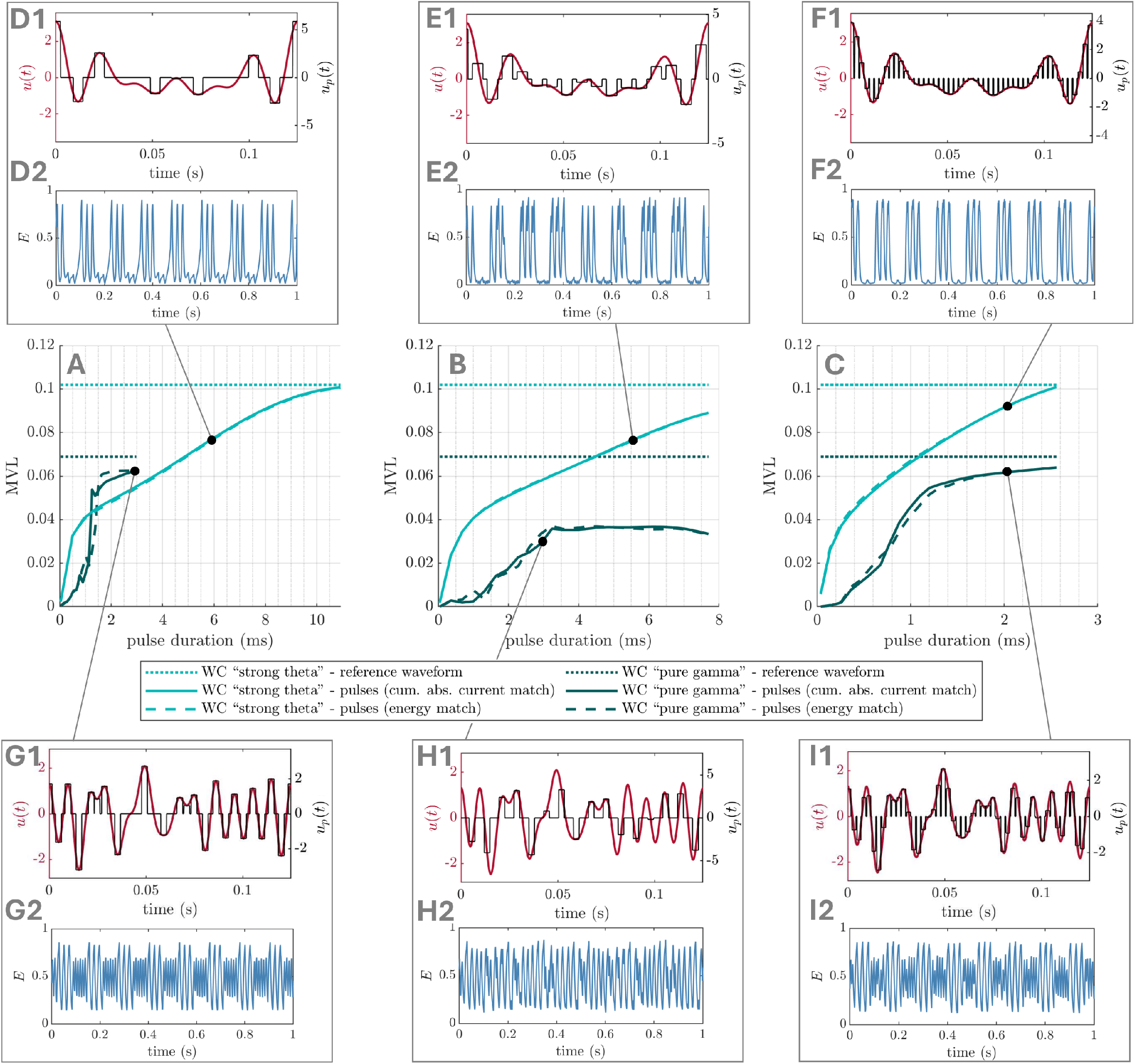
Optimal smooth waveforms can be approximated with pulses – WC model. MVL is shown as a function of pulse duration for pulses centered on the local peaks of the smooth waveform (A), as well as for a frequency of 130Hz (B) and 390Hz (C). In these panels, the strong theta case (parameters corresponding to figure 7B-D) is shown in light green, and the pure gamma case (parameters corresponding to figure 8B-D) is shown in dark green. Solid lines correspond to pulsatile waveforms obtained by matching the cumulative absolute intensity of the optimal smooth waveforms, while dashed lines correspond to pulsatile waveforms obtained by matching the energy of the optimal smooth waveforms. Dotted lines correspond to the smooth optimal waveforms. Panels D-I show the smooth optimal waveform in red and the pulsatile approximation in black (top), as well as the resulting activity of the fast population (bottom). Pulse frequencies/durations are as follow: 75.4Hz(average)/5.9ms in D, 130Hz/5.8ms in E and I, 390Hz/2.0ms in F, 197Hz(average)/3.0ms in G, and 130Hz/2.9ms in H.

## 3 Discussion

In this work, we developed a framework to guide the development of optimal PAC-enhancing stimulation. Our framework is for stimulation acting on the neural population generating the fast rhythm, and assumes that neither stimulation nor the fast rhythm significantly affect the slow rhythm (assumed to be generated by another neural population). Using a SL model, we showed that the ARC of the fast population determines which Fourier coefficients should be included and optimised in the stimulation waveform. Specifically, if the amplitude response to stimulation of the fast rhythm does not depend on its phase, optimal stimulation is at the slow frequency *f*_*s*_ (figure 2A). If the amplitude response of the fast rhythm does depend on its phase and its mean is negligible, then the optimal stimulation is a combination of the fast frequency *f*_*f*_, as well as *f*_*f*_ ± *f*_*s*_ (and corresponding harmonics if there are strong harmonics in the amplitude response curve of the fast population), see figure 2B. Otherwise, the optimal stimulation is a combination of these two strategies (figure 2C). Neighbouring frequency components may be added if the result is not satisfactory, potentially indicating a dependence of the amplitude response on the amplitude of the fast population.

Additionally, the predictions obtained with the SL model appeared to carry over in several dynamical regimes of a more realistic neural mass model representing interacting neural populations, the WC model (figure 9). Moreover, we showed in the SL model that changes in the balance between Fourier amplitudes as a function of the strength of endogenous fast and slow oscillations are relatively minor but can be explained intuitively (figure 11). We also established that when stimulation includes fast frequency component, it is likely that locking these with the fast population (as specified by the ARC) will be necessary. When stimulation does not include fast components, the importance of phase alignment between stimulation and the slow rhythm depends on PAC levels in the absence of stimulation, and on whether overriding the existing phase-amplitude relationship is acceptable (figure 12). Finally, for neuromodulation modalities that can only generate square waves, the optimal waveforms predicted by our framework can be approximated by pulsatile waveforms.

### Modelling PAC generation

Models with various levels of biophysical details have been used to investigate PAC. For example, detailed single and multi-compartment models can generate PAC (see [46] for a review), and the emergence of PAC was investigated in a simulated hippocampal CA1 microcircuit with morphologically detailed neurons [47]. Neural mass models with various types of coupling can also produce PAC, from simple E-I loops [44] to realistic circuits comprising four cortical layers and dozens of populations [48]. The canonical types of population interactions leading to PAC have been reviewed in [49] (our models correspond to unidirectional coupling from a slow population to a fast population), and the bifurcation types responsible for PAC in these models are studied in [50]. Importantly, the effects of brain stimulation on PAC were only explored in a couple of modelling studies to date, namely in a neuronal network consisting of one thousand cells simulated in NEURON [51], and recently in a model connecting a biophysically-detailed representation of the hippocampus with Kuramoto oscillators portraying input from the medial septum [52].

We chose the SL model for its ability to represent a neural oscillator with a phase and an amplitude variable going through a Hopf bifurcation, and for its analytical tractability which allowed us to gain insights into optimal PAC-enhancing stimulation. We chose the WC model to test the predictions obtained with the SL model since the WC model has been proposed as a canonical E-I circuit to generate PAC [44] and has been commonly used to study neural oscillations and optimise therapeutic brain stimulation [53, 54, 44, 55, 56, 57, 58, 59, 60]. Crucially, the WC model is a relatively inexpensive to simulate neural mass model (as opposed to models requiring to simulate individual neurons), which makes numerical optimisation of the stimulation waveform possible. Additionally, in both our SL and WC models, the fast rhythm can be periodically inhibited by the slow rhythm as in detailed neurons models reviewed in [46], but the fast population can also be quiescent (*δ* ≤ 0 in the SL model and trough of the strong theta regime in the WC model). In that case, fast oscillations are only brought about by the rising slow input causing the model to traverse a Hopf bifurcation [44, 27] (see the WC model strong theta case in section 2.2).

### Comparing predictions with experimental data

Results of recent experimental studies are in line with some of the predictions made in this work. In particular, bursts of stimulation phased-locked to the peak of the slow rhythm were found to increase PAC compared to baseline and to stimulation provided at the trough of the slow rhythm [25, 26]. These could correspond to purely excitatory pulses acting through the mechanism presented in section 2.1.1 (see figure 2A), or to pulses with excitatory and inhibitory components acting through the mechanism presented in section 2.1.2 (see figure 2B) where phase-alignment with the peak of the fast oscillations happens through entrainment. Other studies reported improvements in memory performance [61], motor skill acquisition [62], and cognitive task performance [63] after open-loop transcranial alternative current stimulation with bursts of gamma stimulation superimposed to the peak of a theta stimulation waveform. No improvement was reported in [61, 62] when gamma bursts are superimposed to the trough of the theta stimulation waveform. Although PAC was not measured during stimulation, these behavioural improvements are likely mediated by an increase in PAC. The effective stimulation waveform in these studies correspond to the combination of mechanisms one and two, and is similar to the optimal waveform in the WC model strong theta case (see figure 9A). Since stimulation was open-loop, entrainment of both the slow and fast rhythms may have played a role. Furthermore, the additive effect of both mechanisms (i.e slow and fast components of the stimulation waveform) on memory performance was confirmed in [61], as well as the frequency specificity of the fast component of the stimulation waveform.

Nevertheless, more experimental work is required to validate our framework, in particular with regards to the relationship between the characteristics of the ARC of the fast population and the optimal PAC-enhancing waveform. The ARC of the fast population of interest could be measured experimentally using phase-locked stimulation as in previous studies [45, 35]. Recent advances in continuous real-time phase estimation with zero filter delay [64] (also see link to code in [35]) make phase-locking to fast oscillations feasible (robust phase-locking was achieved at 40 Hz in [35]). Another method was recently shown to reliably estimate in real-time the phase of oscillations up to 250 Hz in synthetic data [65]. These advances will also be key to phase-locking stimulation to the fast rhythm according to the mechanism described in figure 2B.

### Limitations

In this work, the slow rhythm involved in PAC was considered to be generated by an external neural population (for example by the medial septum in the case of hippocampal theta-gamma PAC). It was assumed that neither the fast population nor stimulation significantly influence the slow rhythm. This is justified if stimulation is local to the fast population, and the influence of the fast population on the slow rhythm averages out on the slow timescale [27] or the projections from the fast population to the slow population are weak. However, if stimulation significantly affects the slow rhythm, the impact on PAC can be substantial as shown recently [52]. Including in our framework the potential effects of stimulation on the slow rhythm will be a focus for future work. Additionally, how our framework may generalise to more detailed models of neural populations has not been studied beyond the WC. Since the SL model used to develop the framework is the normal form of a Hopf bifurcation, we can speculate that our predictions may hold in more detailed models (e.g. including detailed representations of individual neurons) that operate in the vicinity of a Hopf bifurcation.

Further limitations of this study include the absence of noise and the absence of synaptic plasticity in our models, as well as technical assumptions in derivations (the ARC is assumed to be a separable function of phase and amplitude, and we assumed small stimulation and *ω*_*f*_ *≫ ω*_*s*_ or entrainment). We also used of a non-normalised MVL measure, although this is offset by the fact that stimulation waveform energy is constrained in numerical optimisations. Numerical optimisations were limited by available supercomputing resources, and the optimal balance of stimulation waveform Fourier coefficients across model parameters could not be investigated with a finer Fourier amplitude grid in the SL model, or at all in the WC model. Lastly, our framework assumes a smooth stimulation waveform (as in tACS). While we propose different ways of approximating the smooth optimal waveforms with pulsatile waveforms for neuromodulation devices that can only generate pulses (section 2.4), the pulse frequencies and durations required may not be achievable by some of these devices.

## Conclusion

We have presented a framework to design optimal PAC-enhancing (or PAC-decreasing) stimulation based on the amplitude response of the fast population, assuming that stimulation acts solely on the neural population generating the fast rhythm. We hope that this framework can help guide the development of innovative therapeutic brain stimulation aiming of restoring healthy levels of PAC, for example in patients with AD, where theta-gamma PAC is abnormally low and correlates with cognitive symptoms.

## 4 Appendix

In this section, we provide methodological details pertaining to PAC measurement, numerical optimisation of the stimulation waveform to enhance PAC, the perturbation analysis of stimulation Fourier coefficients, the WC model, and the estimation of ARCs in the WC model. We also present derivation details for the analytical approaches pursued in the SL model with direct stimulation coupling and general stimulation coupling.

### 4.1 Measuring PAC levels in simulations

In our simulations, the MVL is obtained in discrete time as

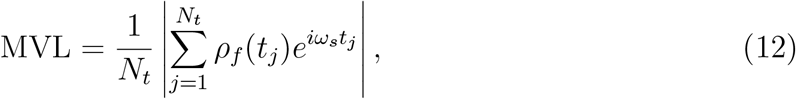

where *N*_*t*_ is the number of sampling points in the time period considered. The amplitude *ρ*_*f*_ is the Hilbert amplitude of the filtered model output (model output taken as ℜ(*z*_*f*_) for SL [42], and as *E* for WC). A third-order butterworth filter with zero-phase filtering is used to avoid phase distortions, and the Hilbert amplitude is given by the modulus of the analytic signal constructed with the Hilbert transform. It was also found empirically that stimulation waveform optimisations become unstable for narrow filter half-widths such as Δ*f*_*f*_ = 5 Hz. We therefore use filter half-widths of 10 or 20 Hz (see corresponding figure captions for specific values). Note that the phase of the slow signal is given in equation (12) by *ω*_*s*_*t*_*i*_ since in the examples considered, the slow signal is always cos(*ω*_*s*_*t*_*i*_), or a scaled and shifted version of it.

### 4.2 Numerical optimisation of stimulation waveform

To test our theory in the SL and WC models, we numerically optimise the Fourier coefficients of the stimulation waveform to maximise PAC. Each optimisation on a given variant of the SL or WC model consists of many local optimisations starting from different initial values of the stimulation Fourier coefficients. These are drawn from a uniform distribution on a logarithmic scale between 10^−3^ and 1, and then rescaled so that the energy of each initial waveform is the target stimulation energy Ξ. Local optimisations are performed in Matlab using the non-linear optimiser fmincon based on the interior-point algorithm [66], under the constraint that the energy of the waveform (given by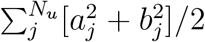) is kept within a small tolerance (*ϵ* = 0.1) of the target waveform energy Ξ. The target energy is chosen for each model variant so that the optimal stimulation has a significant effect on PAC (see corresponding figure captions for the specific target values). Hard bounds between 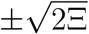 are also enforced during optimisation. For optimisation speed and accuracy, the models are simulated using Matlab’s solver ode113 (variable-step, variable-order Adams-Bashforth-Moulton solver of orders 1 to 13). At each optimisation step, models are simulated for 30s for variants of the SL model, and 20s for variants of the WC model (reduced duration to improve optimisation speed). The transient is discarded by removing the first third of the simulation output.

The objective function to minimise during optimisation is

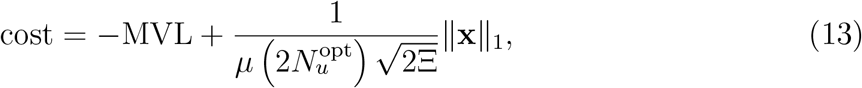

where the first term ensures that the level of PAC is maximised (MVL obtained as described in section 4.1), and the second term is a regularisation term. The Fourier coefficients being optimised are denoted 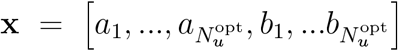, and the norm ∥**x**∥_1_ = Σ _*i*_ |*x*_*i*_| denotes the 1-norm. The regularisation term is scaled using the absolute energy bound for Fourier coefficients during the optimisation 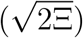, the number of Fourier coefficients being optimised 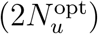, and a regularisation parameter *μ*. Regularisation was only used to guide the more challenging optimisations — we used *μ* = 1 for the variant of the SL model with dependence on *ρ, μ* = 15 for the WC models, and a very large number otherwise (no regularisation).

The number of local optimisations was increased when optimising all Fourier coefficients compared to when optimising only the Fourier coefficients predicted by theory. At least 3000 local optimisations were performed in the latter case, while at least 11000 local optimisations were performed in the former case. This was to mitigate the increase in the number of optimised parameters (within the limits of supercomputing resources available). In both cases, the best-ranked parameters coming out of the local optimisations were put through one other round of local optimisation (except for the SL models with mean-field and direct stimulation coupling, which were easier to optimise).

### 4.3 Perturbation analysis

As an additional investigation into which Fourier coefficients of the stimulation waveform are key to enhancing PAC, we perturb individual Fourier coefficients and assess changes in MVL. For each of the model variants considered, we perturb the *n*_pert_-best PAC-enhancing waveforms (obtained from numerically optimising all the Fourier coefficients), as well as *n*_pert_ random waveforms. Random waveforms are generated by drawing Fourier coefficients from a uniform distribution and re-scaling the coefficients such that the waveform energy is Ξ. We take *n*_pert_ = 100 for SL models, and *n*_pert_ = 200 for WC models (more variability in the latter case). We perturb each Fourier coefficient in turn by adding the perturbation 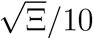, where Ξ is the waveform energy before perturbation. We measure the absolute change in PAC as |MVL − MVL_0_|, where MVL is the PAC level with the perturbation, and MVL_0_ the PAC level in the absence of perturbation. For each of the model variants considered, the absolute change in PAC is averaged separately across the *n*_pert_-best PAC-enhancing waveforms and the *n*_pert_ random waveforms.

We also follow the approach above to show that *φ*_*u*_ does not introduce significant dependences on other Fourier coefficients than those predicted by theory in the SL model. The only difference is that we fit a straight line to the time evolution of *θ*_*f*_ (first third of the data discarded to remove transient), and measure *φ*_*u*_ as its intercept. For each perturbation, we measure the absolute change in *φ*_*u*_ as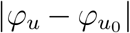, where 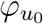 is the phase shift in the absence of perturbation. We average absolute differences as in the MVL case above.

### 4.4 Derivation details for foundational case two

In this section, we derive a relationship between stimulation waveform Fourier coefficients and the amplitude of the fast population in the case of direct coupling in the SL model (foundational case two). This will allow us to gain insight into which Fourier coefficients can have a significant impact on PAC. Using the approximation for *θ*_*f*_ mentioned in section 2.1.2 in the Results, the time evolution of *ρ*_*f*_ is given by equation (10). In the steady-state, solutions with PAC will be periodic with period 2*π*/*ω*_*s*_. Such solutions can therefore be approximated as Fourier series 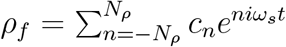 truncated at order *N*_*ρ*_. We also have 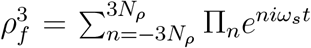, where the Fourier coefficients Π_*n*_ can be obtained as functions of the coefficients of *ρ*. Equation (10) becomes

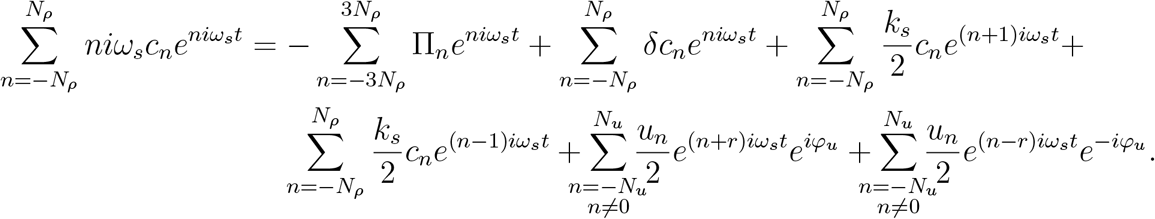

By manipulating indices and identifying terms corresponding to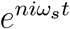, we obtain

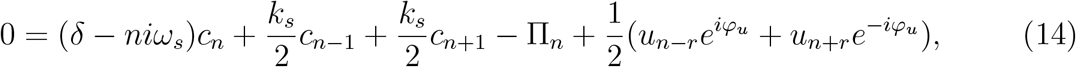

with *u*_0_ = 0, *u*_*n*_ = 0 for |*n*| > *N*_*u*_, *c*_*n*_ = 0 for |*n*| > *N*_*ρ*_, and Π_*n*_ = 0 for |*n*| > 3*N*_*ρ*_.

Most of the PAC strength is captured by the first harmonic of *ρ*, we therefore consider equation (14) for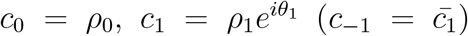, and *c*_*n*_ = 0 for |*n*| > 1. Since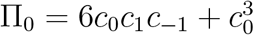, and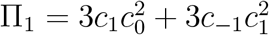, we get

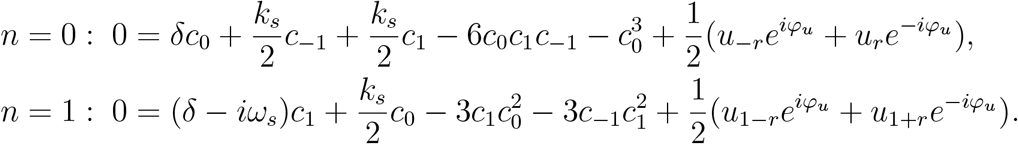

These two equations translate to three equations in *ρ*_0_, *ρ*_1_, and *θ*_1_ given by

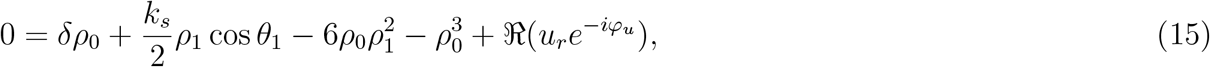

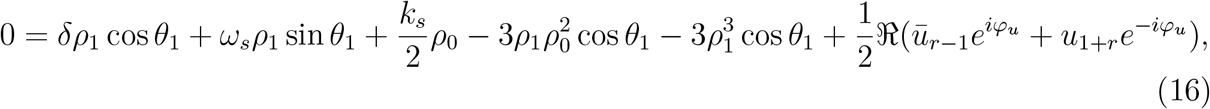

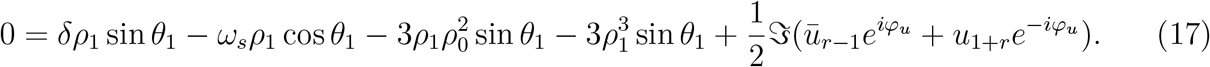

We demonstrate numerically through a perturbation approach that the phase shift *φ*_*u*_ (which also depends on the stimulation waveform) does not introduce dependences on additional Fourier coefficients than those explicitly present in these equations (see figure S.1A in Supplementary Material and methodological details in section 4.3, perturbation of size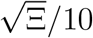). We present the insights obtained from these equations and test predictions arising from them in section 2.1.2 in the Results.

### 4.5 Derivation details for general stimulation coupling in the SL model

We generalise the derivation presented in the previous section to a general stimulation coupling, where the amplitude response curve of the fast population is a separable function of *θ*_*f*_ and *ρ*_*f*_ (see section 2.1.3 in the Results), with a view to gain insight into which Fourier coefficients can have a significant impact on PAC. From equation (1) with general coupling, the time evolutions of *ρ*_*f*_ and *θ*_*f*_ are given by

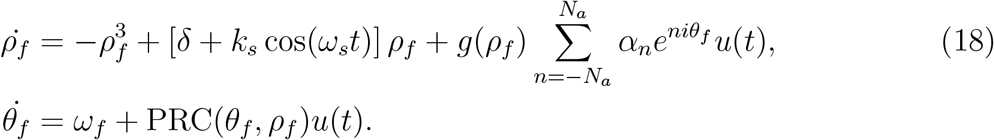

As previously, equation (18) can be approximated by

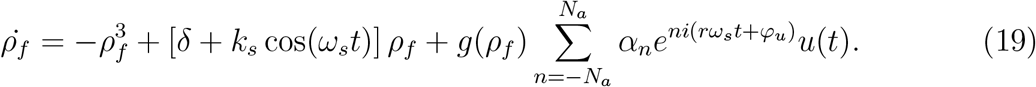

Since *ρ*_*f*_ is periodic, *g*(*ρ*_*f*_) can be approximated by a truncated Fourier series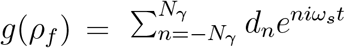. Note that each *d*_*n*_ depends on the Fourier coefficients of *ρ*_*f*_. Using the Fourier expansions of the various terms as before, equation (19) becomes

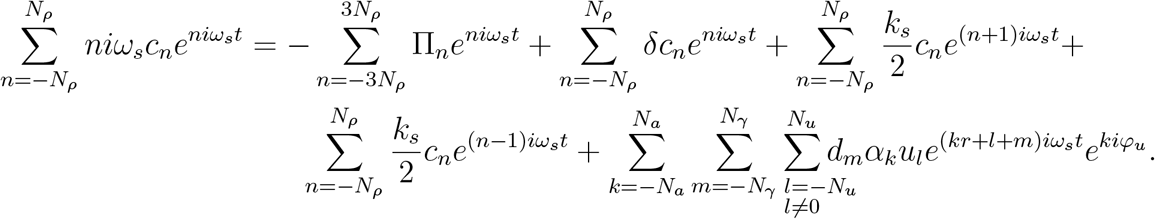

Using *n* = *kr* + *l* + *m*, we have

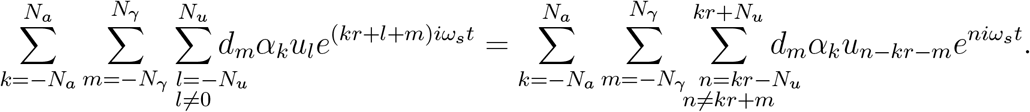

By manipulating indices and identifying terms corresponding to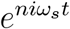, we obtain

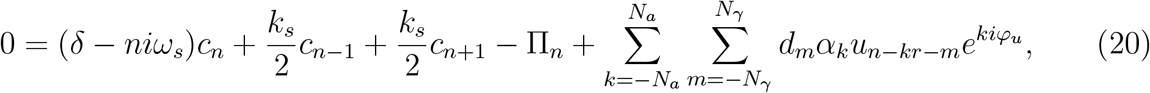

with *u*_0_ = 0, *u*_*n*_ = 0 for |*n*| > *N*_*u*_, *c*_*n*_ = 0 for |*n*| > *N*_*ρ*_, Π_*n*_ = 0 for |*n*| > 3*N*_*ρ*_, *a*_*n*_ = 0 for |*n*| > *N*_*a*_, and *d*_*n*_ = 0 for |*n*| > *N*_*γ*_.

As before, most of the PAC strength is captured by the first harmonic of *ρ* with coefficients *c*_0_ = *ρ*_0_,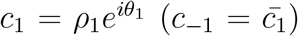. Neglecting the higher order harmonics of *ρ*, the coefficients *d*_*m*_ will only depend on *c*_0_ and *c*_1_. We have

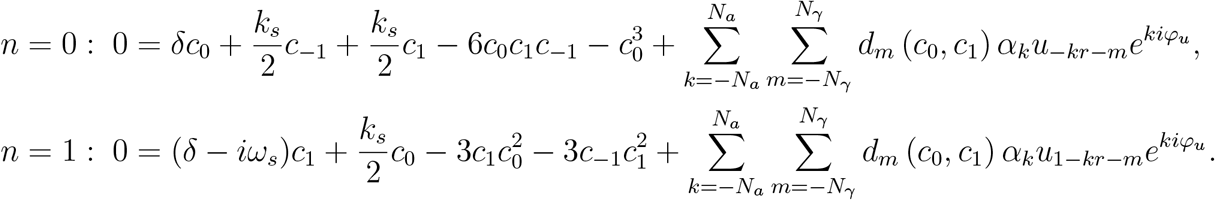

These two equations translate to three equations in *ρ*_0_, *ρ*_1_, and *θ*_1_ given by

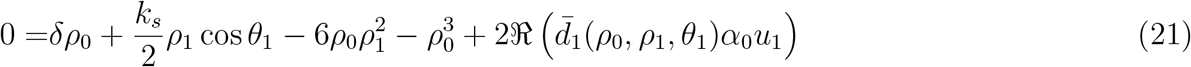

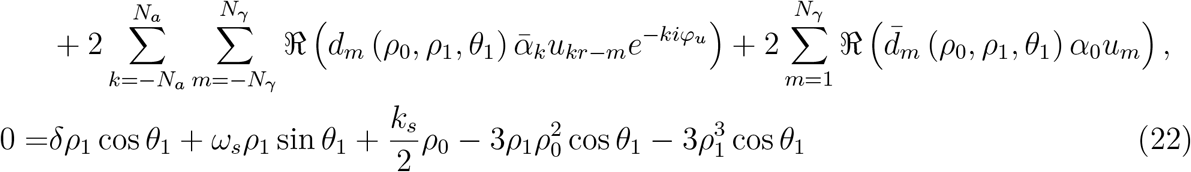

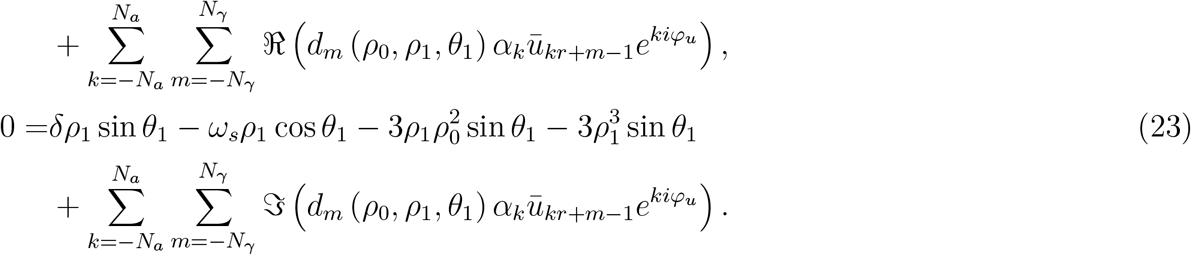

As before, we demonstrate numerically through a perturbation approach detailed in section 4.3 that the phase shift *φ*_*u*_ (which also depends on the stimulation waveform) does not introduce dependences on additional Fourier coefficients than those explicitly present in these equations (see figure S.1C for *g*(*ρ*_*f*_) = 1, and figure S.1D for *g*(*ρ*_*f*_) = 1/*ρ*_*f*_ in Supplementary Material). We present the insights obtained from these equations and test predictions arising from them for two examples of *g*(*ρ*_*f*_) in section 2.1.3 in the Results.

### 4.6 Wilson-Cowan model

To test whether the predictions obtained from the SL model may apply in a more biologically realistic context, we make use of a neural mass model, the Wilson-Cowan model. The WC model depicts the interactions of a population of excitatory neurons, whose activity is denoted by *E*, and a population of inhibitory neurons, whose activity is denoted by *I*(see figure 6). Two heuristically derived mean-field equations [43] describe the time evolution of the populations’ activities as

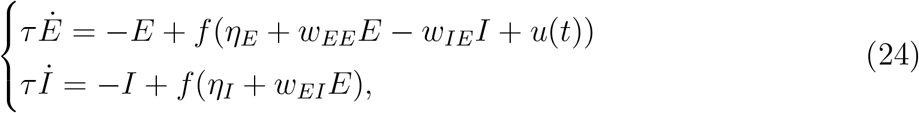

with *w*_*PR*_ the weight of the projection from population “P” to population “R”, *η*_*P*_ the external input to population “P”, *u*(*t*) the external stimulation, and *τ* a time constant (assumed to be the same for both populations). As in [44], the function *f* is the sigmoid function

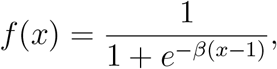

parametrised by a steepness parameter *β*. To get PAC in the absence of stimulation, we follow [44] and provide the slow input

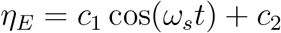

to the excitatory population (*c*_1_ is set to zero in the pure gamma case). We consider two examples with model parameters leading to dynamically distinct behaviours (strong theta case and pure gamma case, see section 2.2). The parameters used in the simulations are reported in Table 1.

### 4.7 Obtaining amplitude-response curves in the Wilson-Cowan model

Assessing whether predictions made with the SL model may hold for the WC model requires the ARC of the WC model in the examples considered. Thus, we approximate the ARC of the excitatory population (the population receiving stimulation) in the strong theta and pure gamma case as follows. Our approach is inspired by [67], and does not rely on the more complicated definitions of the amplitude response involving isostables [68, 69, 70, 71, 59]. The intuition behind our approach is as follows. The instantaneous change in the system’s state due to stimulation will in general change both the phase and the amplitude of the system. In the two-dimensional (*E, I*) phase space, the instantaneous change in phase due to a small stimulation at a given point on a trajectory can be obtained from the component of the shift due to stimulation that is tangent to the trajectory at the stimulation point. Conversely, the instantaneous change in amplitude is given by the component of the shift due to stimulation that is normal to the trajectory at the stimulation point. To obtain the ARC for a periodic trajectory of interest, we therefore need to compute the normal component of the change in state due stimulation at a number of points along the trajectory. These points are chosen such that they span the full range of phases on the periodic trajectory and capture the phase dependence of the ARC with sufficient detail (for the amplitude of the periodic orbit considered).

Along the periodic trajectories of interest, we therefore calculate at regular time intervals the instantaneous change in the activity of the E population due to stimulation Δ*E*_*u*_. As per equation (24), during a time step Δ*t*, the instantaneous change in *E* due to both stimulation and the dynamics of the system is given

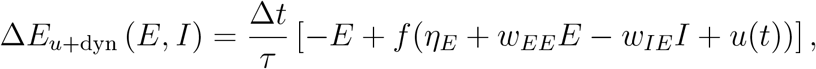

while the instantaneous change in *E* due to the dynamics only is given

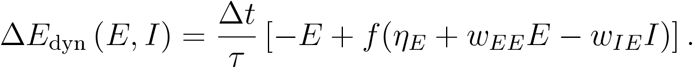

Thus, we obtain the instantaneous change in the activity of the E population due to stimulation as

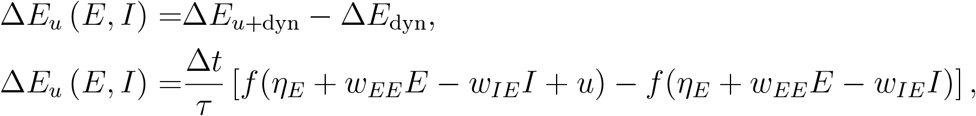

where we choose *u* = 0.3 in our numerical estimations of the ARC. Since stimulation is only provided to the excitatory population (see equation (24)), the instantaneous change in the activity of the I population due to stimulation is Δ*I*_*u*_ = 0. At each point considered along the trajectory, we also get the tangent vector to the trajectory as a numerical approximation of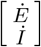) using central differences. We then obtain **n** as the counter-clockwise unit normal vector to the tangent vector. Finally, we approximate the ARC as the projection of Δ*E*_*u*_ onto the normal vector at each point considered along the trajectory of interest,

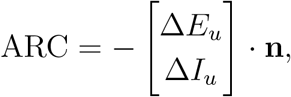

where the negative sign gives a positive value for an increase in amplitude. Each point along the trajectory where the change in amplitude was computed is assigned a phase given by *θ*_*f*_ = *ω*_*f*_ *t*, which allows us to re-parametrise the ARC as a function of phase. This ARC approximation process is illustrated in figure S.6 in the Supplementary Material. In the strong theta case, the trajectory considered is the on-stimulation gamma cycle (see figure S.6A). In the pure gamma case, the significant changes in dynamics for low and high amplitudes require to consider both the low-amplitude trajectory on stimulation (figure S.6B1), and the high-amplitude periodic trajectory (similar on-and off-stimulation, taken off-stimulation in figure S.6B2).

## Acknowledgements

We would like to acknowledge the use of the University of Oxford Advanced Research Computing (ARC) facility in carrying out this work http://dx.doi.org/10.5281/zenodo.22558

## Data availability statement

No data were collected as part of this work.

## References

[1] Florian Mormann, Juergen Fell, Nikolai Axmacher, Bernd Weber, Klaus Lehnertz, Christian E. Elger, and Guillén Fernández. Phase/amplitude reset and theta-gamma interaction in the human medial temporal lobe during a continuous word recognition memory task. Hippocampus, 15(7):890–900, jan 2005. doi:10.1002/hipo.20117.

[2] Adriano B.L. Tort, Robert W. Komorowski, Joseph R. Manns, Nancy J. Kopell, and Howard Eichenbaum. Theta-gamma coupling increases during the learning of item-context associations. Proceedings of the National Academy of Sciences of the United States of America, 106(49):20942–20947, dec 2009. doi:10.1073/pnas.0911331106.

[3] Ryan T Canolty and Robert T Knight. The functional role of cross-frequency coupling. Trends in Cognitive Sciences, 14(11):506–515, 2010. doi:10.1016/j.tics.2010.09.001.

[4] John E. Lisman and Ole Jensen. The Theta-Gamma Neural Code. Neuron, 77(6):1002–1016, 2013. doi:10.1016/j.neuron.2013.03.007.

[5] Kei M. Igarashi, Li Lu, Laura L. Colgin, May Britt Moser, and Edvard I. Moser. Coordination of entorhinal-hippocampal ensemble activity during associative learning. Nature, 510(7503):143–147, apr 2014. doi:10.1038/nature13162.

[6] Ludovico Saint Amour di Chanaz, Alexis Pérez-Bellido, Xiongbo Wu, Diego Lozano-Soldevilla, Daniel Pacheco-Estefan, Katia Lehongre, Estefanía Conde-Blanco, Pedro Roldan, Claude Adam, Virginie Lambrecq, Valerio Frazzini, Antonio Donaire, Mar Carreño, Vincent Navarro, Antoni Valero-Cabré, and Lluís Fuentemilla. Gamma amplitude is coupled to opposed hippocampal theta-phase states during the encoding and retrieval of episodic memories in humans. Current Biology, 33(9):1836–1843.e6, 2023. doi:10.1016/j.cub.2023.03.073.

[7] Jeffrey Z Nie, Robert D Flint, Prashanth Prakash, Jason K Hsieh, Emily M Mugler, Matthew C Tate, Joshua M Rosenow, and Marc W Slutzky. High-gamma activity is coupled to low-gamma oscillations in precentral cortices and modulates with movement and speech. bioRxiv, page 2023.02.13.528325, jan 2023. doi:10.1101/2023.02.13.528325.

[8] Georgios Spyropoulos, Conrado Arturo Bosman, and Pascal Fries. A theta rhythm in macaque visual cortex and its attentional modulation. Proceedings of the National Academy of Sciences, 115(24):E5614–E5623, jun 2018. doi:10.1073/pnas.1719433115.

[9] Yukiko Kikuchi, Adam Attaheri, Benjamin Wilson, Ariane E Rhone, Kirill V Nourski, Phillip E Gander, Christopher K Kovach, Hiroto Kawasaki, Timothy D Griffiths, Matthew A Howard III, and Christopher I Petkov. Sequence learning modulates neural responses and oscillatory coupling in human and monkey auditory cortex. PLOS Biology, 15(4):e2000219, apr 2017. doi:10.1371/journal.pbio.2000219.

[10] Bradley Voytek, Andrew S Kayser, David Badre, David Fegen, Edward F Chang, Nathan E Crone, Josef Parvizi, Robert T Knight, and Mark D’Esposito. Oscillatory dynamics coordinating human frontal networks in support of goal maintenance. Nature Neuroscience, 18(9):1318–1324, 2015. doi:10.1038/nn.4071.

[11] Kazuki Sakakura, Naoto Kuroda, Masaki Sonoda, Takumi Mitsuhashi, Ethan Firestone, Aimee F Luat, Neena I Marupudi, Sandeep Sood, and Eishi Asano. Developmental atlas of phase-amplitude coupling between physiologic high-frequency oscillations and slow waves. Nature Communications, 14(1):6435, 2023. doi:10.1038/s41467-023-42091-y.

[12] Yousef Salimpour and William S Anderson. Cross-Frequency Coupling Based Neuromodulation for Treating Neurological Disorders, 2019. doi:10.3389/fnins.2019.00125.

[13] Coralie De Hemptinne, Elena S. Ryapolova-Webb, Ellen L. Air, Paul A. Garcia, Kai J. Miller, Jeffrey G. Ojemann, Jill L. Ostrem, Nicholas B. Galifianakis, and Philip A. Starr. Exaggerated phase-amplitude coupling in the primary motor cortex in Parkinson disease. Proceedings of the National Academy of Sciences of the United States of America, 110(12):4780–4785, mar 2013. doi:10.1073/pnas.1214546110.

[14] Nicole C. Swann, Coralie De Hemptinne, Adam R. Aron, Jill L. Ostrem, Robert T. Knight, and Philip A. Starr. Elevated synchrony in Parkinson disease detected with electroencephalography. Annals of Neurology, 78(5):742–750, nov 2015. doi:10.1002/ana.24507.

[15] Coralie De Hemptinne, Nicole C. Swann, Jill L. Ostrem, Elena S. Ryapolova-Webb, Marta San Luciano, Nicholas B. Galifianakis, and Philip A. Starr. Therapeutic deep brain stimulation reduces cortical phase-amplitude coupling in Parkinson’s disease. Nature Neuroscience, 18(5):779–786, apr 2015. doi:10.1038/nn.3997.

[16] Efstathios D Kondylis, Michael J Randazzo, Ahmad Alhourani, Witold J Lipski, Thomas A Wozny, Yash Pandya, Avniel S Ghuman, Robert S Turner, Donald J Crammond, and R Mark Richardson. Movement-related dynamics of cortical oscillations in Parkinson’s disease and essential tremor. Brain, 139(8):2211–2223, aug 2016. doi:10.1093/brain/aww144.

[17] Guillaume Etter, Suzanne van der Veldt, Frédéric Manseau, Iman Zarrinkoub, Emilie Trillaud-Doppia, and Sylvain Williams. Optogenetic gamma stimulation rescues memory impairments in an Alzheimer’s disease mouse model. Nature Communications, 10(1), 2019. doi:10.1038/s41467-019-13260-9.

[18] Paolo Bazzigaluppi, Tina L. Beckett, Margaret M. Koletar, Aaron Y. Lai, Illsung L. Joo, Mary E. Brown, Peter L. Carlen, Jo Anne McLaurin, and Bojana Stefanovic. Early-stage attenuation of phase-amplitude coupling in the hippocampus and medial prefrontal cortex in a transgenic rat model of Alzheimer’s disease. Journal of Neurochemistry, 144(5):669–679, mar 2018. doi:10.1111/jnc.14136.

[19] Romain Goutagny, Ning Gu, Chelsea Cavanagh, Jesse Jackson, Jean Guy Chabot, Rémi Quirion, Slavica Krantic, and Sylvain Williams. Alterations in hippocampal network oscillations and theta-gamma coupling arise before Aβ overproduction in a mouse model of Alzheimer’s disease. European Journal of Neuroscience, 37(12):1896– 1902, jun 2013. doi:10.1111/ejn.12233.

[20] Michelle S. Goodman, Sanjeev Kumar, Reza Zomorrodi, Zaid Ghazala, Amay S.M. Cheam, Mera S. Barr, Zafiris J. Daskalakis, Daniel M. Blumberger, Corinne Fischer, Alastair Flint, Linda Mah, Nathan Herrmann, Christopher R. Bowie, Benoit H. Mulsant, Tarek K. Rajji, Bruce G. Pollock, Lillian Lourenco, Meryl Butters, Damian Gallagher, Angela Golas, Ariel Graff, James L. Kennedy, Shima Ovaysikia, Mark Rapoport, Kevin Thorpe, Nicolaas P.L.G. Verhoeff, and Aristotle N. Voineskos. Theta-Gamma coupling and working memory in Alzheimer’s dementia and mild cognitive impairment. Frontiers in Aging Neuroscience, 10(APR):101, apr 2018. doi:10.3389/fnagi.2018.00101.

[21] Christian Sandøe Musaeus, Malene Schjønning Nielsen, Jørgen Sandøe Musaeus, and Peter Høgh. Electroencephalographic Cross-Frequency Coupling as a Sign of Disease Progression in Patients With Mild Cognitive Impairment: A Pilot Study. Frontiers in Neuroscience, 14:790, aug 2020. doi:10.3389/fnins.2020.00790.

[22] Ruihua Zhang, Ye Ren, Chunyan Liu, Na Xu, Xiaoli Li, Fengyu Cong, Tapani Ristaniemi, and Yu Ping Wang. Temporal-spatial characteristics of phase-amplitude coupling in electrocorticogram for human temporal lobe epilepsy. Clinical Neurophysiology, 128(9):1707–1718, sep 2017. doi:10.1016/j.clinph.2017.05.020.

[23] Nabi Rustamov, Joseph Humphries, Alexandre Carter, and Eric C. Leuthardt. Theta-gamma coupling as a cortical biomarker of brain-computer interface-mediated motor recovery in chronic stroke. Brain Communications, 4(3), may 2022. doi:10.1093/braincomms/fcac136.

[24] Songjian Wang, Chunlin Li, Yi Liu, Mengyue Wang, Meng Lin, Liu Yang, Younuo Chen, Yuan Wang, Xinxing Fu, Xu Zhang, and Shuo Wang. Features of beta-gamma phase-amplitude coupling in cochlear implant users derived from EEG. Hearing Research, 428:108668, 2023. doi:10.1016/j.heares.2022.108668.

[25] Yousef Salimpour, Kelly A. Mills, Brian Y. Hwang, and William S. Anderson. Phase-targeted stimulation modulates phase-amplitude coupling in the motor cortex of the human brain. Brain Stimulation, 15(1):152–163, jan 2022. doi:10.1016/j.brs.2021.11.019.

[26] Zhenyu Xie, Shuxun Dong, Yiyao Zhang, and Yi Yuan. Transcranial ultrasound stimulation at the peak-phase of theta-cycles in the hippocampus improve memory performance. NeuroImage, 283:120423, dec 2023. doi:10.1016/j.neuroimage.2023.120423.

[27] Yuzhen Qin, Tommaso Menara, Danielle S Bassett, and Fabio Pasqualetti. Phase-amplitude coupling in neuronal oscillator networks. Physical Review Research, 3(2):23218, 2021. doi:10.1103/PhysRevResearch.3.023218.

[28] Joana Cabral, Francesca Castaldo, Jakub Vohryzek, Vladimir Litvak, Christian Bick, Renaud Lambiotte, Karl Friston, Morten L. Kringelbach, and Gustavo Deco. Metastable oscillatory modes emerge from synchronization in the brain space-time connectome. Communications Physics, 5(1):1–13, jul 2022. doi:10.1038/s42005-022-00950-y.

[29] Francesca Castaldo, Francisco Páscoa dos Santos, Ryan C Timms, Joana Cabral, Jakub Vohryzek, Gustavo Deco, Mark Woolrich, Karl Friston, Paul Verschure, and Vladimir Litvak. Multi-modal and multi-model interrogation of large-scale functional brain networks. NeuroImage, 277:120236, 2023. doi:10.1016/j.neuroimage.2023.120236.

[30] Christoffer G. Alexandersen, Willem de Haan, Christian Bick, and Alain Goriely. A multi-scale model explains oscillatory slowing and neuronal hyperactivity in Alzheimer’s disease. Journal of the Royal Society Interface, 20(198), jan 2023. doi:10.1098/rsif.2022.0607.

[31] György Buzsáki. Theta Oscillations in the Hippocampus. Neuron, 33(3):325–340, jan 2002. doi:10.1016/S0896-6273(02)00586-X.

[32] Balázs Hangya, Zsolt Borhegyi, Nóra Szilágyi, Tamás F Freund, and Viktor Varga. GABAergic Neurons of the Medial Septum Lead the Hippocampal Network during Theta Activity. The Journal of Neuroscience, 29(25):8094 LP –8102, jun 2009. doi:10.1523/JNEUROSCI.5665-08.2009.

[33] Laura Lee Colgin. Mechanisms and Functions of Theta Rhythms. Annual Review of Neuroscience, 36(1):295–312, jul 2013. doi:10.1146/annurev-neuro-062012-170330.

[34] Abbey B Holt, Eszter Kormann, Alessandro Gulberti, Monika Pötter-Nerger, Colin G McNamara, Hayriye Cagnan, Magdalena K Baaske, Simon Little, Johannes A Köppen, and Carsten Buhmann. Phase-dependent suppression of beta oscillations in Parkinson’s disease patients. Journal of Neuroscience, 39(6):1119–1134, 2019. doi:10.1523/JNEUROSCI.1913-18.2018.

[35] Colin G. McNamara, Max Rothwell, and Andrew Sharott. Stable, interactive modulation of neuronal oscillations produced through brain-machine equilibrium. Cell Reports, 41(6):111616, nov 2022. doi:10.1016/j.celrep.2022.111616.

[36] D Wilson and J Moehlis. Determining individual phase response curves from aggregate population data. Phys Rev E Stat Nonlin Soft Matter Phys, 92(2):22902, 2015. doi:10.1103/PhysRevE.92.022902.

[37] Bharat Monga and Jeff Moehlis. Phase distribution control of a population of oscillators. Physica D: Nonlinear Phenomena, 398:115–129, nov 2019. 1811.10562, doi:10.1016/j.physd.2019.06.001.

[38] Benoit Duchet, James J Sermon, Gihan Weerasinghe, Timothy Denison, and Rafal Bogacz. How to entrain a selected neuronal rhythm but not others: open-loop dithered brain stimulation for selective entrainment. Journal of Neural Engineering, 20(2):026003, apr 2023. doi:10.1088/1741-2552/acbc4a.

[39] R T Canolty, E Edwards, S S Dalal, M Soltani, S S Nagarajan, H E Kirsch, M S Berger, N M Barbaro, and R T Knight. High Gamma Power Is Phase-Locked to Theta Oscillations in Human Neocortex. Science, 313(5793):1626–1628, sep 2006. doi:10.1126/science.1128115.

[40] Mareike J. Hülsemann, Ewald Naumann, and Björn Rasch. Quantification of phaseamplitude coupling in neuronal oscillations:comparison of phase-locking value, mean vector length, modulation index, and generalized-linear-modeling-cross-frequency-coupling. Frontiers in Neuroscience, 13(JUN):573, 2019. doi:10.3389/fnins.2019.00573.

[41] Ali Bakhshandeh Rostami. Exact Solution of Abel Differential Equation with Arbitrary Nonlinear Coefficients. arXiv, 2015. 1503.05929.

[42] Gihan Weerasinghe, Benoit Duchet, Hayriye Cagnan, Peter Brown, Christian Bick, and Rafal Bogacz. Predicting the effects of deep brain stimulation using a reduced coupled oscillator model. PLOS Computational Biology, 15(8):e1006575, aug 2019. doi:10.1371/journal.pcbi.1006575.

[43] Hugh R Wilson and Jack D Cowan. Excitatory and inhibitory interactions in localized populations of model neurons. Biophysical journal, 12(1):1–24, 1972. doi:10.1016/S0006-3495(72)86068-5.

[44] A C Onslow, M W Jones, and R Bogacz. A canonical circuit for generating phase-amplitude coupling. PloS one, 9(8):e102591, 2014. doi:10.1371/journal.pone.0102591.

[45] H Cagnan, D Pedrosa, S Little, A Pogosyan, B Cheeran, T Aziz, A Green, J Fitzgerald, T Foltynie, P Limousin, L Zrinzo, M Hariz, K J Friston, T Denison, and P Brown. Stimulating at the right time: phase-specific deep brain stimulation. Brain, 140(Pt 1):132–145, 2017. doi:10.1093/brain/aww286.

[46] N Kopell, C Börgers, D Pervouchine, P Malerba, and A Tort. Gamma and Theta Rhythms in Biophysical Models of Hippocampal Circuits BT - Hippocampal Micro-circuits: A Computational Modeler’s Resource Book. pages 423–457. Springer New York, New York, NY, 2010. doi:10.1007/978-1-4419-0996-1_15.

[47] Adam Ponzi, Salvador Dura-Bernal, and Michele Migliore. Theta-gamma phase amplitude coupling in a hippocampal CA1 microcircuit. PLoS Computational Biology, 19(3 March):e1010942, mar 2023. doi:10.1371/journal.pcbi.1010942.

[48] Roberto C Sotero. Topology, Cross-Frequency, and Same-Frequency Band Interactions Shape the Generation of Phase-Amplitude Coupling in a Neural Mass Model of a Cortical Column. PLOS Computational Biology, 12(11):e1005180, nov 2016. URL: 10.1371/journal.pcbi.1005180.

[49] Alexandre Hyafil, Anne Lise Giraud, Lorenzo Fontolan, and Boris Gutkin. Neural Cross-Frequency Coupling: Connecting Architectures, Mechanisms, and Functions, nov 2015. doi:10.1016/j.tins.2015.09.001.

[50] Osvaldo Matías Velarde, Eugenio Urdapilleta, Germán Mato, and Damián Dellavale. Bifurcation structure determines different phase-amplitude coupling patterns in the activity of biologically plausible neural networks. NeuroImage, 202:116031, nov 2019. doi:10.1016/j.neuroimage.2019.116031.

[51] Yousef Salimpour, Anish Nayak, Elizaveta Naydanova, Min Jae Kim, Brian Y. Hwang, Kelly A. Mills, Pawel Kudela, and William S. Anderson. Phase-dependent Stimulation for Modulating Phase-amplitude Coupling: A Computational Modeling Approach. In Proceedings of the Annual International Conference of the IEEE Engineering in Medicine and Biology Society, EMBS, volume 2020-July, pages 3590–3593. Institute of Electrical and Electronics Engineers Inc., jul 2020. doi:10.1109/EMBC44109.2020.9175966.

[52] Nikolaos Vardalakis, Amélie Aussel, Nicolas P Rougier, and Fabien B Wagner. A dynamical computational model of theta generation in hippocampal circuits to study theta-gamma oscillations during neurostimulation. Elife, 12:2003–2023, 2023. doi:10.7554/ELIFE.87356.

[53] A Gillies, D Willshaw, and Z Li. Subthalamic-pallidal interactions are critical in determining normal and abnormal functioning of the basal ganglia. Proc Biol Sci, 269(1491):545–551, 2002. doi:10.1098/rspb.2001.1817.

[54] A J Holgado, J R Terry, and R Bogacz. Conditions for the generation of beta oscillations in the subthalamic nucleus-globus pallidus network. J Neurosci, 30(37):12340– 12352, 2010. doi:10.1523/JNEUROSCI.0817-10.2010.

[55] Alejo J. Nevado-Holgado, Nicolas Mallet, Peter J. Magill, and Rafal Bogacz. Effective connectivity of the subthalamic nucleus-globus pallidus network during Parkinsonian oscillations. Journal of Physiology, 592(7):1429–1455, apr 2014. doi:10.1113/jphysiol.2013.259721.

[56] I Haidar, W Pasillas-Lepine, A Chaillet, E Panteley, S Palfi, and S Senova. Closed-loop firing rate regulation of two interacting excitatory and inhibitory neural populations of the basal ganglia. Biol Cybern, 110(1):55–71, 2016. doi:10.1007/s00422-015-0678-y.

[57] Benoit Duchet, Gihan Weerasinghe, Hayriye Cagnan, Peter Brown, Christian Bick, and Rafal Bogacz. Phase-dependence of response curves to deep brain stimulation and their relationship: from essential tremor patient data to a Wilson–Cowan model. The Journal of Mathematical Neuroscience, 10(1):4, 2020. doi:10.1186/s13408-020-00081-0.

[58] Benoit Duchet, Filippo Ghezzi, Gihan Weerasinghe, Gerd Tinkhauser, Andrea Kühn, Peter Brown, Christian Bick, and Rafal Bogacz. Average beta burst duration profiles provide a signature of dynamical changes between the ON and OFF medication states in Parkinson’s disease. PLOS Computational Biology, 17(7):e1009116, jul 2021. doi:10.1101/2020.04.27.064246.

[59] Benoit Duchet, Gihan Weerasinghe, Christian Bick, and Rafal Bogacz. Optimizing deep brain stimulation based on isostable amplitude in essential tremor patient models. Journal of Neural Engineering, 18(4):046023, mar 2021. doi:10.1088/1741-2552/abd90d.

[60] James J. Sermon, Maria Olaru, Juan Anso, Stephanie Cernera, Simon Little, Maria Shcherbakova, Rafal Bogacz, Philip A. Starr, Timothy Denison, and Benoit Duchet. Sub-harmonic entrainment of cortical gamma oscillations to deep brain stimulation in Parkinson’s disease: Model based predictions and validation in three human subjects. Brain Stimulation, 16(5):1412–1424, sep 2023. doi:10.1016/j.brs.2023.08.026.

[61] Ivan Alekseichuk, Zsolt Turi, Gabriel Amador de Lara, Andrea Antal, and Walter Paulus. Spatial Working Memory in Humans Depends on Theta and High Gamma Synchronization in the Prefrontal Cortex. Current Biology, 26(12):1513–1521, 2016. doi:10.1016/j.cub.2016.04.035.

[62] Haya Akkad, Joshua Dupont-Hadwen, Amba Frese, Irena Tetkovic, Liam Barrett, Sven Bestmann, and Charlotte J. Stagg. Increasing motor skill acquisition by driving theta-gamma coupling. eLife, 2021. doi:10.7554/eLife.67355.

[63] Justin Riddle, Amber McFerren, and Flavio Frohlich. Causal role of cross-frequency coupling in distinct components of cognitive control. Progress in Neurobiology, 202:102033, 2021. doi:10.1016/j.pneurobio.2021.102033.

[64] Colin Gerard McNamara and Andrew David Sharott. Apparatus and method for phase tracking an oscillatory signal, jun 2022.

[65] José Á Ochoa, Irene Gonzalez-Burgos, María J Nicolás, and Miguel Valencia. Open Hardware Implementation of Real-Time Phase and Amplitude Estimation for Neurophysiologic Signals, 2023. doi:10.3390/bioengineering10121350.

[66] Richard H. Byrd, Mary E. Hribar, and Jorge Nocedal. An interior point algorithm for large-scale nonlinear programming. SIAM Journal on Optimization, 9(4):877–900, jul 1999. doi:10.1137/S1052623497325107.

[67] K C Wedgwood, K K Lin, R Thul, and S Coombes. Phase-amplitude descriptions of neural oscillator models. J Math Neurosci, 3(1):2, 2013. doi:10.1186/2190-8567-3-2.

[68] A. Mauroy, I. Mezić, and J. Moehlis. Isostables, isochrons, and Koopman spectrum for the action-angle representation of stable fixed point dynamics. Physica D: Nonlinear Phenomena, 261:19–30, 2013. doi:10.1016/j.physd.2013.06.004.

[69] Dan Wilson and Jeff Moehlis. Isostable reduction of periodic orbits. Physical Review E, 94(5):52213, 2016. doi:10.1103/PhysRevE.94.052213.

[70] S Shirasaka, W Kurebayashi, and H Nakao. Phase-amplitude reduction of transient dynamics far from attractors for limit-cycling systems. Chaos, 27(2):23119, 2017. doi:10.1063/1.4977195.

[71] A. Mauroy and I. Mezić. Global computation of phase-amplitude reduction for limit-cycle dynamics. Chaos, 28(7):073108, jul 2018. doi:10.1063/1.5030175.

