## Supplementary Material for "How to design optimal brain stimulation to modulate phase-amplitude coupling?"

### A Exact semi-analytical approach for stimulation coupled through the mean-field in the Stuart-Landau model

The analytical approach presented in section 2.1.1 in the main text neglects the relaxation time to the limit cycle arising from equation (4). This can be avoided by noting that Equation (4) decouples in $\rho_f$ and $\theta_f$, yielding

$$\begin{aligned}\dot{\rho_f} &= -\rho_f^3 + \left[\delta + k_s \cos(\omega_s t) + u(t)\right]\rho_f,\\ \dot{\theta_f} &= \omega_f.\end{aligned} \tag{S.1}$$

Equation (S.1) is a Bernoulli differential equation and can solved by considering $h(t) = \rho_f(t)^{-2}$. We get

$$\rho_f(t) = \frac{e^{\delta t + \frac{k_s}{\omega_s}\sin(\omega_s t) + \sum_{n=1}^{N_u}\left\{\frac{a_n}{n\omega_s}\sin(n\omega_s t) - \frac{b_n}{n\omega_s}\cos(n\omega_s t)\right\}}}{\left[\frac{\rho_f(0)^2}{e^{2\sum_{n=1}^{N_u}\frac{b_n}{n\omega_s}}} + 2\int_0^t e^{2\left(\delta t_1 + \frac{k_s}{\omega_s}\sin(\omega_s t_1) + \sum_{n=1}^{N_u}\left\{\frac{a_n}{n\omega_s}\sin(n\omega_s t_1) - \frac{b_n}{n\omega_s}\cos(n\omega_s t_1)\right\}\right)} dt_1\right]^{\frac{1}{2}}}.$$

This expression for $\rho_f(t)$ can be plugged in equation (5), and the Fourier coefficients of $u(t)$ can be numerically optimised up to order $N_u$ to maximise the PAC metric $\Gamma$.

### B Supplementary figures

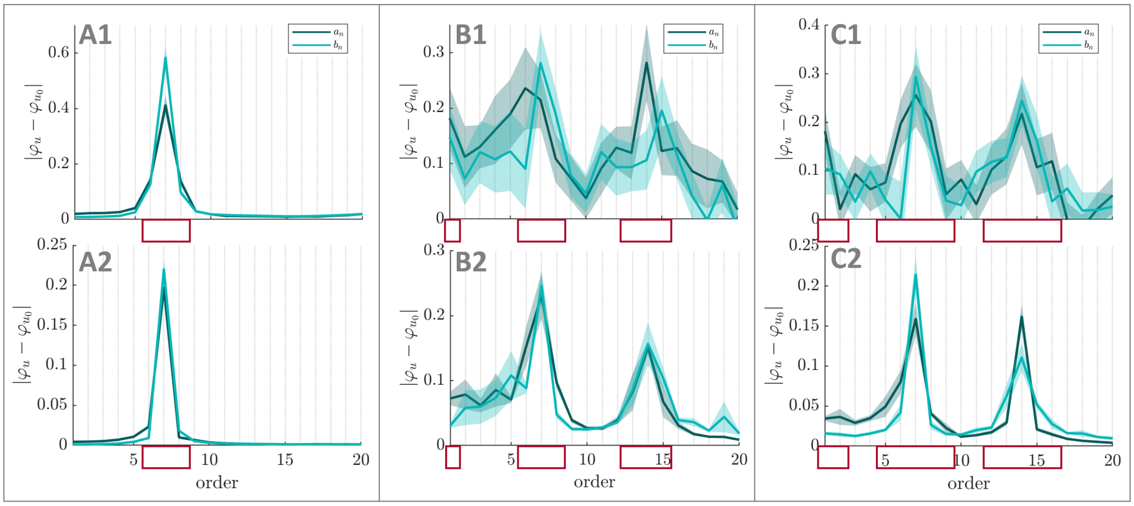


**Figure S.1: Dependence of $\varphi_u$ on Fourier coefficients of the stimulation waveform in the Stuart-Landau model.** The absolute change in $\varphi_u$ when increasing the energy of a given stimulation Fourier coefficient is provided in the first row when starting from PAC-enhancing waveforms obtained from the numerical optimisation process with all coefficients optimised, and in the second row when starting from random waveforms. The first column correspond to direct stimulation coupling (parameters from figure 4), the second column to the parameters from figure 5 ($g(\rho_f) = 1$), and the third column to the parameters from figure S.4 ($g(\rho_f) = 1/(\rho_f + 0.01)$). Error bars represent the standard error of the mean.

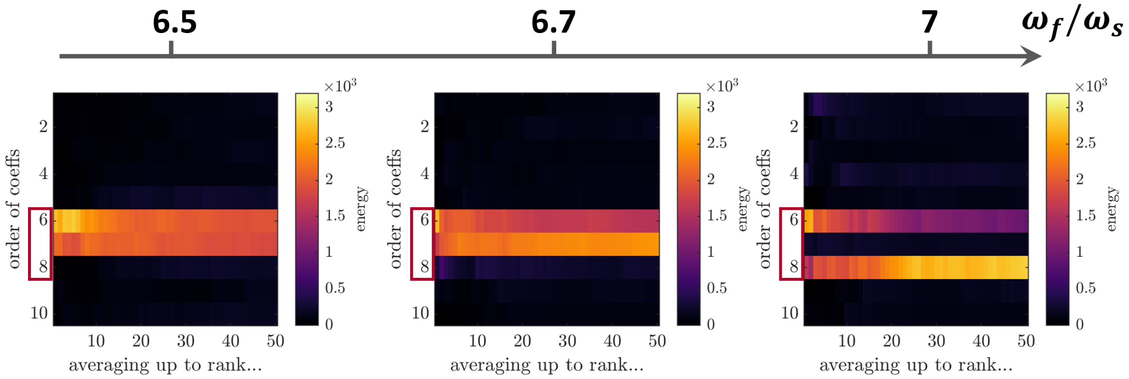


**Figure S.2: Influence of $\omega_f/\omega_s$ not being an integer on optimal PAC-enhancing waveforms in the Stuart-Landau model (case of direct stimulation coupling).** In all panels, the energy of PAC-enhancing waveforms obtained from numerical optimisation for all Fourier coefficient orders (vertical axis) when averaging the $x$-best optimisation results ($x$ being the horizontal axis value) is represented (all Fourier coefficients were optimised). As the ratio of the fast to slow frequencies deviates from an integer value, the optimal balance of energy between Fourier coefficients of the optimal waveform changes. However no Fourier coefficients other than those predicted by the theory (highlighted with red rectangles) have significant energy even when the ratio of the fast to slow frequencies is not an integer.

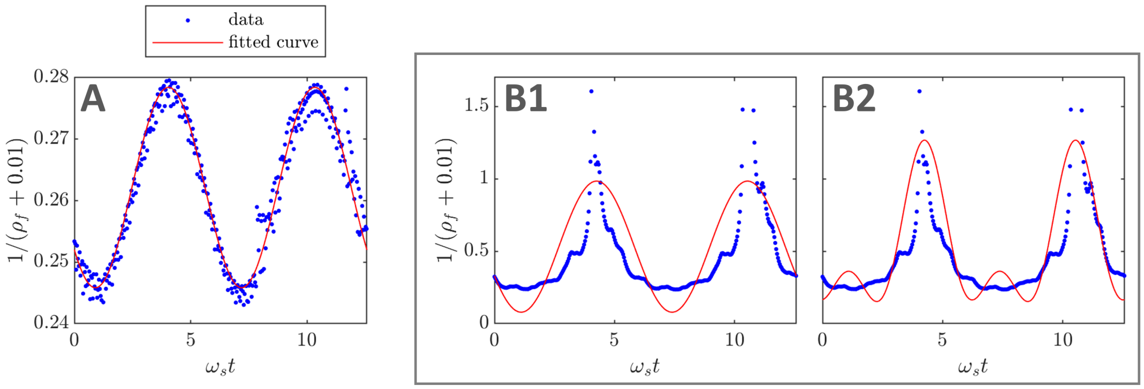


**Figure S.3: Approximating $\mathbf{g(\rho_f)}$ using truncated Fourier series.** A: $g(\rho_f)$ is well described off-stimulation by one harmonic (simulated system as in figure S.4A). B: $g(\rho_f)$ can be approximated on-stimulation by one harmonic as shown in B1. A better approximation is obtained with two harmonics in B2 (simulated system as in figure S.4B, using optimal PAC-enhancing waveform shown in figure S.4C). In all panels, data points obtained from model simulations are shown in blue, and the truncated Fourier series fit is shown in red.

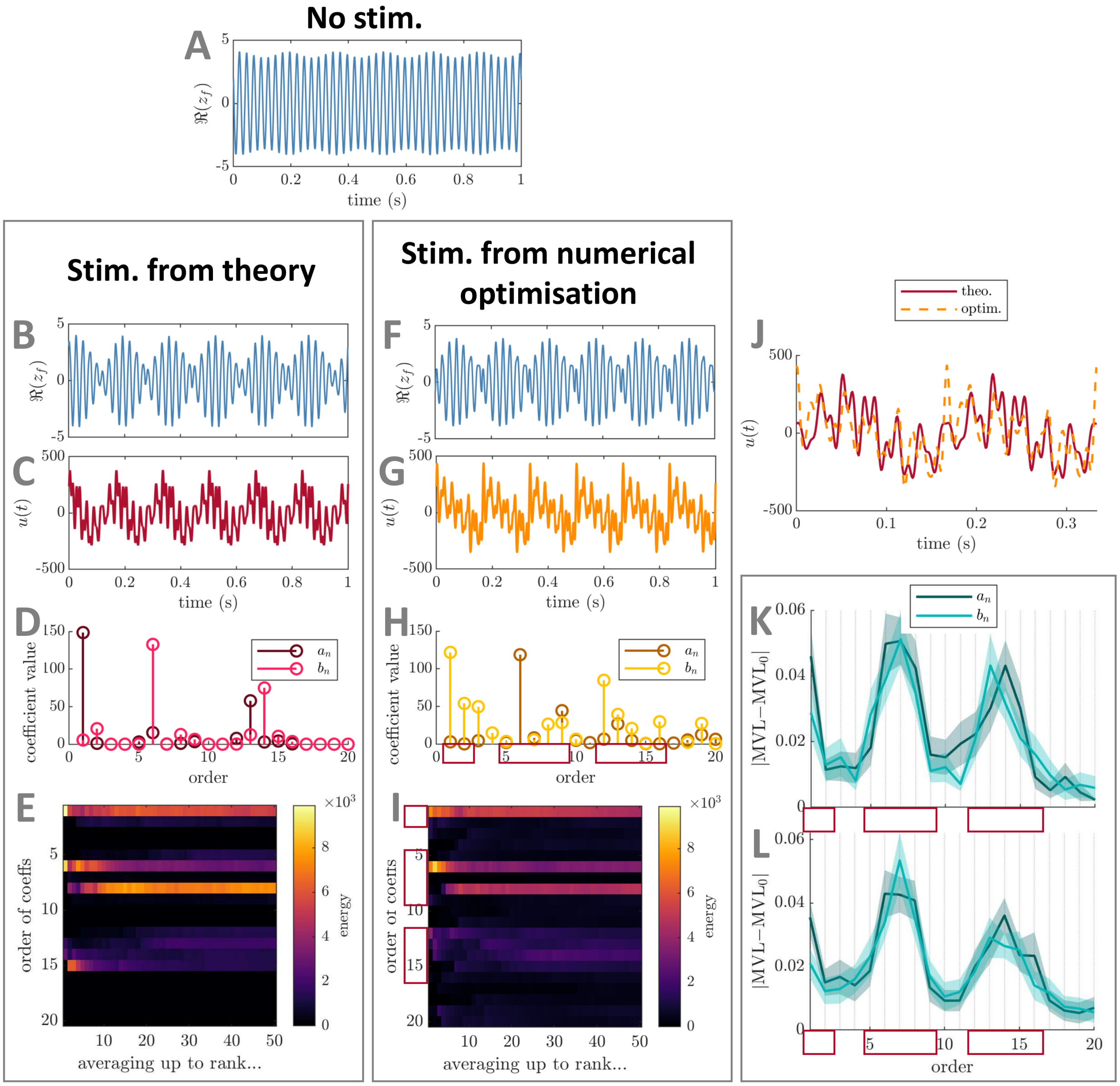


**Figure S.4: Comparison between best PAC-enhancing waveforms predicted by theory and by numerical optimisation – example for the general stimulation coupling case (with $\propto 1/\rho_f$ dependence) in the Stuart-Landau model.** The model output in the absence of stimulation is shown in panel A. The model output when receiving PAC-enhancing stimulation is shown in panels B (best stimulation waveform obtained when optimising only Fourier coefficients predicted by theory) and F (best stimulation waveform obtained when optimising all Fourier coefficients). The corresponding best PAC-enhancing stimulation waveforms are shown in panels C and G, respectively, and are overlaid for comparison in panel J (aligned to maximise their cross-correlation). Their Fourier coefficients are shown in panels D and H, respectively. The energy of PAC-enhancing waveforms obtained from numerical optimisation for all Fourier coefficient orders (vertical axis) when averaging the $x$-best optimisation results ($x$ being the horizontal axis value) is represented in panels E (only Fourier coefficients predicted by theory were optimised) and I (all Fourier coefficients were optimised). The absolute change in MVL when increasing the energy of a given stimulation Fourier coefficient is provided in panels K (when starting from PAC-enhancing waveforms obtained from the numerical optimisation process with all coefficients optimised), and L (when starting from random waveforms). Error bars represent the standard error of the mean. The Fourier coefficients predicted to be (potential) key contributors to PAC levels by theory are highlighted by red rectangles in panels H, I, K, and L. MVL for the stimulation waveform with only coefficients predicted by theory optimised is 0.502, MVL for the stimulation waveform with all coefficients optimised is 0.441, MVL in the absence of stimulation is 0.082 ($\Delta f_f = 20$ Hz). In all cases, waveform energy is fixed at $\Xi = 5000$. The parameters of the Stuart-Landau model used are $\delta = 15$, $k_s = 3$, $f_f = 42$ Hz, and $f_s = 6$ Hz ($r = 7$). Stimulation is acting through $\mathrm{PRC}(\theta_f, \rho_f) = [0.2 - \sin(\theta_f) + 0.7\cos(2\theta_f)]\,/(\rho_f{+}0.01)$ and $\mathrm{ARC}(\theta_f) = [0.4 + \cos(\theta_f) - 0.5\sin(2\theta_f)]\,/(\rho_f{+}0.01)$.

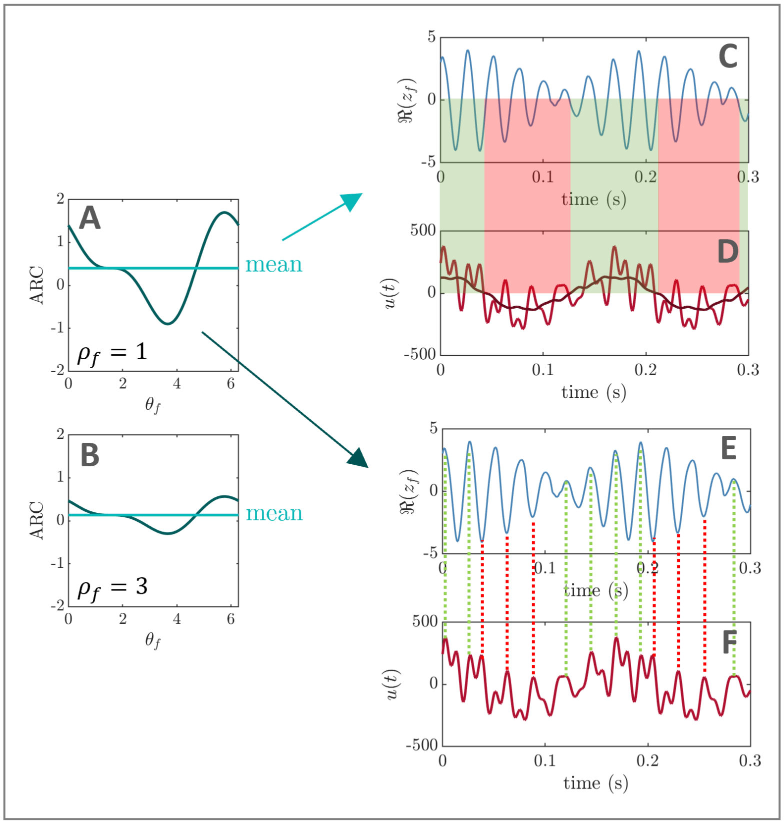


**Figure S.5: PAC-enhancing mechanisms in the Stuart-Landau model depend on the amplitude response of the fast population (case of a general stimulation coupling with dependence on $\rho_\mathbf{f}$).** General case where the amplitude response of the fast population depends on its phase and amplitude, and has a non-zero mean (highlighted in light green in A and B). Panel A shows the ARC of the fast population for $\rho_f = 1$, and panel B for $\rho_f = 3$. The optimal stimulation waveform (taken from figure S.4C) combines mechanisms of PAC-enhancement from foundational cases one and two (see figure 2A and B), as shown in panels C, D, E, and F. The dark red line in D represents a moving average of the optimal stimulation waveform (sliding window corresponding approximately to two fast-population cycles.)

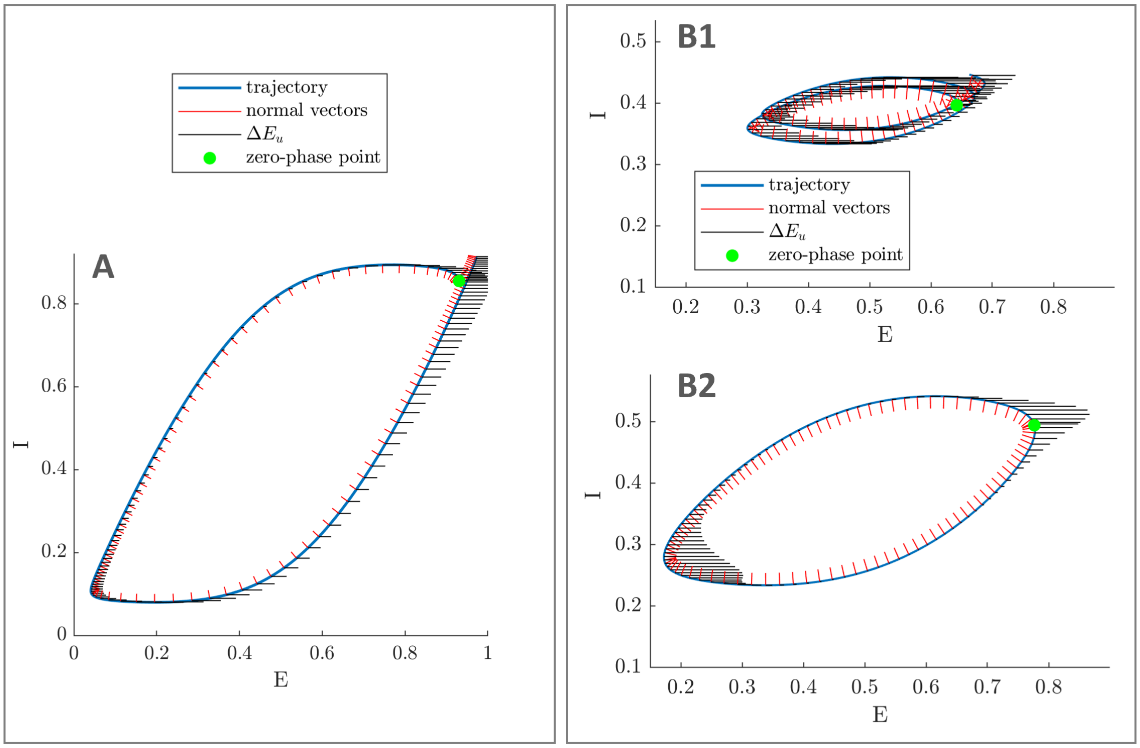


**Figure S.6: Obtaining amplitude response curves in the Wilson-Cowan model.** In both the strong theta (panel A), and pure gamma cases (panel B), the ARC is obtained along a trajectory of interest (blue line). The instantaneous change in the activity of the E population due to stimulation is shown in black at regular time intervals, and vectors normal to the trajectory are shown in red. The points taken as zero-phase references are shown in green. Normal vectors are represented with arbitrary scale for clarity, and $\Delta E_u$ is scaled by 8 times in A, and 20 times in B1-2. In all panels, $u = 0.3$. In the strong theta case (A), the trajectory considered is the on-stimulation gamma cycle, and $f_f = 41$ Hz. In the pure gamma case, the significant changes in dynamics for low and high amplitudes require to consider both the low-amplitude trajectory on-stimulation (B1), and the high-amplitude periodic trajectory (similar on- and off-stimulation, taken off-stimulation in B2). We take $f_f = 51$ Hz in B1-2.

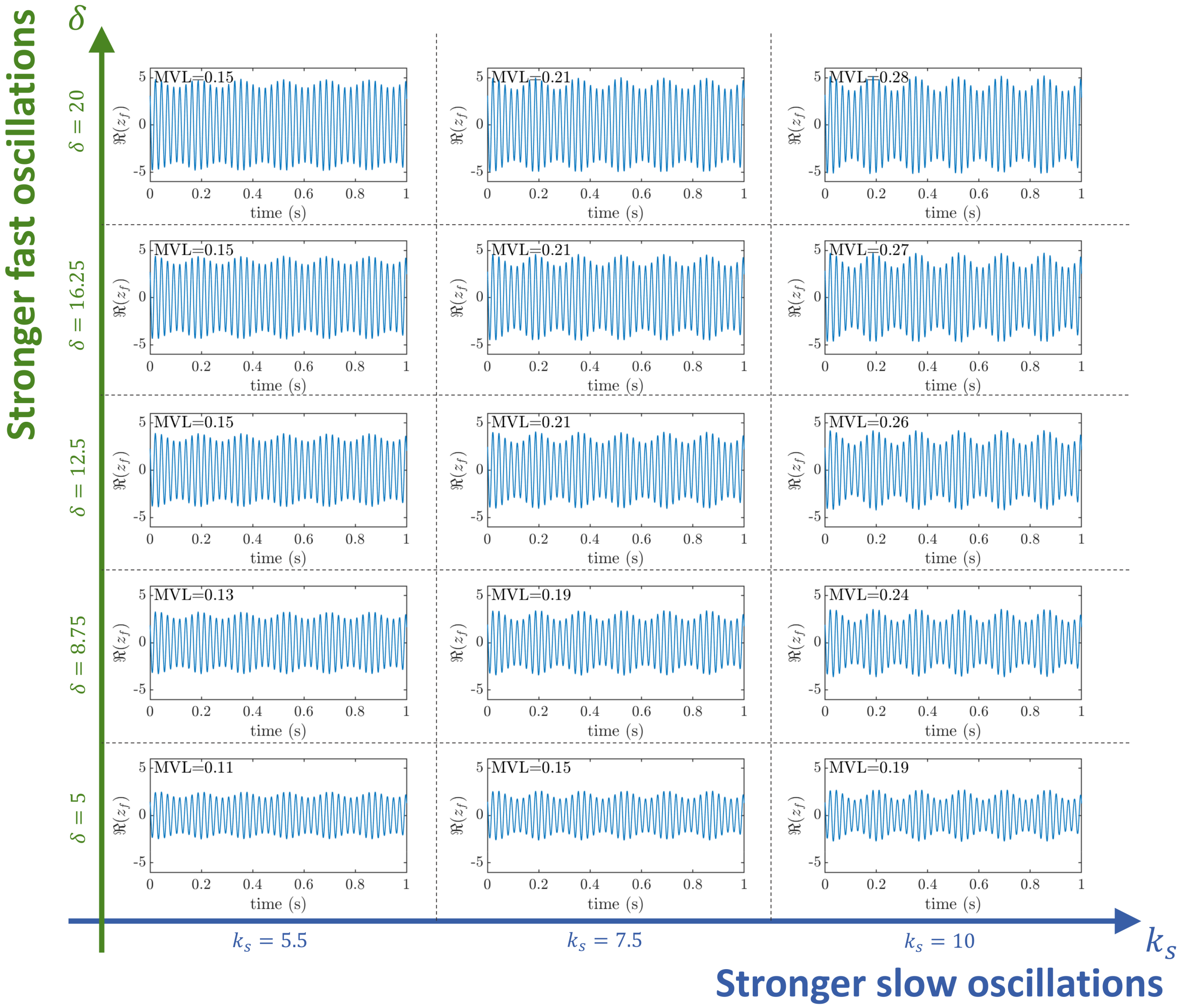


**Figure S.7: Off-stimulation output as a function of the strength of endogenous fast and slow oscillations in the Stuart-Landau model.** The strength of endogenous slow oscillations is controlled by model parameter $k_s$ (blue arrow), and the strength of endogenous fast oscillations by model parameter $\delta$ (green arrow). Model outputs (real part of the order parameter) are shown in the absence of stimulation for the values of $k_s$ and $\delta$ explored in figure 11. In all panels, the frequency of endogenous oscillations is $f_f = 42$ Hz, and $f_s = 6$ Hz.

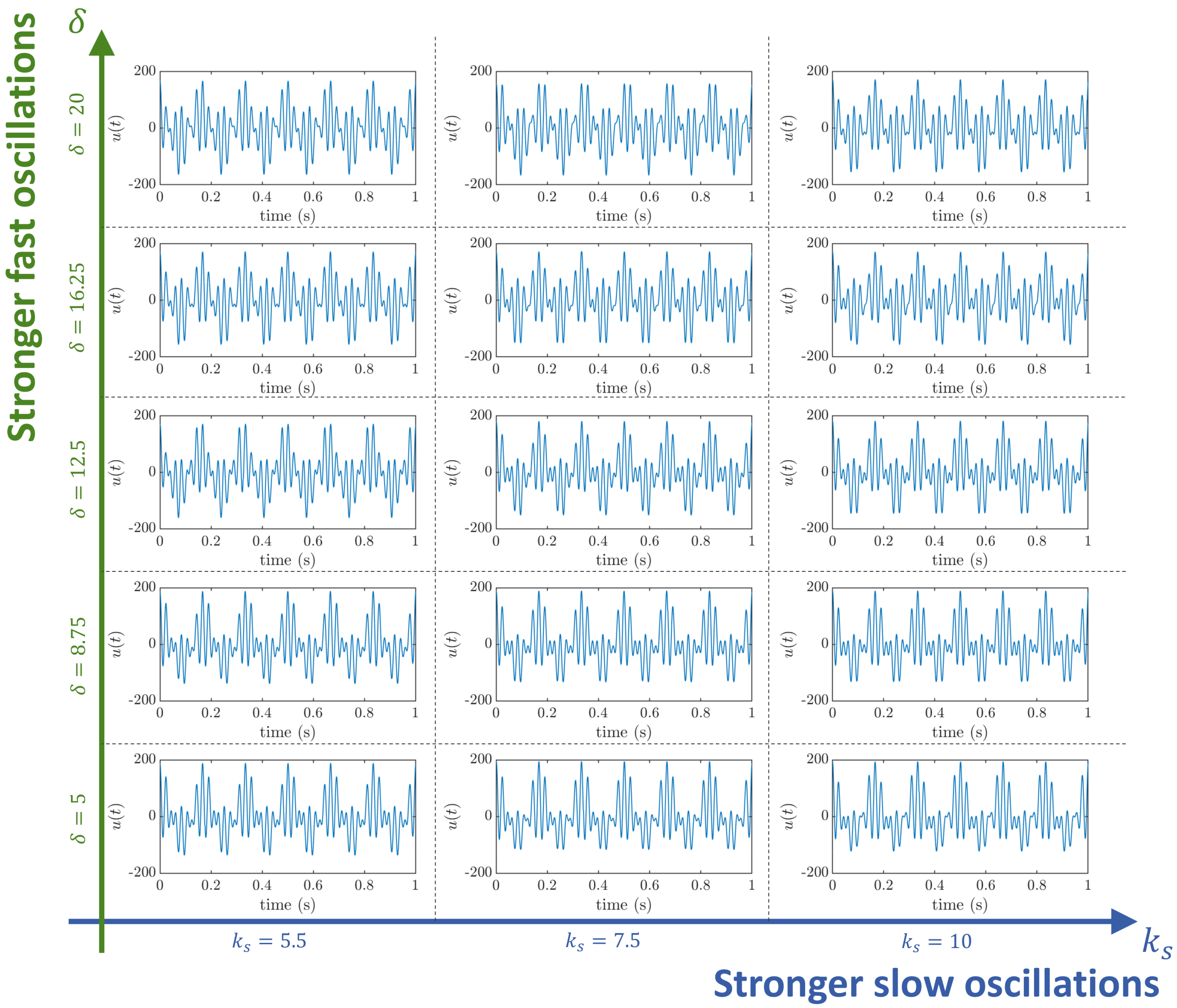


**Figure S.8: Optimal PAC-enhancing stimulation waveform as a function of the strength of endogenous fast and slow oscillations in the Stuart-Landau model.** The strength of endogenous slow oscillations is controlled by model parameter $k_s$ (blue arrow), and the strength of endogenous fast oscillations by model parameter $\delta$ (green arrow). The optimal PAC-enhancing stimulation waveforms (resulting from sweeping through $A_1$, $A_{r-1}$, $A_r$, and optimising the Fourier phases) are shown for the values of $k_s$ and $\delta$ explored in figure 11. In all panels, the total stimulation waveform energy is kept at $\Xi = 5000$, and the frequency of endogenous oscillations is $f_f = 42$ Hz, and $f_s = 6$ Hz. Stimulation is coupled to the fast population through $\text{ARC}(\theta_f) = 0.5 + \cos(\theta_f)$ and $\text{PRC}(\theta_f) = \sin(\theta_f)$.

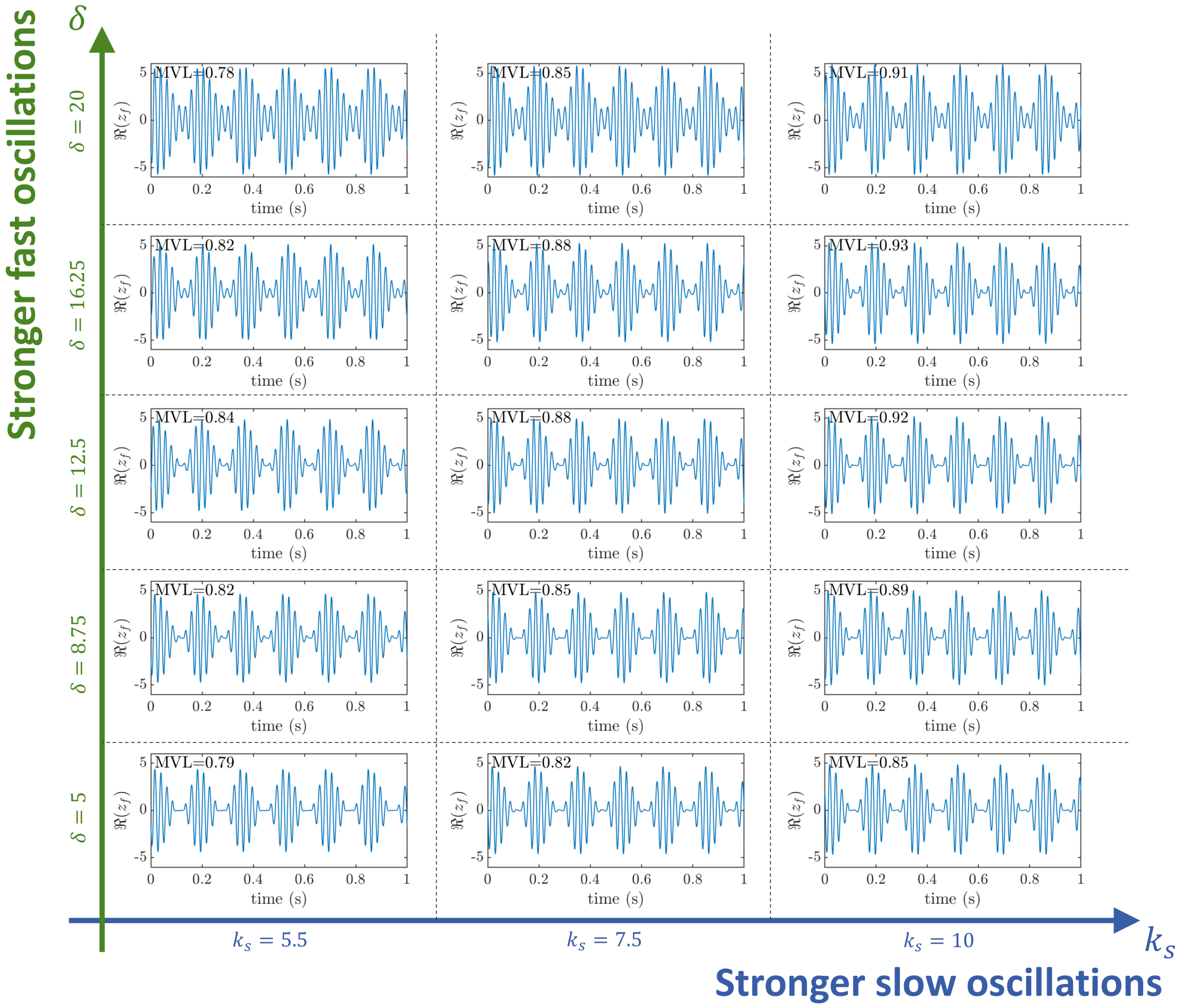


**Figure S.9: Model output with optimal PAC-enhancing stimulation waveform as a function of the strength of endogenous fast and slow oscillations in the Stuart-Landau model.** The strength of endogenous slow oscillations is controlled by model parameter $k_s$ (blue arrow), and the strength of endogenous fast oscillations by model parameter $\delta$ (green arrow). Model outputs (real part of the order parameter) under optimal PAC-enhancing stimulation waveforms (resulting from sweeping through $A_1$, $A_{r-1}$, $A_r$, and optimising the Fourier phases) are shown for the values of $k_s$ and $\delta$ explored in figure 11. In all panels, the total stimulation waveform energy is kept at $\Xi = 5000$, and the frequency of endogenous oscillations is $f_f = 42$ Hz, and $f_s = 6$ Hz. Stimulation is coupled to the fast population through $\text{ARC}(\theta_f) = 0.5 + \cos(\theta_f)$ and $\text{PRC}(\theta_f) = \sin(\theta_f)$.

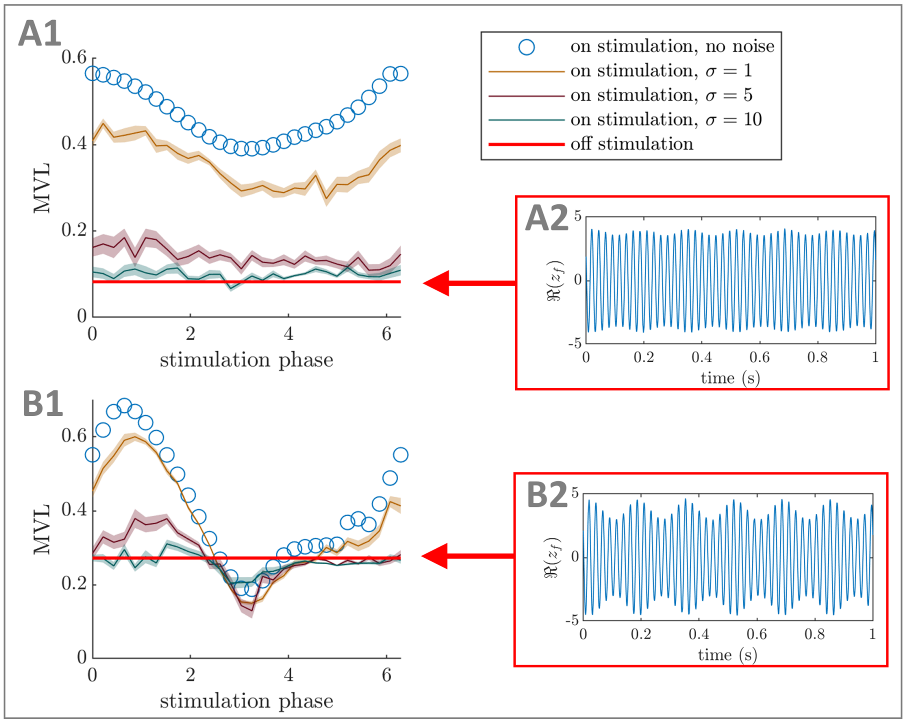


**Figure S.10: Fluctuations in the frequency of the fast population limit the efficacy of open-loop PAC modulation.** PAC-enhancing stimulation waveforms are provided at different phases of the slow rhythm, when the frequency of the fast population varies according to a Wiener process starting from $f_f$ with standard deviation $\sigma/(2\pi)$ Hz at the end of the simulation (50s). Panels A1 and B1 show the MVL as a function of stimulation phase of the slow rhythm in the absence of noise in blue (a stimulation phase of zero corresponds to the peaks of the stimulation waveform and the slow rhythm being aligned), and the off-stimulation MVL level in red. Other colors correspond to the mean MVL (averaged over 30 realisations) for varying levels of noise ($\sigma$, as indicated in the legend), with shaded areas corresponding to the standard error of the mean. The SL model is simulated with direct stimulation coupling and the stimulation waveform given in figure 4C. Panels A2 and B2 represent the corresponding off-stimulation model output in the absence of noise (real part of the order parameter). Parameters used are $f_f = 42$Hz, $f_s = 6$Hz, $\delta = 15$, and $\Xi = 5000$. Panels A1-2 correspond to $k_s = 3$, and panels B1-2 to $k_s = 10$.

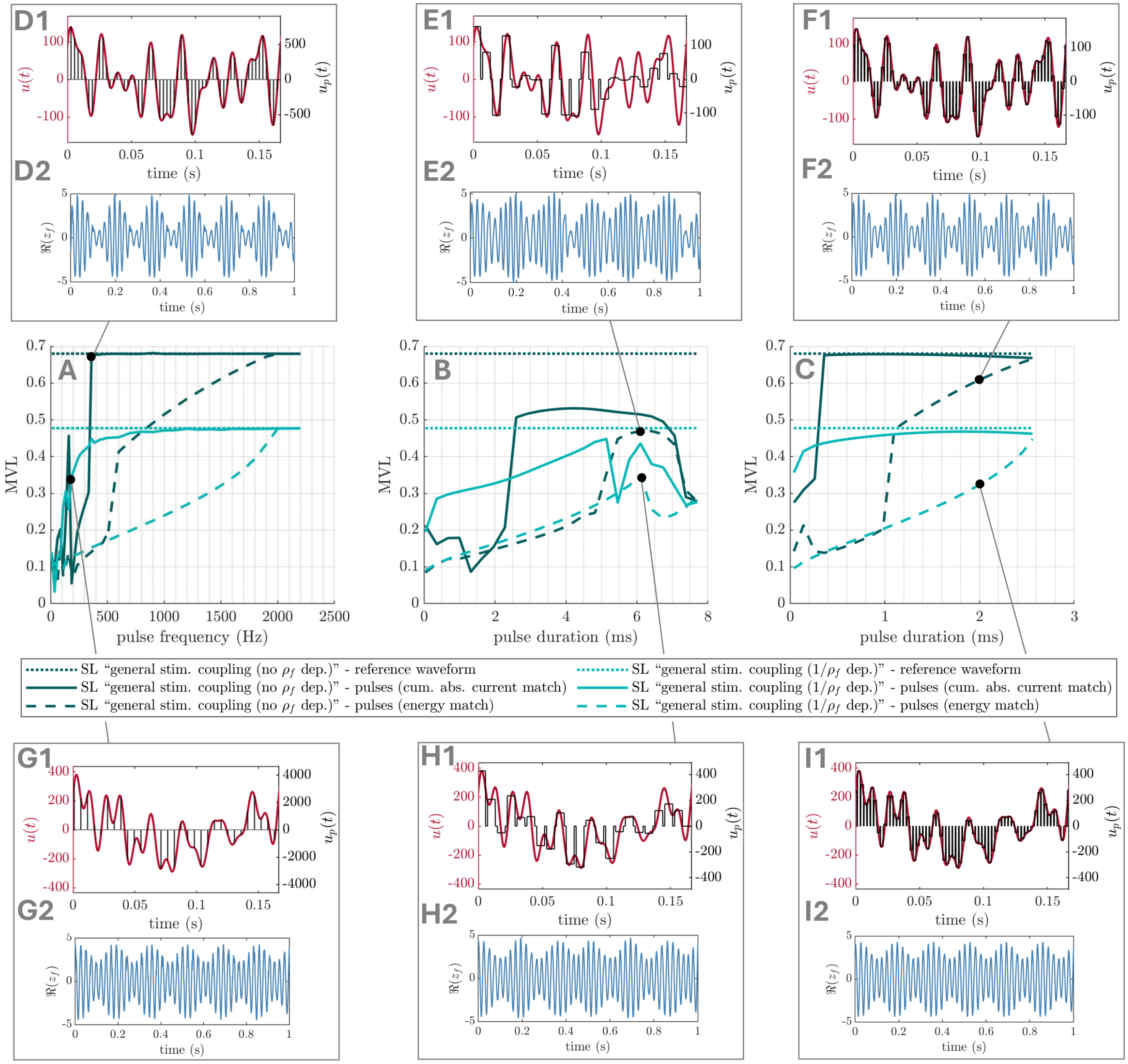


**Figure S.11: Optimal smooth waveforms can be approximated with pulses – general coupling cases in the SL model.** MVL is shown as a function of pulse frequency, for a pulse duration of 0.5ms (A), and as a function of pulse duration, for a pulse frequency of 130Hz (B) and 390Hz (C). In these panels, the case with no dependence on $\rho_f$ (parameters corresponding to figure 5B-D) is shown in dark green, and the case with $\propto 1/\rho_f$ dependence (parameters corresponding to figure S.4B-D) is shown in light green. Solid lines correspond to pulsatile waveforms obtained by matching the cumulative absolute intensity of the optimal smooth waveforms, while dashed lines correspond to pulsatile waveforms obtained by matching the energy of the optimal smooth waveforms. Dotted lines correspond to the smooth optimal waveforms. Panels D-I show the smooth optimal waveform in red and the pulsatile approximation in black (top), as well as the resulting activity of the fast population (bottom). Pulse frequencies/durations are as follow: 360Hz/0.5ms in D and G, 130Hz/6.1ms in E and H, 390Hz/2.0ms in F and I, and 185Hz/0.5ms in I.

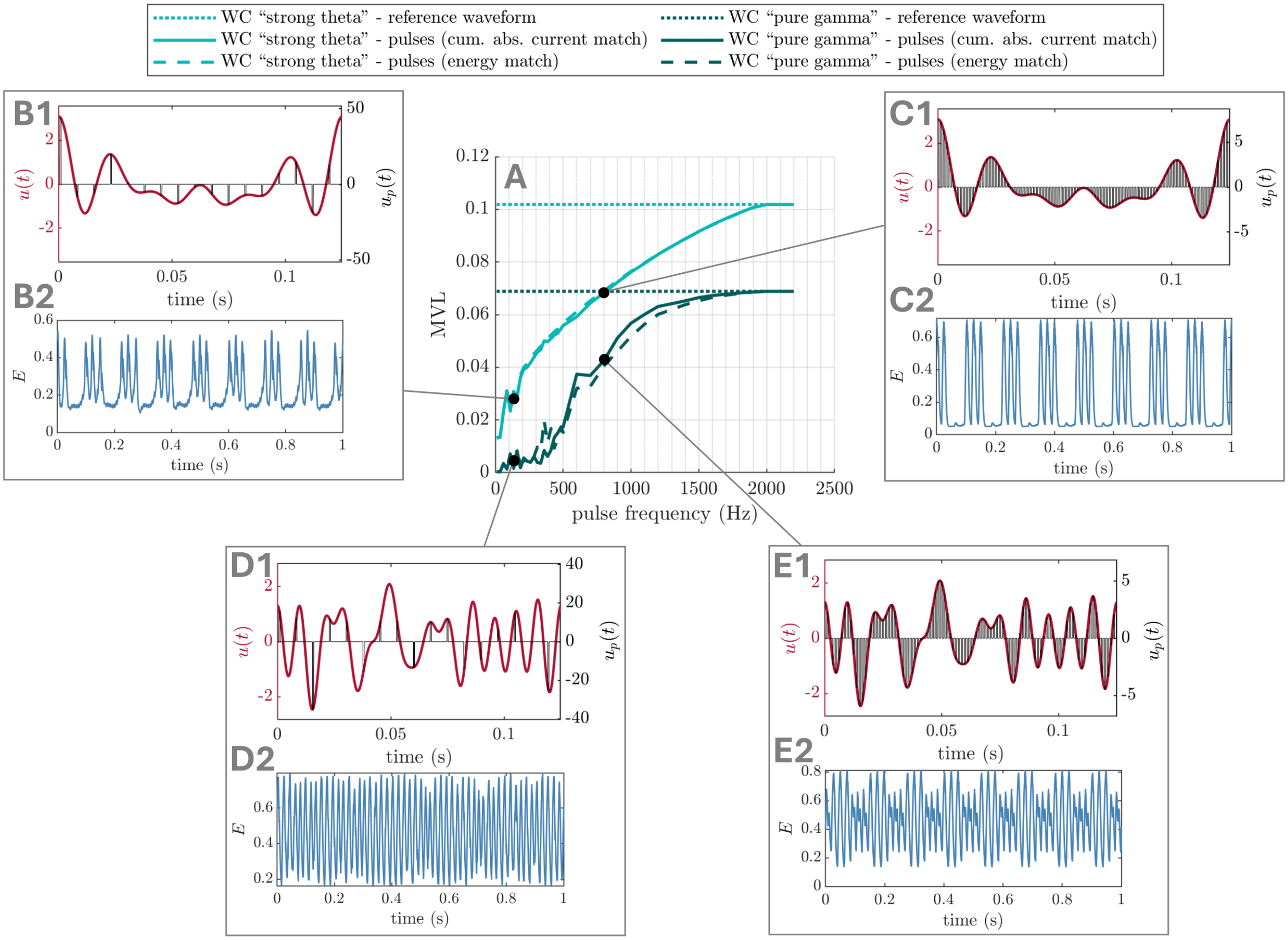


**Figure S.12: Optimal smooth waveforms can be approximated with pulses – WC model.** MVL is shown as a function of pulse frequency, for a pulse duration of 0.5ms. In these panels, the strong theta case (parameters corresponding to figure 7B-D) is shown in light green, and the pure gamma case (parameters corresponding to figure 8B-D) is shown in dark green. Solid lines correspond to pulsatile waveforms obtained by matching the cumulative absolute intensity of the optimal smooth waveforms, while dashed lines correspond to pulsatile waveforms obtained by matching the energy of the optimal smooth waveforms. Dotted lines correspond to the smooth optimal waveforms. Panels B-E show the smooth optimal waveform in red and the pulsatile approximation in black (top), as well as the resulting activity of the fast population (bottom). Pulse frequencies/durations are as follow: 135Hz/0.5ms in B and D, and 800Hz/0.5ms in C and E.
